# Structural Snapshots of B_12_-Dependent Methionine Synthase’s Catalytic Conformations

**DOI:** 10.1101/2024.08.29.610163

**Authors:** Johnny Mendoza, Kazuhiro Yamada, Carmen Castillo, Valerie Greenwood, Soumalya Pramanik, Rashi Salampuriya, Catherine A. Wilhelm, Markos Koutmos

**Author notes:** Johnny Mendoza and Kazuhiro Yamada contributed equally to this work. Institute for Protein Design, Seattle, WA, 98105. Indiana Biosciences Research Institute, Indianapolis, IN, 46202. Department of Cell and Developmental Biology, University of Michigan, Ann Arbor, MI, 48109.

## Abstract

Cobalamin (vitamin B_12_) and its derivatives play an essential role in biological methylation, with cobalamin-dependent methionine synthase (MS) serving as a canonical example. MS catalyzes multiple methyl transfers within a single, dynamic multi-domain architecture that has proven challenging to study, hampering efforts to elucidate its catalytic mechanism(s). Utilizing a thermostable MS homolog and non-native cobalamin cofactors, we have captured crystal structures of transient conformational states of MS, including those directly involved in folate demethylation and homocysteine methylation. These snapshots reveal the mechanistic significance of five-coordinate, His-off methylcobalamin in homocysteine methylation and highlight the crucial role of the folate-binding domain and interdomain linkers in orchestrating the intricate structural rearrangements required for catalysis. This expanded conformational ensemble, including the unexpected capture of novel ‘Cap-on’ conformations, underscores the remarkable plasticity of MS, exceeding previous estimations. Our findings provide crucial insights into the catalytic mechanism of MS, laying the foundation for harnessing cobalamin’s biocatalytic potential and elucidating how nature exploits protein dynamics to facilitate complex transformations.

## Main

Methionine synthase (MS) is a multi-modular enzyme capable of binding and activating three substrates (homocysteine, HCY; methyltetrahydrofolate, MTF/folate, FOL; and *S*-adenosylmethionine, SAM, Fig. 1a, c) to achieve three distinct methylations (Reaction I, II, and III, Fig. 1a). Reactivity is governed by substrate access to the cobalamin cofactor and its oxidation state (Fig. 1b). Each reaction is associated with the formation of a distinct catalytically competent ternary complex and an active site between the Cob domain and its respective substrate domain (Figs. 1d and Fig. 2)^1^. An additional module, the four helical bundle Cap domain, is thought to shield the cobalamin cofactor, preventing reactivity by gating access to its upper axial face (Figs. 1d and Fig. 2)^2^. The cobalamin cofactor’s oxidation state alternates between Co(III) and Co(I) states during the catalytic cycle (Reaction I and II, Fig. 1a) but the highly reactive Co(I) is oxidized to the catalytically inactive Co(II) once every 2000 turnovers^3,4^; MS can restore catalytic competency by entering the reactivation cycle, during which reductive methylation of Co(II) regenerates methylcobalamin, in the Co(III) state, allowing for re-entry to the catalytic cycle (Reaction III, Fig. 1a)^4,5^. While rich biochemical studies exist on factors that govern MS domain rearrangements, atomic level insights into the conformations associated with the catalytic cycle have eluded structural characterization^6,7,8,9^.

**Figure 1.**
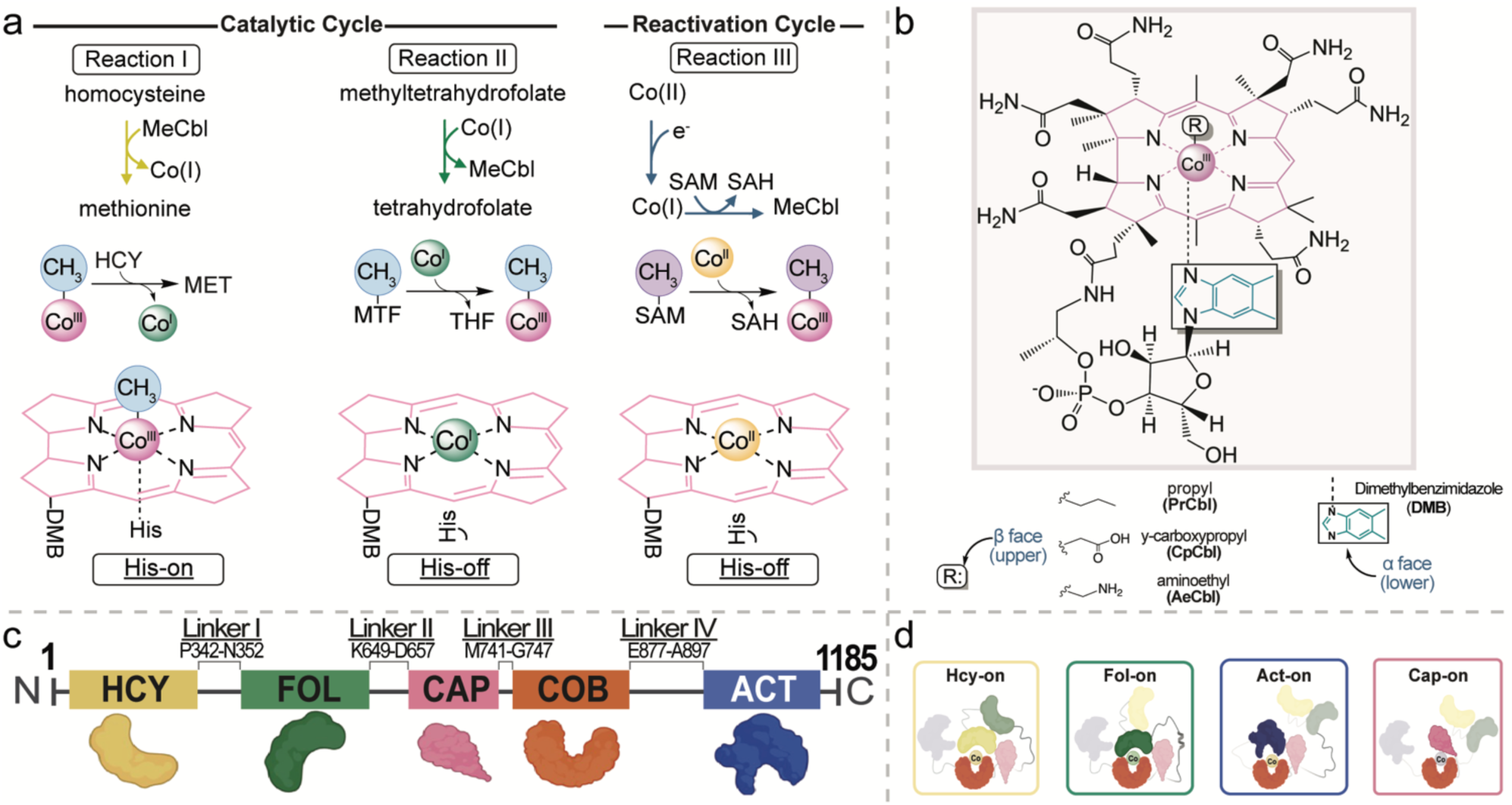
Methionine synthase harnesses protein dynamics and its cobalamin cofactor to dictate chemical outcome. **a** <u>Upper</u>: The three methylations performed by methionine synthase. The cobalamin cofactor cycles between redox states and coordination numbers (Reaction I: Co(III), six coordinate, pink; Reaction II: Co(I), four coordinate, green; Reaction III: Co(II), five coordinate, yellow). The primary catalytic cycle allows for the cobalamin cofactor to alternate between the Co(III) and Co(I) states for homocysteine methylation (Reaction I) and folate demethylation (Reaction II), respectively. Due to the microaerophilic conditions present in vivo, MS can undergo inactivation when Co(I) is oxidized to the inactive Co(II) state. During the reactivation cycle, the active state of the cofactor is restored via reductive methylation with SAM/AdoMet, regenerating the Co(III) state and allowing for re-entry to the catalytic cycle (Reaction III). The coordination number, redox state, and His ligation status are particular to each methylation. <u>Lower</u>: Directly below are the cobalamin cofactor oxidation states and coordination numbers associated with MS activity. The cobalamin cofactor is shown with the DMB base/lower-ligand ligated (Base-on) for simplicity. The cobalt center is color coded using the same color scheme as the upper panel of **a**. **b** The structure of the cobalamin cofactor, showcasing the corrin ring (pink), its upper axial (β face) ligand and its lower axial (α face) ligating nucleotide tail (dimethylbenzimidazole, DMB). Methionine synthase binds its native cobalamin cofactor, methylcob(III)alamin (R: CH_3_, methyl) in the base-off, His-on form. Synthetic cobalamin cofactors with alternate upper axial ligands incapable of catalysis can and have been used to modulate function via cofactor mimicry and subsequent inhibition and can capture distinct conformations (R: propyl, γ-carboxypropyl, aminoethyl). **c** Four compact substrate/cofactor binding (Hcy, Fol, Cob, and Act) and cofactor capping (Cap) domains are color-coded, with linkers displayed. **d** The minimum set of conformations MS must adopt during catalysis includes a Cap-on state that protects the cobalamin cofactor. Uncapping of the Cob domain allows for substrate domain access to the cobalamin cofactor, with three unique ternary complexes required for each methylation (Hcy-on, homocysteine methylation, Reaction I; Fol-on, folate demethylation, Reaction II; Act-on, cobalamin reductive reactivation, Reaction III).

**Figure 2.**
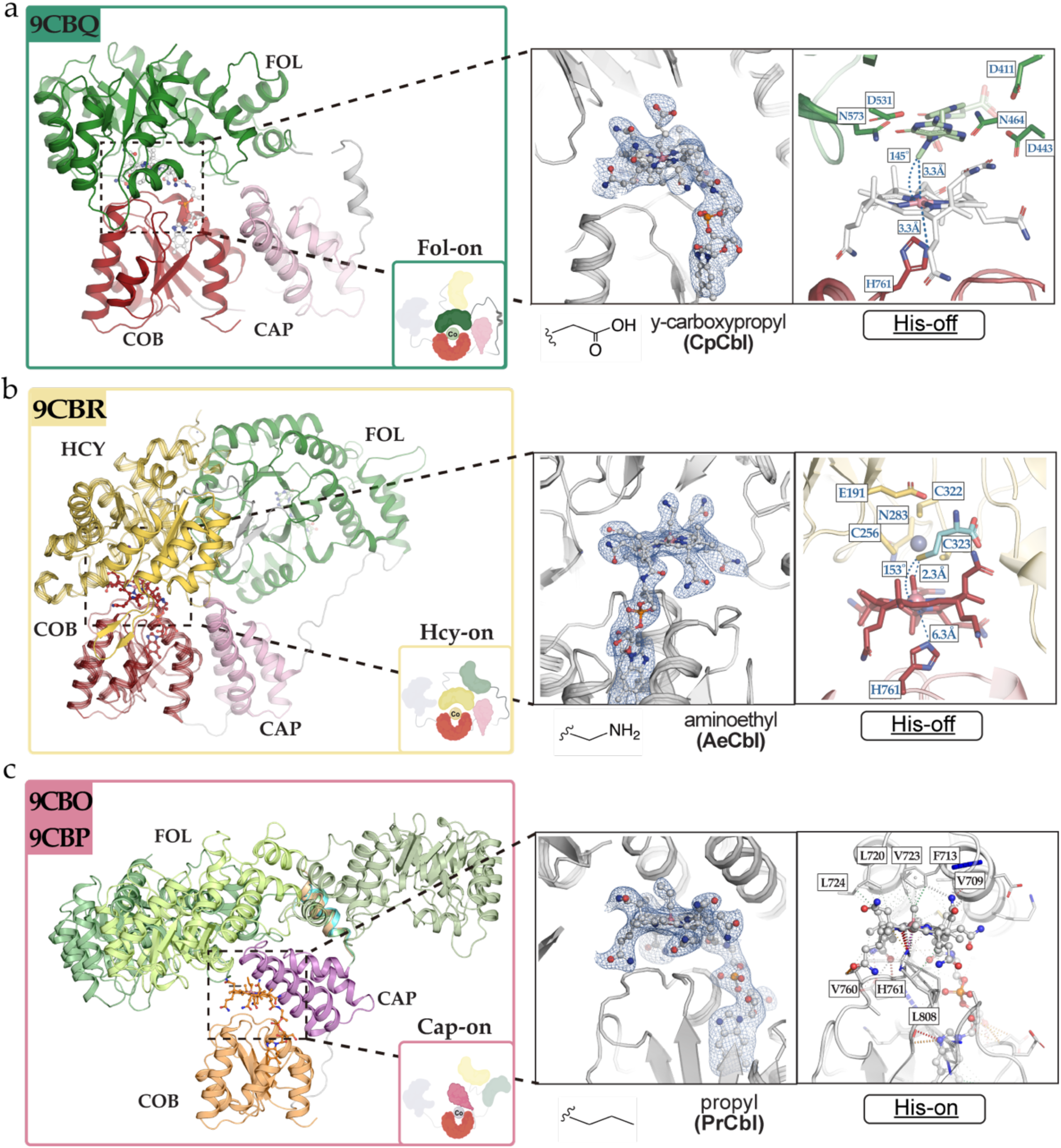
Methionine synthase structures captured via crystallography reveal a more expansive conformational ensemble and the capture of two elusive catalytic conformations. Four compact substrate/cofactor binding (Hcy, Fol, Cob, and Act) and cofactor capping (Cap) domains are color-coded, with linkers displayed. The minimum set of conformations MS must adopt during catalysis includes a Cap-on state that protects the cobalamin cofactor. Uncapping of the Cob domain (Cap-off) allows for substrate domain access to the cobalamin cofactor, with three unique ternary complexes required for each methylation: Fol-on, folate demethylation, Reaction II (shown in **a**), Hcy-on, homocysteine methylation, Reaction I (shown in **b**), Act-on, cobalamin reductive reactivation, Reaction III**). c** Experimentally determined Cap-On structures enable MS ‘resetting’, while Cap-Off structures facilitate ternary complex formation between the Cob and substrate binding domains as part of the catalytic cycle (Reactions I, II). Similarly, Cap-on to Cap-off transitions enable chemistry as a part of the reactivation cycle (Reaction III). The Cap-on states protect the cobalamin cofactor, and the uncapping transition results in a ∼25 Å displacement of the Cap domain to a predetermined position nestled to the side of the Cob domain, allowing for substrate-binding domain access and chemistry by enabling access to the cobalamin cofactor. Note the use of non-native, non-reactive cobalamin cofactors used to capture transient Cap-off states: γ-carboxypropylcobalamin (cpCbl) for the Fol-on state (9CBQ), aminoethylcobalamin (aeCbl) for the Hcy-on state (9CBR), and propylcobalamin (prCbl) for the capture of the pre-Hcy-on state (9CBP). The use of the native methylcobalamin cofactor (MeCbl) resulted in the capture of two distinct Cap-on conformations in one crystal structure (9CBO), one designated as the pre-Fol-on state and the other awaiting further designation (tentatively ‘resting’, 9CBO). Cobalamin cofactors are shown (gray) and their corresponding electron density (2F_o_-F_c_) contoured at 1.5 σ are shown in blue (left inset). The His-ligation status (His761) of each state along with residues present in the upper axial portion of the cofactor are shown (right inset).

Biochemical studies have faced limitations imposed by the inherent instability of traditional MS homologs (e.g. *E. coli, H. sapiens*), primarily the inability to purify the apoenzymes, their susceptibility to proteolytic degradation, and the limited capability to conduct mutagenesis studies. Historically, a ‘divide-and-conquer’ strategy, where the more stable excised domains are used in lieu of the full-length enzyme, has been employed to structurally characterize MS in a piecemeal approach. Several studies have also reconstituted MS activity *in trans* using these excised domains, the N-terminal half (containing the Hcy and Fol domains) and the C-terminal half (containing the Cap, Cob, and Act domains), or exogenously provided cobalamin^10,11,12,13^.

Rigorous biochemical studies that leveraged the use of truncated constructs as minimal MS models proposed a working model of four conformations representing the minimal set of states necessary for MS to sample and perform catalysis (Fig. 1d).^8^ Methylcobalamin serves as a key focal point, acting as a bridge between the catalytic and reactivation cycles. Cob(I)alamin generated by homocysteine methylation reacts solely with methyltetrahydrofolate, whereas cob(I)alamin generated by reduction of cob(II)alamin reacts solely with *S*-adenosylmethionine^8^. This methylation of cob(I)alamin by specific methyl sources demonstrates that the MS conformational ensemble sequesters the supernucleophilic cob(I)alamin between each cycle, preventing futile cycling by limiting substrate access to the highly reactive cofactor.

Previously, we demonstrated the versatility of a thermophilic MS homolog (*t*MS) from *Thermus thermophilus* that displayed robust stability, completely bypassing the systematic challenges associated with MS purification: notably, to our knowledge, *t*MS is the only example of an MS construct that can be purified and obtained in its *apo*-form, allowing for holoenzyme formation at will. In addition, it displayed remarkable robustness to biochemical manipulation, allowing for exhaustive functional mutagenesis studies, and facile reconstitution with non-native cobalamin cofactors to form novel holoenzymes (Fig. 1b)^14^. This model system shares the same multi-modular architecture, comparable functional properties to previously characterized MS homologs, and allowed for the determination of the first full-length structure of MS (8SSC) captured via crystallography and the observation of holoenzyme formation *in crystallo* (8SSD, 8SSE)^14^. The ability to purify excised domain constructs, coupled with the ability to purify all mutants generated, indicate *t*MS is highly amenable to biochemical studies and structural-functional characterization, thereby allowing us to begin interrogating MS mechanistically using a combination of rationally chosen non-native cobalamins and constructs^14^.

Here, we leverage our *t*MS model to capture five new crystal structures of MS in action, three distinct Cap-on conformations and two catalytically competent conformations, that provide insights into the structural motifs used to guide and gate the large conformational rearrangements required for MS catalysis. We show that the MS conformational ensemble is more expansive than previously appreciated, adopting additional states prior to sampling catalytic conformations.^15,16^ These additional conformations highlight the helix-loop restructuring of Fol:Cap linker (Linker II, Figs. 2c and Supplementary Fig. 1) during conformation transitions. The structures show that the “uncapping” transition (Cap-on to Cap-off) required for active site formation (Cap:Cob linker, Linker III, Supplementary Figs. 2 and 3), is predetermined, adopting the same placement relative to the Cob domain in all observed Cap-off structures. The two catalytic conformations (Fol-on, Reaction II, Fig. 2a; Hcy-on, Reaction I, Fig. 2b) provide the first structural blueprints of a B_12_-dependent methyltransferase active site, one required for folate demethylation and the other for homocysteine methylation. Our biochemical data highlight the role the Fol domain plays in steering the conformational ensemble towards catalytic states, while also demonstrating that the cobalamin cofactor adopts a His-off, five-coordinate Co(III) state during catalysis. Notably, we demonstrate the importance of the ligating His residue in signaling cofactor status (redox state, coordination number) and guiding the conformational ensemble, positing that His-on ligation gates the ensemble into Cap-on states (pre-catalytic conformations), enabling MS to ‘reset’ and interconvert between these states, while His-off ligation allows for cobalamin flexibility and substrate-bound domain access after uncapping (Cap-off, catalytic/reactive). We propose a revised mechanistic and structural model for MS, one that emphasizes the role of a linker (Fol:Cap, Linker II), His-off ligation, and guided domain uncapping in gating and thereafter guiding rearrangements that allow for cofactor access required for catalysis.

**Figure 3.**
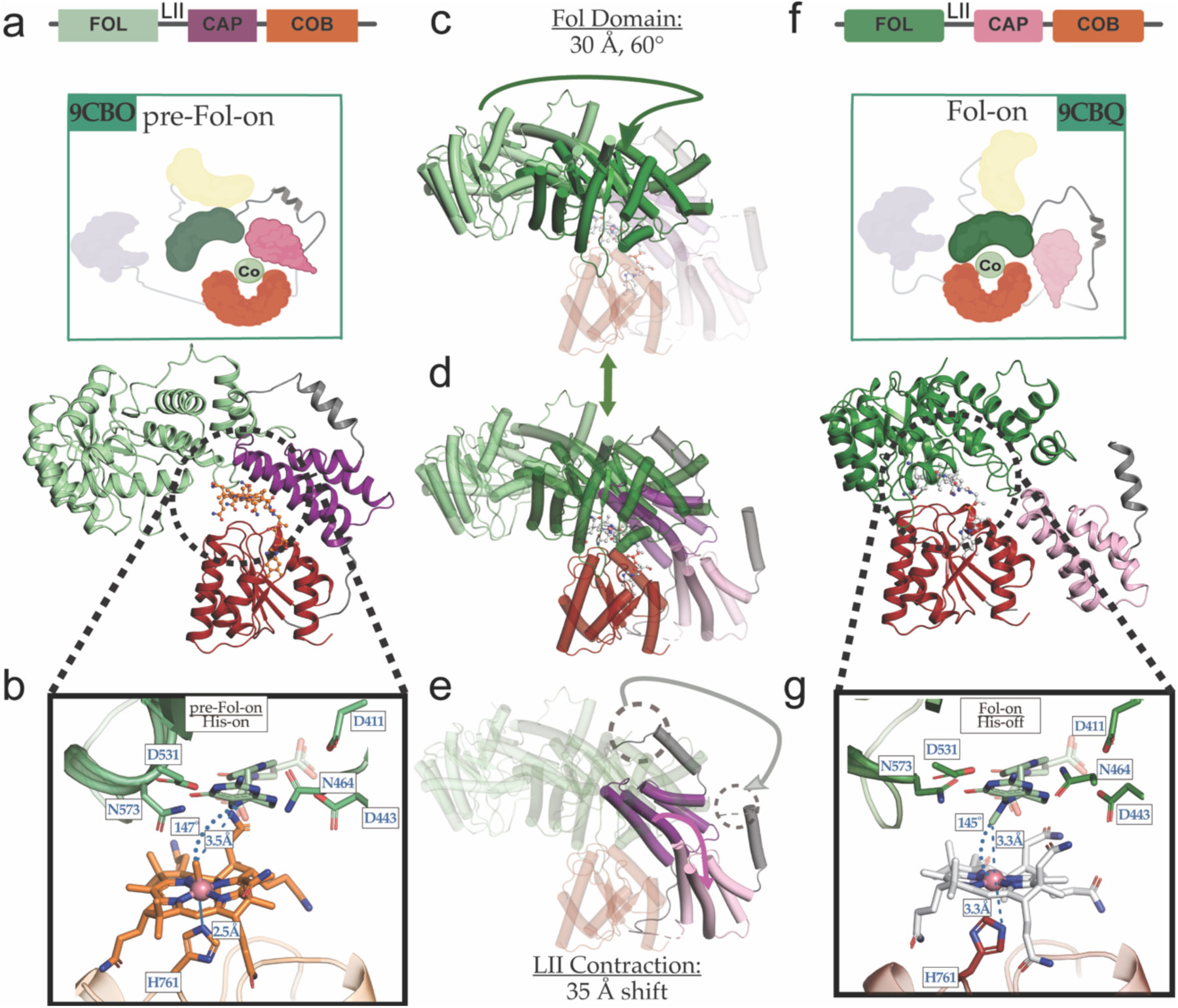
Structural comparison of pre-catalytic states and catalytic states (pre-Folate-on and Folate-on). **a-b** Structure of *t*MS^FolCapCob^ captured in the pre-Fol-on conformation colored by domain (Fol domain, pale-green; Cap, dark purple; Cob, orange) with the Fol:Cap linker shown in grey. **f-g** Structure of *t*MS^FolCapCobD762G^ captured in a catalytic conformation (Fol, forest green; Cob, dark red; Cap, light pink) with the Fol:Cap linker shown in grey. His761 distance between pre-catalytic and catalytic states in *t*MS^FolCapCob^ are shown in **b** and **g**, respectively. The interactions between His761-Nε2 and the Co center are shown in dark orange (2.5 Å) and magenta (3.3 Å) dotted lines for pre-Fol-on and Fol-on, respectively. Superposition of those structures are illustrated in **c**. The Cob domain was used as the reference for the structure alignment. The cobalamin cofactor must traverse 25 Å to form the Fol-on ternary complex. **d** Superposition using the Cob domain as a reference of the Fol domain *t*MS^FolCapCob^ structures using the same coloring scheme as **a** and **f**. Contraction of the Fol:Cap linker causes a 60° rotation coupled with a 30 Å translation of the Fol domain (**c**), while simultaneously uncapping the cobalamin cofactor bound in the Cob domain. **e** The Fol-Cap linker (Linker II) contracts by 35 Å, the movement of which seems to be coupled to the Cap-on to Cap-off transition and the Fol domain rearrangement to interact with the cobalamin domain, forming the ternary complex involved in methyltetrahydrofolate demethylation (Reaction II).

### Cap-on conformations expand Methionine Synthase’s conformational ensemble

The ability to purify *t*MS as an apoenzyme, load non-native cobalamins, and generate constructs that were previously unattainable, provided an avenue to strategically limit the structural conformations adopted by MS via tactical truncations. By using constructs lacking the reactivation domain (Act), one can reduce the conformational space MS can sample, as the reactivation conformation (Act-on) cannot be formed. By using the FolCapCob tridomain construct, we envisioned capturing the ternary catalytic folate demethylation complex (Fol-on). Similarly, we employed the HcyFolCapCob tetradomain construct to capture the ternary catalytic homocysteine methylation complex (Hcy-on).

Accordingly, *t*MS^FolCapCob•D759A^ was crystallized bound to its native cofactor, methylcobalamin (MeCbl), and its structure was solved to 3.34 Å (Figs. 2c and 3a, Supplementary Fig. 1, and Supplementary Table 1, 9CBO) as well as *t*MS^HcyFolCapCob•mut1^ bound to propylcobalamin (prCbl), chosen for its precedence sterically disfavoring Cap-on states^20^. This structure was solved to 2.45 Å (Figs. 2c and 4b, and Supplementary Table 1, 9CBP). The former captured two novel Cap-on conformations in one crystal, distinct from the only other reported Cap-on conformation of a muti-domain MS (HcyFolCapCob, 8G3H). The two novel Cap-on conformations differ in the orientation of the Fol domain and folate binding site relative to the Cob domain (Supplementary Figs. 1 and 2, 9CBO), along with the length of the linker connecting the Fol and Cap domains (Linker II, Supplementary Fig. 1).

**Figure 4.**
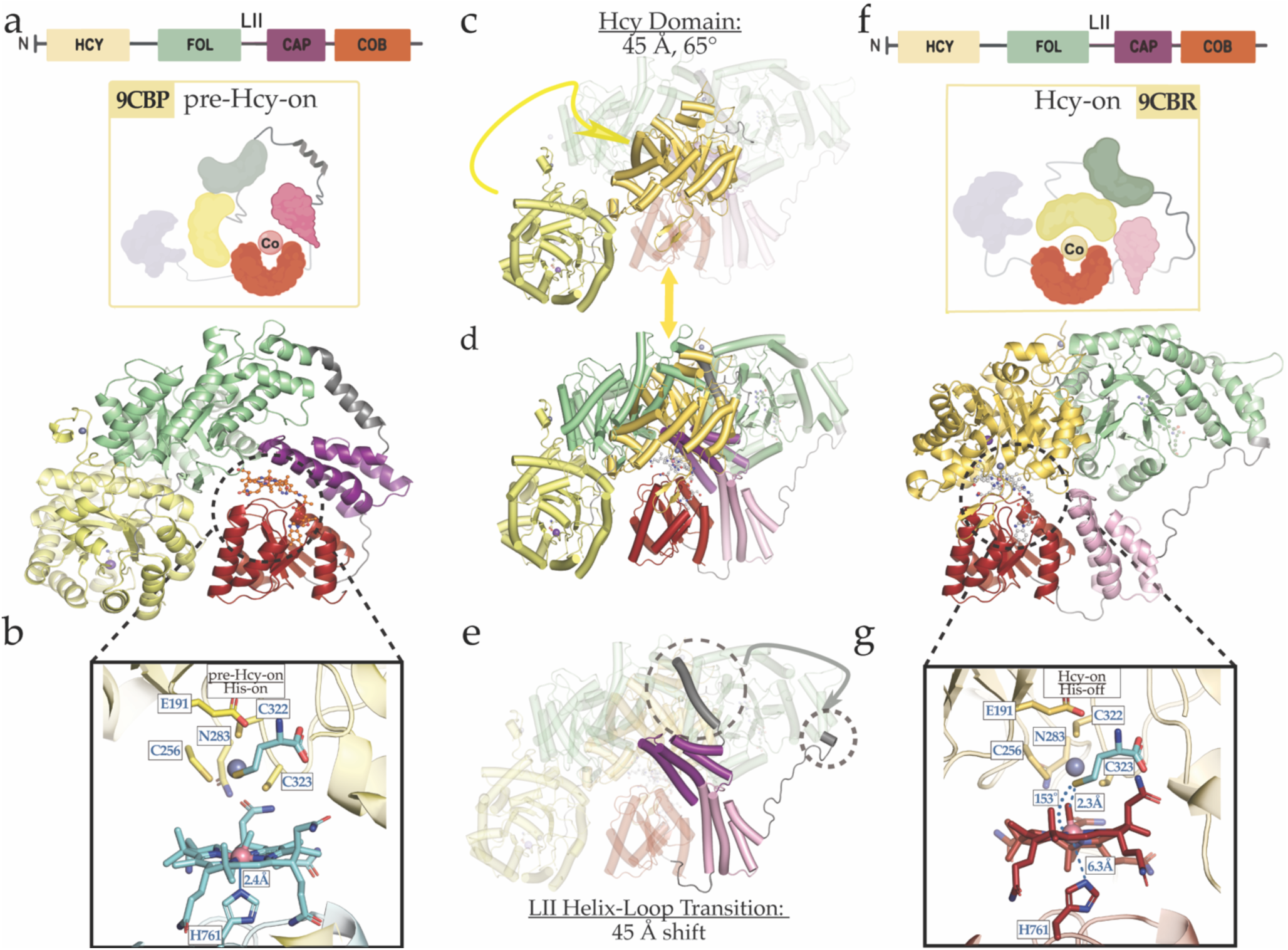
Structural comparison of pre-catalytic states and catalytic states (pre-Homocysteine-on and Homocysteine-on). **a-b** Structure of *t*MS^HcyFolCapCobmut1^ captured in in the pre-Hcy-on conformation colored by domain (Hcy domain, pale yellow; Fol domain, pale-green; Cap, dark purple; Cob, orange) with the Fol:Cap linker shown in grey. **f-g** Structure of *t*MS^HcyFolCapCobmut2^ captured in a catalytic conformation (Hcy, yellow orange, Fol, forest green; Cob, dark red; Cap, light pink) with the Fol:Cap linker shown in grey. **b, g** His761-Nε2 distance between pre-catalytic (light cyan, pre-Hcy-on) and catalytic states in *t*MS^HcyFolCapCob^ (brick red, Hcy-on). The cobalamin cofactor is displaced 3.8 Å laterally and is bound His-off in the catalytic conformation (2.32 Å versus 6.32 Å, ∼Δ4 Å). Superposition of those structures are illustrated in **c**, using the Cob domain as a reference. **d-e** Contraction of the Fol:Cap linker causes a dramatic 45 Å repositioning of the Cob domain with the Hcy domain to form the Hcy-on ternary complex. The Fol:Cap linker undergoes a helix-loop transition. The near complete melting of the helix enables the Hcy:Fol domains to move as rigid bodies after uncapping, repositioning the Hcy binding site and domain directly above the Cob domain and its bound cobalamin cofactor.

The three observed Cap-on states bind cobalamin in the His-on, base-off manner, and they differ in the proximity and orientation of the two domains (Hcy for 9CBP, Fol for 9CBO) and their substrate-binding sites relative to the Cob domain and cobalamin. This reveals that the HcyFol domains, which have previously been shown to move as a tandem rigid body, can undergo dynamic reorientations *en route*/prior to the cofactor ‘uncapping’ required for catalysis (Cap-on to Cap-off). Capturing these three Cap-on conformations, indicates that the premise that a single Cap-on state represents a “resting” state is an oversimplification. Indeed, while the authors termed their structure (8G3H) as *the* “resting” state, we argue that it represents one of *at least* three ‘resting’ conformations. The capture of three distinct Cap-on states was wholly unexpected and indicates that the structural space MS samples is more complex than previously thought, encompassing more than the predicted minimal set of four conformations (Fig. 1d)^1,7,8,9,16,18^.

The defining feature of the Cap-on conformations is the structured Fol:Cap linker (Linker II), which was found as an unstructured loop in the full-length *t*MS structure (8SSC)^14^. In the three Cap-on states presented here (Supplementary Figs. 1 and 2), Linker II is found as a flexible helix. The length of this linker varies in the different structures, and with it, the orientation of the adjacent Fol domain. This partial melting of the helix allows Linker II to act as a hinge, as the domains themselves maintain their topology and move as rigid bodies (Supplementary Fig. 2). The hinge contracts and rotates the Fol domain 120° to orient the folate binding site toward the Cap:Cob domain, as opposed to its orientation in the Cap-on state recapitulated and previously captured in 83GH, where the folate binding site is facing away (9CBP, Supplementary Fig. 2, RMSD = 1.812 Å over 5236 to 5236 atoms).

The use of truncated constructs with methylcobalamin and even propylcobalamin did not yield the desired Cap-off states, despite literature precedence for the strategic use of propylcobalamin (prCbl) to disfavor Cap-on states due to the bulkier nature of the upper axial ligand^13,14^. Coupled with the differing status of Linker II (e.g. its helical versus loop transition) and given that the use of alkylcobalamins led to Cap-on states, an attempt was made to leverage other cobalamin analogs while simultaneously introducing mutations to steer the ensemble towards Cap-off states.

### Visualizing methyl-transfer in *t*MS

#### Folate demethylation (Fol-on) conformation

We next sought to exploit the robustness of our *t*MS model to protein engineering by introducing mutations designed to steer the conformational ensemble away from the His-on, Cap-on state and explored the use of bulkier, non-alkyl upper axial ligands for the cobalamin cofactor that would further destabilize the Cap-on conformations ^9,17,18,19,20^. A D762G mutation was introduced, based on the finding that the Cap-on structures contained a stabilizing interaction between that residue and one of the propionamide arms of cobalamin. In addition, the Cap domain provided several hydrophobic interactions that stabilized the alkyl upper ligands (F713, L720, V723) by providing a hydrophobic pocket (Fig. 2c, right inset). To that end, we selected γ-carboxypropyl cobalamin (cpCbl) (Fig. 1b), a synthetic cobalamin that does not support catalysis and carries a negatively charged upper ligand to destabilize the hydrophobic pocket provided by the Cap domain; using this analog, we were able to crystallize *t*MS^FolCapCob•D762G^, solving the structure to 2.87 Å (Figs. 2a and 3f, and Supplementary Table 1, 9CBQ).

In the captured conformation (Fol-on), the Fol-domain is positioned over the Cob-domain, placing the MTF-binding site directly above cobalamin cofactor (Fig. 3g). His761 is found to coordinate one of the propionamide arms of the cofactor through its imidazole side chain 3.3 Å from Co, with His761-Nε2 facing away from the cobalt center (Fig. 3g, inset). This results in an increased distance between the ligating His761 residue and the cobalamin cofactor (∼Δ0.8 Å as compared to the Cap-on, pre-Fol-on conformation, Fig. 3b). This His-on to His-off like transition results in repositioning of the cobalamin cofactor: the Co center is shifted ∼1.5 Å laterally with an associated downward tilt of ∼10° in the A ring (corrin helicity change from +4.3 to −5.3, ∼Δ10°) The corrin ring, which experiences significant distortion, is more planar in the captured Fol-on state (Supplementary Fig. 4; inter-planar angle *φ* of ∼10.8° versus ∼5.3°). The increased distance and subtly altered orientation of the ligating His761 (2.3-2.5 Å from the Co center in Cap-on versus 3.3 Å in the Cap-off, Fol-on state, His761-Nε2) results in a relieving of downwards “doming” of the corrin towards the ligating His.

The Cap domain is displaced ∼25 Å from the Cap-on state conformations^15,16^ in a manner virtually identical in form and position to previously captured Cap-off structures, lying to the periphery of the Cob domain (8SSC, 8SSD, 8SSE, Supplementary Fig. 3)^14^. This suggests that the position of the displaced Cap domain is programmed, rather than random. The dramatic rearrangement of the cobalamin cofactor and the domain rearrangements (Supplementary Fig. 5a), coupled with the restructuring of Linker II in lieu of interdomain interactions, validates our previously proposed hypothesis that the flexibility of the linkers must be key in facilitating these domain movements^14^.

Starting from a Cap-on state (9CBO), Linker II must undergo a loop-helix transition to form the Fol-on ternary catalytic complex. In this case, the N-terminal portion of the linker is disordered (4 residues unmodeled); again, this small change results in a corresponding 34.5 Å shift of the linker and 60° rotation of the Fol domain relative to the Cob domain (Fig. 3c, d, and e, Supplementary Fig. 2). Combined with the Cap-off transition, this shift allows for the cobalamin cofactor to be completely buried by the new Fol:Cob interface (buried surface area of ∼1310 Å^2^, 89.4% total versus ∼1156 Å^2^, 74.3% total in the assigned pre-Fol-on Cap-on state, Supplementary Fig. 3). The removal of the hydrophobic pocket formed by the Cap domain allows for the formation of new stabilizing hydrogen bonds with the Fol domain that fully anchor the propionamide arms of cobalamin including some involving an intricate water-mediated network (Supplementary Fig. 6).

Manual inspection of the electron density maps (2Fo-Fc, Supplementary Fig. 7c) and composite omit electron density maps (Fo-Fc, Supplementary Fig. 8a) showed unmodelled electron density in the upper axial position (β-face) of cobalamin. As such, the upper axial ligand of cpCbl (butanoic acid, γ-carboxypropyl, Ligand ID BUA) was modelled. This ligand was found to form a salt bridge with the strictly conserved Asp531 (Supplementary Figs. 20 and 21)^21^ and coordinate an active site water (Supplementary Fig. 8a). Mutagenesis of this corresponding residue (Asp522 in *E. coli* MS, Asp522Asn mutant) completely abolished methyltetrahydrofolate binding *and* activity^12^. Asp531 or other functionally equivalent Asp residues have been shown to be essential to folate binding across several folate-activating enzymes^22^, forming a bidentate interaction with N3 of the pterin ring and the exocyclic amine at position 2 (NA2, Supplementary Fig. 6), while the active site water provides a bridging interaction between Asp531 and O4 of the pterin ring, providing a connection to an extended hydrogen-bonded network (Supplementary Figs. 6, 22a)^21^. Though purification of *t*MS constructs containing the Fol domain routinely result in copurification of methyltetrahydrofolate^21^, no folate signature was observed via UV-Vis or in the solved crystal structure.

##### Capture of the Elusive Fol-on Conformation reveals how methyltetrahydrofolate demethylation can occur

To determine if the orientation captured in the Fol-on structure could support catalysis, we modeled MTF into the active site and found that the N5-methyl of MTF is positioned directly above the cofactor, ∼3.2 Å from Co-center (Fig. 3g, ∼149° angle between N5 bearing the C11 methyl group and the cobalt center, Supplementary Fig. 4b)^21^, in a linear arrangement consistent with that required for the S_N_2 nucleophilic displacement mechanism favored for cobalamin-dependent methyltransferases^21,22^. Notably, the modelled MTF ligand, particularly its pterin ring, interacts with Asp531 and the aforementioned active site water with which the upper axial ligand of cpCbl is bound (O1 and O2 of γ-carboxypropyl vs. N3 and O4 of MTF, respectively). In addition, the C11 methyl group of MTF is found in a similar position to the C1 group of γ-carboxypropyl that coordinates the cobalt center. Thus, binding of cpCbl directly competes with the ‘pterin hook’ required for folate binding, while also coordinating the active site water postulated to allow for activation of the tertiary amine methyl donor in MTF (protonation of N5 via keto-enol tautomerization of O4, Supplementary Fig. 6).

Both the arrangement and proximity of the reacting centers indicate the captured state represents a catalytically competent structure, or a ‘right-after-catalysis’ state. The cobalamin cofactor, cpCbl, is found bound in an intermediary His-off state, where His761 (Nε2) is facing away from the cobalt center, ∼3.3 Å away from the corresponding Cap-on, His-on state (pre-Fol-on). In total, this structure represents the Fol-on (Fol:Cob) state, adopted for the methyl transfer from MTF to Cob(I). As the first catalytic structure visualized for any corrinoid folate enzyme, this structure gives unique insight into the required orientation of the Fol and Cob domains and informs how the initial methyl transfer to form methylcobalamin can proceed. Encouraged by the ability to capture transient conformations using crystallography, we next sought to structurally interrogate the homocysteine methyltransfer reaction using a similar approach (Reaction I, Fig. 1).

#### Homocysteine methylation (Hcy-on) conformation

##### Coupled/Tandem Chemical Tuning and Biochemical Tuning can steer the *t*MS^HcyFolCapCob^ ensemble towards a Cap-off state

Having capitalized on the tridentate approach of rational non-native cofactor, truncated construct, and protein engineering to capture the transient Fol-on conformation via crystallography, we next sought to capture the elusive homocysteine methylation conformation (Hcy-on). As before, a construct lacking the Act domain was used (*t*MS^HcyFolCapCob^). We found that using cobalamin analogs that contained bulkier, non-alkyl upper axial ligands with this construct was insufficient to destabilize the Cap-on conformation, as judged by the crystallographic capture of multiple Cap-on, His-on, pre-Hcy-on structures, UV-Vis spectroscopy, and trypsin-mediated limited proteolysis to determine conformational changes (Supplementary Fig. 9). Therefore, the tandem approach of strategic mutations *and* non-native cofactor loading was employed. The serendipitous capture of three Cap-on states allowed us to leverage new structural insights to strategically steer the ensemble away from the Cap-on state through steric repulsion (bulky upper axial non-alkyl ligands for Cbl to destabilize the Cap-domain provided a hydrophobic pocket, F713, L720, V723), disruption of Cap:Cob contacts (N774H) and Hcy:Cob contacts (R268A, R826A), removal and alteration of the helical nature of the Fol:Cap linker (D651A, P652A, G653A), and removal of a residue involved in the so-called ligating triad crucial for His-on coordination (D759A) (Supplementary Fig. 10).

To narrow the choice of the upper axial ligand for the cobalamin cofactor, conformational changes of *t*MS^HcyFolCapCob^, in the presence or absence of cobalamin, were visualized by limited proteolysis followed by SDS-PAGE (Supplementary Fig. 9). In tandem, the nature of how these cofactor analogs were bound to our designed *t*MS^HcyFolCapCob^ mutants, and their resulting coordinative environments were studied via UV-Vis spectroscopy (Fig. 5e). *t*MS^HcyFolCapCob^ exhibits increased susceptibility to trypsin in the presence of cobalamins with non-alkyl upper axial ligands (e.g. γ-carboxypropyl, cpCbl, aminoethyl, aeCbl, etc.). Intriguingly, alkyl cobalamin holoenzymes yielded similar digest patterns to the apoenzyme, though the use of mutants confirmed that the construct design principle was indeed shifting the conformational ensemble (Supplementary Fig. 9).

**Figure 5.**
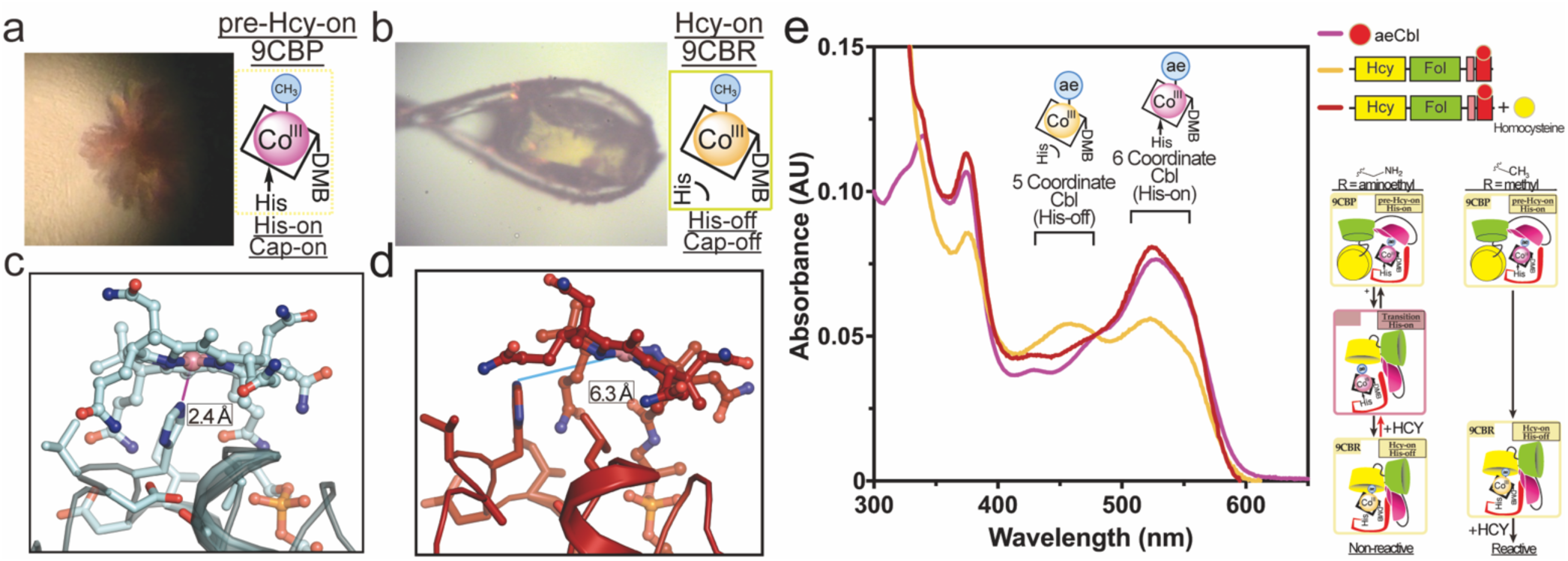
Fol:Cap linker helical melting defines structural transition associated with homocysteine methylation (pre-Homocysteine-on, Cap-on to Homocysteine-on, Cap-off). **a** Color of the crystal which captured the pre-Hcy-on (9CBP) conformation versus **b** that of the crystal used to capture the Hcy-on (9CBQ) conformation. Note the change in color from red to yellow associated with the His-on to His-off ligation associated with the Co(III) state. **c, d** The cobalamin cofactor is displaced ∼3.8 Å laterally and is bound His-off in the catalytic conformation (**c**, 2.32 Å versus **d**, 6.32 Å, ∼Δ4 Å). **e** Free aminoethylcobalamin (aeCbl) shows a peak at approximately 530 nm, characteristic of six-coordinate alkyl-cobalamin (magenta). Upon binding to *t*MS, the aeCbl spectrum exhibits an additional peak at around 460 nm, suggesting that a fraction of aeCbl binds in the His-off mode (five-coordinate) (golden yellow). Furthermore, the addition of excess exogenous homocysteine to the sample (tMS^HcyFolCapCobmut2^•aeCbl) induces changes in the spectrum (brick red), shifting the equilibrium back to the His-on mode, presumably due to the induction of steric clashes; its non-reactive nature could likewise cause a shift in the conformational equilibrium towards a His-on, Cap-on state to protect the cofactor (left panel). Given that cobalamin in the Cap-on state is known to adopt a His-on, six-coordinate state, it is proposed that cobalamin in the Hcy-on conformation adopts a His-off five-coordinate state. Data in panel **e** is of a representative experiment, which has been repeated ≥3 times.

The differences in digest patterns indicate that the combination of non-native cofactors (chemical tuning), coupled with mutagenesis (biochemical tuning), was steering the ensemble towards a different set of states: the protease accessibility of interdomain linkers could be due to uncapping the cobalamin domain, helical melting of the Fol:Cap linker, and/or rearrangement encouraged by the presence of the bulky non-alkyl groups of cpCbl and aeCbl. Local changes in the electrostatics, particularly between the Hcy and Cob domains (Hcy:Cob contacts, R268A, R826A), could trigger linker rearrangement and transition from the capped pre-Hcy-on state to the uncapped Hcy-on state, where part of the Hcy:Fol linker wedged between the Cap:Cob domains (Gln364, Glu365, His408 interact with two of the amide arms of Cbl), becomes accessible (Supplementary Fig. 10).

In light of these findings, mutations designed to disrupt the helical nature of the Fol:Cap linker (D651A, P652A, G653A), favor His-off cobalamin binding (D759A), and that disrupted stabilizing interactions between the Hcy:Cob domain (R268A, R826A) were introduced (Supplementary Fig. 10), with aeCbl used as the non-reactive cobalamin of choice. Using this mutated *t*MS construct and alternate cofactor coupling, we were able to crystallize *t*MS^HcyFolCapCobmut2^, solving the structure to 2.38 Å (Figs. 2b and 4f, Supplementary Table 1).

In the Hcy-on conformation, the Hcy-domain is positioned over the Cob-domain, placing the HCY-binding site directly above the β-face of the cobalamin cofactor (Figs. 4f and g). The cobalamin cofactor is bound in the His-off state, where His761-Nε2 is found ∼6.32 Å from Co (Fig. 4g, inset). This drastic increase in the distance between the ligating His761 residue and the cobalamin cofactor (∼Δ4 Å as compared to the Cap-on, pre-Hcy-on conformation, Fig. 4b, inset) results in the lateral shift of cobalamin ∼3.8 Å relative to its pre-catalytic state with a concurrent ∼20° tilt, culminating in a change from a distorted to more planar corrin ring (Supplementary Figs. 11, 12). This lateral shift of the cobalamin cofactor allows it to orient itself deeper into the Hcy domain, priming it for catalysis, all mediated via the His-on to His-off ligation resulting from the pre-Hcy-on to Hcy-on transition. Indeed, crystals were found to be yellow in color, as compared to the red/pink color observed for the pre-Hcy-on crystals (Figs. 5a and b, Supplementary Fig. 11 a and b). The corresponding changes associated with the cobalamin cofactor are analogous to those observed in previous MS structures in the reactivation conformation (Act-on), particularly the His-on versus His-off states/transition^7,14,23,24^.

The Cap domain is similarly displaced ∼25 Å relative to the Cap-on states^15,16^ lying to the periphery of the Cob domain (8SSC, 8SSD, 8SSE, Supplementary Fig. 3)^14^, validating the notion that the Cap-on to Cap-off transition and resulting position of the displaced Cap domain is specified by the enzyme. The rearrangement of the cobalamin cofactor and its associated domain (Cob) is even more dramatic, transversing 45 Å after uncapping (Supplementary Figs. 5b and 12a), with the bound cobalamin cofactor traveling an astounding 40 Å. The Fol:Cap linker, previously found as highly-structured helix, undergoes a helix-loop transition, resulting in the complete helical melting of Linker II (Fig. 5e), further validating the role of interdomain linkers in large domain rearrangements^14^. Starting from a Cap-on state (9CBP), this helical-hinge must undergo a complete helix-loop transition to form the Hcy-on ternary catalytic complex. In this case, only one of the residues at the N-terminal portion of the linker remains part of a helix, with the rest of the linker (16 residues) forming a loop; this results in a corresponding 30 Å shift of the linker and 60° rotation of the HcyFol domain relative to the Cob domain (Figs. 5c and e). Coupled with the Cap-off transition, this allows for the cobalamin cofactor to be completely buried by the new Hcy:Cob intradomain interface/surface (buried surface area of ∼1361 Å^2^, 89.1% total versus ∼1246 Å^2^, 82.8% total in the assigned pre-Hcy-on Cap-on state, Supplementary Fig. 3).

Inspection of the electron density maps (2Fo-Fc, Supplementary Fig. 7d) and composite omit electron density maps (Fo-Fc, Fig. 8b) showed electron density on the β-face of cobalamin. As such, the upper axial ligand of aeCbl (aminoethyl, Ligand ID NEH) was modelled. The upper axial ligand was found to form two hydrogen bonding interactions with substrate binding site residues, Cys256 and Cys322 (2.6 and 3.5 Å, respectively, Supplementary Fig. 8b) while also coordinating to Zn (2.3 Å). Homocysteine activation is tied to the Lewis-acid activation by Zn, which is coordinated by three cysteine residues (Cys256, Cys322, and Cys323) (Supplementary Fig. 13)^21^. Homocysteine methylation/methionine formation has been postulated to occur via an S_N_2 mechanism whereby Zn primes homocysteine for nucleophilic abstraction of the methyl group from methylcobalamin^25,26,27^. While HCY was found bound in the pre-Hcy-on structure (9CBP), HCY was not observed in the solved Hcy-on crystal structure (Supplementary Figs. 19a and 22b).

##### Capture of the Elusive Hcy-on Conformation reveals how homocysteine methylation can occur

We modeled HCY into the active site and found that the sulfur moiety of HCY is in-line with the cofactor, ∼4.4 Å from Co (Fig. 4g, inset, ∼155° angle between the thiol of homocysteine and cobalt center (Supplementary Figs. 12b and 13a). This is an arrangement required for methyl transfer in MS^21,22^. Modeling methylcobalamin in the active site shows that the distance between the sulfur/thiolate moiety and the methyl group of methylcobalamin is ∼2.1 Å. Notably, the thiol moiety of the modeled HCY ligand occupies a position similar to that of the amino moiety of the aminoethyl upper ligand, likewise ligating Zn while also occupying a position almost equidistant from the cobalt center (4.4 Å for HCY versus 4.3 for aminoethyl, ∼Δ0.1 Å, Supplementary Figs. 8b, 12b, and 13b). The binding of aeCbl directly competes with HCY for the free coordination site on the active site zinc. The cofactor also interacts with two of the three cysteine residues (Cys256, Cys322) that coordinate this zinc. Proper HCY-Zn binding is critical because it facilitates the Lewis-acid mediated activation of a thiol on HCY via zinc coordination, the mechanism for which is thought to involve zinc inversion (Supplementary Fig. 13)^26^.

Both the arrangement and proximity of the reacting centers indicate that the captured state represents a catalytically competent structure. The cobalamin cofactor, aeCbl, is found bound in a His-off state, where His761 (Nε2) is oriented away from the cobalt center, ∼4 Å away from the corresponding Cap-on, His-on state (pre-Hcy-on). In total, this structure represents the Hcy-on (Hcy:Cob) state, one that is catalytically competent for HCY methylation and which is adopted for the methyl transfer from methylcobalamin to homocysteine to yield methionine and Cob(I) (Reaction II, Fig. 1a), indicating that the combination of non-native cofactors and rational protein engineering allowed for the crystallographic snapshot of a transient complex.

The change in coordination of the cobalamin cofactor, from hexacoordinate His-on to pentacoordinate His-off, was unexpected, as the active methylating agent involved in homocysteine methylation/methionine formation was proposed to be His-on MeCbl (Co(III)); however, the change in coordination to His-off ligation allowed for a coupled 3.8 Å lateral translation and corresponding 18° tilt of the cobalamin cofactor towards the Hcy domain and its active site, in a manner strikingly similar to that previously observed in the reactivation conformation (Act-on)^7,14,23,24^. The complete restructuring of the Fol:Cap linker (Linker II) via helical melting, from a helix in the pre-catalytic state to a loop in the active state, provided a striking demonstration that interdomain linkers, particularly Linker II in the case of MS, can be involved in large domain rearrangements required for catalysis. Indeed, it seems that instead of relying on the restructuring of entire domains, hinge regions in the form of interdomain linkers can allow for adjacent domains to move as rigid bodies, ensuring preformed active sites can shuttle their precious cargo unperturbed, awaiting their next stop.

#### The Fol domain is required for Homocysteine Methylation

The restructuring of the Fol:Cap linker between pre-catalytic and catalytic states highlights one of the proposed roles associated with the Fol domain in guiding the conformational changes MS must undergo to achieve catalysis. The dramatic reorientation of the Fol domain between Cap-on states provides an alternate method by which MS can sample different orientations *en route* to catalytic states, all without the need to deprotect the reactive cobalamin cofactor via its uncapping. The assignment of 9CBP as the pre-Hcy-on state, versus the ‘resting state’ proposed previously^16^, required us to interrogate the biochemical basis for the proposed importance of the Fol domain, particularly in the homocysteine methyltransferase reaction.

Therefore, we employed a UV-Vis assay using the Hcy, Hcy:Fol, and Cap:Cob domains coupled in trans^10^. The excised domains were chosen to provide a simpler, minimal model sufficient to study the ability of MS to perform homocysteine methylation^11^. Methylcobalamin consumption was monitored using UV-Vis to quantify the activity of various kinds of *t*MS domains (Fig. 6). The Hcy:Fol didomain and the isolated Hcy domain of tMS (designated *t*MS^HcyFol^ and *t*MS^Hcy^, respectively) contain the homocysteine-binding sites. Methylcobalamin was complexed with the Cap:Cob didomain (*t*MS^CapCob^). Consumption of exogenous methylcobalamin was detected in the presence of both *t*MS ^HcyFol^ and *t*MS^Hcy^ (Fig. 6) but not in their absence (Supplementary Fig. 14), while consumption of methylcobalamin when bound to *t*MS^CapCob^ was observed with *t*MS^HcyFol^ (Fig. 6c) but not with *t*MS^Hcy^ (Fig. 6d). These data suggest that the Fol domain is important for the formation of the Hcy-on conformation, and without it, a catalytically competent Hcy-on ternary complex cannot be formed^8,9,16,28^.

**Figure 6.**
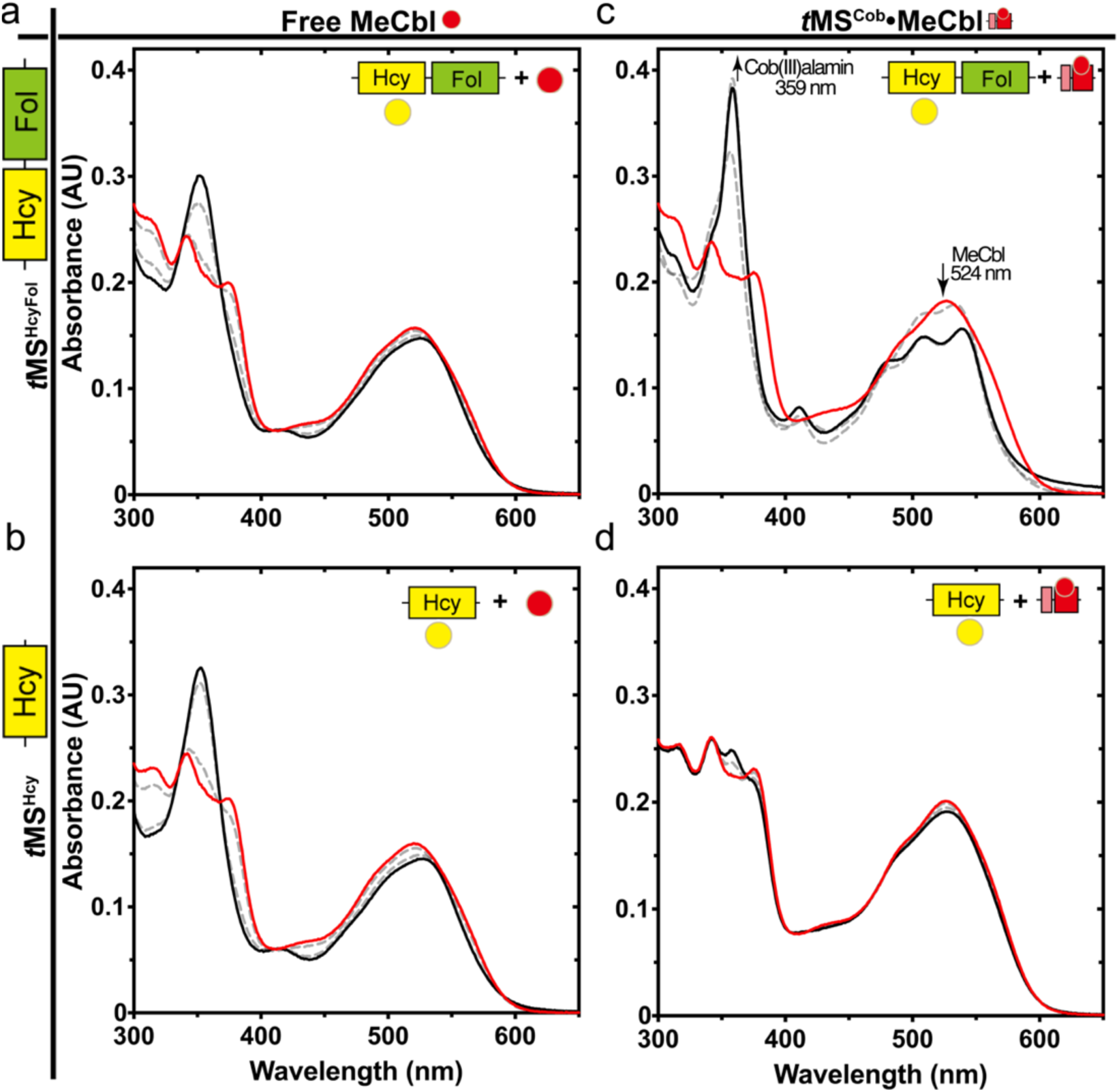
Homocysteine methylation requires the Fol domain. **a** Effects of *t*MS domain fragments on methylcobalamin:homocysteine methyltransferase reaction. The time-dependent spectral changes of methylcobalamin (MeCbl) in the presence of homocysteine were monitored at 50 °C for up to 10 minutes after adding homocysteine (final concentration of 100 µM) using 20 µM of free (non-protein bound) MeCbl incubated in the presence of 2 µM *t*MS constructs (**a** and **b**) or MeCbl (23 µM) was bound to the isolated *t*MS^CapCob^ domains first (**c** and **d**). Red lines represent the initial time point, with black lines representing the endpoint (final 10 min); gray lines represent intermediary time points (2, 5 min). Note that the presence of the Fol domain is required for activity when methylcobalamin is protein bound (**c**). Free homocysteine and methylcobalamin did not react (Supplementary Fig. 14), similar to *t*MS bound cobalamin (**d**). Free cobalamin can react minimally with *t*MS bound homocysteine, presumably due to the lack of the Cap domain and the ability of the Hcy domain to activate homocysteine. Data are of representative experiments, which have been repeated ≥2 times.

##### His-off Cob(III)alamin is the active methylating agent required for homocysteine methylation

The observed change in coordination of the cobalamin cofactor, from hexacoordinate His-on (pre-Hcy-on) to pentacoordinate His-off (Hcy-on) was unexpected, as the active methylating agent involved in homocysteine methylation/methionine formation was proposed to be His-on MeCbl (Co(III), Fig. 1a). Methionine synthase binds cobalamin in a so-called His-on, base-off manner, where the dimethylbenzimidazole (DBI) tail inserts into the conserved nucleotide binding cleft of the Cob domain and is replaced by His761 in the lower axial position. Comparison of the cobalamin binding pocket (using the Cob domain as the reference for the structure alignment) reveals a stark change in His761-Nε2 and Co distance (2.32 Å versus 6.32 Å, ∼Δ4 Å) between the pre-Hcy-on and Hcy-on (Figs. 4b and g, Figs. 5c and d). Notably, no repositioning of the DBI moiety is observed between the pre-Hcy-on and Hcy-on structures (Supplementary Fig. 11d), indicating that the cobalamin cofactor conformational change observed is limited to the corrin ring. The Hcy-on state and its associated His-off ligation allows for the Cbl cofactor to move 3.8 Å laterally into the Hcy domain and its active site with an associated 4.2 Å tilt, relieving any potential steric clashes (Supplementary Fig. 11).

To further explore the nature of how the cofactor analogs were bound to our designed *t*MS^HcyFolCapCob^ mutants, their resulting coordinative environments were studied. UV-Vis analysis of free cofactor analogs versus their *t*MS^HcyFolCapCob^ bound states verified that the free cofactor predominantly adopts a Base-on, hexacoordinate state (Co(III)), while revealing that their protein bound state shifts to adopt both His-on and His-off binding modes (Fig. 5e, golden yellow line). Protein bound aminoethylcobalamin (aeCbl) resulted in the most pronounced effect, yielding a visible color change from yellow (His-off, 5 coordinate, Co(III)) (Figs. 5a and b, Supplementary Figs. 11a and b) to red/pink (presumably Cap-on, His-on, 6 coordinate, Co(III), Fig. 5e, brick red line). The addition of exogenous homocysteine (HCY) to a solution of the construct used to capture the Hcy-on state complexed with aeCbl induces a shift in the spectrum consistent with a change towards the His-on form. Given that HCY cannot react with aeCbl and occupies the substrate binding site in the Hcy domain, the aminoethyl group on cobalamin likely results in a steric clash with HCY, disfavoring the Hcy-on conformation (Fig. 5e). As a result, homocysteine pushes the conformational equilibrium *back* toward the pre-Hcy-on (Cap-on) conformation. The resulting reversion to the His-on, Cap-on state upon addition of homocysteine, despite the use of a non-reactive cobalamin, could also function to protect the cofactor from sampling a non-reactive/productive conformation (Fig. 5e).

Notably, the color of the crystals used to solve the Hcy-on conformation was yellow, while that used to solve the pre-Hcy-on form was a red (Figs. 5a and Supplementary Fig. 11a). The Co atom in unbound MeCbl, cpCbl, and aeCbl is hexacoordinate, resulting in a red color in a neutral solution. In these cofactor analogs, the Co atom is coordinated by four nitrogen atoms from the corrin ring, with the non-native ligand binding to Co from the β face, and the DBI moiety coordinated from the α face. Upon binding to MS, the lower ligand of cobalamin (DBI) is replaced by His761 ligation. MS accommodates the DBI moiety in a deep pocket within the Cob domain, effectively anchoring cobalamin to the enzyme. The Co-His coordination undergoes an on/off cycle during catalysis. When the MS•MeCbl complex has six-coordinate Co(III), it exhibits a red color; however, if His residue dissociation occurs, it would result in five-coordinate Co(III) and appear yellow. The yellow color of the crystal, coupled with the UV-Vis spectra, indicate that MS binds cobalamin in a His-off mode in the Hcy-on conformation, and that the observed pentacoordinate Co(III) state found in the Hcy-on structure is not a crystallographic artefact.

The use of the chemically inert aeCbl, coupled with the intriguing observation of a yellow color typically associated with cob(II)alamin (and thus inactivation and the reactivation state), implied another set of cobalamin states were being sampled in solution and *in crystallo*. His-off, pentacoordinate cob(III)alamin, which also presents as a yellow color, provides a neat distinction between the pre-catalytic, Cap-on state (His-on ligation) and the catalytic, Cap-off state (His-off ligation), (Supplementary Figs. 11 and 13). This adds another layer of intricacy to the already important role His plays in guiding, telegraphing, and governing cobalamin movement in MS, with the extensive list now including proper gating between pre-catalytic (Cap-on, His-on) and catalytic (Cap-off, His-off) states.

##### The Fol domain governs access to catalytic conformations

Our previously captured *t*MS structures (8SSC, 8SSD, and 8SSE) highlighted the preformed cobalamin pocket/cleft, which is solvent-exposed and thus readily accessible even *in crystallo*^14^. Similarly, previous structures of MS excised domains revealed that the Hcy and Fol domains were preformed, with minimal microenvironment changes in the substrate binding pocket induced upon ligand/substrate binding^21,29,30^. Even so, MS must exert global control over the conformational rearrangements that would allow access to the β face of the cobalamin cofactor (i.e. the formation of a ternary complex with the Cob domain and a substrate domain) following substrate binding^28^.

The first step in the MS catalytic cycle is the methylation of HCY using methylcob(III)alamin to form methionine and Co(I)alamin, with Co(I)alamin used to demethylate MTF and regenerate methylcob(III)alamin in the second step. Intriguingly, while purification of *t*MS routinely results in copurification with MTF^21^, use of the *t*MS^FolCapCob^ tridomain did not yield any MTF-bound structure. However, the use of *t*MS^HcyFolCapCob^ tetradomain did yield a folate-bound structure, modelled as *tetrahydrofolate* (Ligand ID THG), indicating that the captured pre-Hcy-on (Cap-on, 9CBP) and Hcy-on (9CBR) states copurified with the folate demethylation byproduct (Supplementary Fig. 24). The finding that only *t*MS^HcyFol^ was active in the trans assay using methylcobalamin loaded *t*MS^CapCob^ indicates that the Fol domain plays a pivotal role in guiding the conformation rearrangement necessary for the formation of the Hcy-on ternary complex (Fig. 6), likely through destabilizing the Cap-on state (uncapping activity)^16,21^. Whether the Hcy domain itself is necessary for proper MTF-binding and reactivity remains unknown. The flexible interdomain linker region connecting Fol:Cap (Linker II) must be considered as an additional, significant factor. Its coil-helix transition is a tantalizing explanation for why the Fol domain and Linker II are necessary for the formation of the Hcy-on conformation and provides a guided mechanism by which MS integrates substrate-binding information with cofactor status to undergo the pre-requisite rearrangements necessary for catalysis^8,9,16,28^.

##### Linker regions guide domain rearrangements centered on the cobalamin cofactor

The captured structures provide the first experimental validation that interdomain linkers play an important role in facilitating the formation of specific conformations. The Hcy:Fol linker (Linker I), being rigid, is not found to change significantly^29^. The Fol:Cap linker (Linker II) seems to play an outsized role in allowing for the formation of catalytically-competent ternary complexes in the MS catalytic cycle, along with intermediates contained therein (Supplementary Figs. 1, 2, and 3). Alignment of the catalytic structures, including the previously obtained Act-on reactivation structures^14^, using the Cob domain as reference, indicate that the Cap-off Cob domain interface can vary, particularly with the positioning of the Cap domain (RMSD = 2.414 Å, 202 to 202 atoms for 9CBQ and 8SSC, RMSD = 3.146 Å, 214 to 214 atoms for 9CBR and 8SSC, Supplementary Fig. 3), though the Cob and Cap domains from each conformation show they are superimposable (Supplementary Figs. 3 and 16).

The structural rearrangements observed center on the Cap:Cob domains, with substrate-binding domains vying for access to the protected (Cap-on) Cob domain (Supplementary Figs. 1, 2, and 3). Concurrently, the Cap:Cob linker (Linker III) could play a role in mediating the uncapping motion required for cobalamin access that is interdependent on the cofactor redox state and coordinate number, placing the Cap domain in a predetermined position. Morphing of pre-catalytic structures (Cap-on states, Supplementary Movie 2) and catalytic structures (Cap-off, Fol-on, Hcy-on, Supplementary Movies 3 and 4) show the Fol:Cap linker acting as a swiveling point around which the Hcy:Fol domains, presumably, use the Fol domain to initiate “uncapping” of the Cob domain. Even in the Cap-on structures, the Cap:Cob domain remains relatively static (Supplementary Figs. 1, Supplementary Fig. 16, Supplementary Movie 1). It seems that the Fol domain, in conjunction with its associated Fol:Cap linker, plays a critical role in restructuring and thereafter guiding the conformational ensemble during catalysis, while the Cap:Cob linker allows for proper repositioning of the Cap domain in one of two states (Cap-on or Cap-off).

## Discussion

While the conformational plasticity of MS in solution is well-established, the precise regulatory mechanisms governing these domain arrangements remain elusive^6,7,8,9^. In this work, we have leveraged our previously discovered thermophilic methionine synthase homolog (*t*MS)^14^ to structurally probe the conformations that underpin MS catalysis. This work has expanded the known conformational landscape of MS, revealing Cap-on states that we propose function as pre-catalytic intermediates to the catalytic cycle. Furthermore, we have captured the first catalytic structures of MS poised for MTF demethylation (Fol-on) and HCY methylation (Hcy-on), providing novel insights into the residues and global rearrangements that dictate catalytic activity. This study addresses long-standing questions in the field that have been the subject of research for decades and contributes to a deeper understanding of MS, potentially leading to its exploitation as a biocatalytic tool.

Our unified structural and functional model proposes that Cap-on states act as regulatory checkpoints, controlling access to the catalytically active Cap-off conformations. The precise mechanisms underlying transitions between catalytic states, however, warrants further investigation. Our model outlines two potential pathways: direct transitions between Hcy-on and Fol-on states (Supplementary Fig. 17, Model 1, Supplementary Fig. 15, Supplementary Movie 5) or recapping-dependent transitions involving multiple Cap-on states (Supplementary Fig. 17, Model 2, Supplementary Movies 6 and 7). In both scenarios, the Fol:Cap (Linker II) hinge orchestrates/helix-loop transition allow for the necessary domain rearrangements. If transitions between Hcy-on and Fol-on states can occur without recapping, a single Cap-on to Cap-off transition would be sufficient for catalytic entry. Thereafter, the Hcy:Fol domains can use rigid body motions as guided by Linker II to form their respective ternary complexes (Fig. 7, Supplementary Fig. 17, Model 1, Supplementary Figs. 15 and 18, Supplementary Movie 5). Conversely, if recapping is obligatory after catalysis, multiple Cap-on transitions, also orchestrated by Linker II helix-loop transitions, would be necessary for gated re-entry into the catalytic cycle (Fig. 7, Supplementary Fig. 17, Model 2, Supplementary Figs. 15 and 18, Supplementary Movies 6 and 7). The contrasting features of these models offer a valuable framework that facilitates hypothesis-driven experimentation and iterative model refinement, allowing for a more comprehensive understanding of the intricate structural ensemble that governs MS’s multifaceted functionality. Currently, the catalytic efficiency of MS leads us to favor Model 1, where transitions between catalytic conformations are allowed, given that biochemical studies have shown that the rate limiting steps in MS are associated with conformational changes, and not its chemistry^8,28,29^.

**Figure 7.**
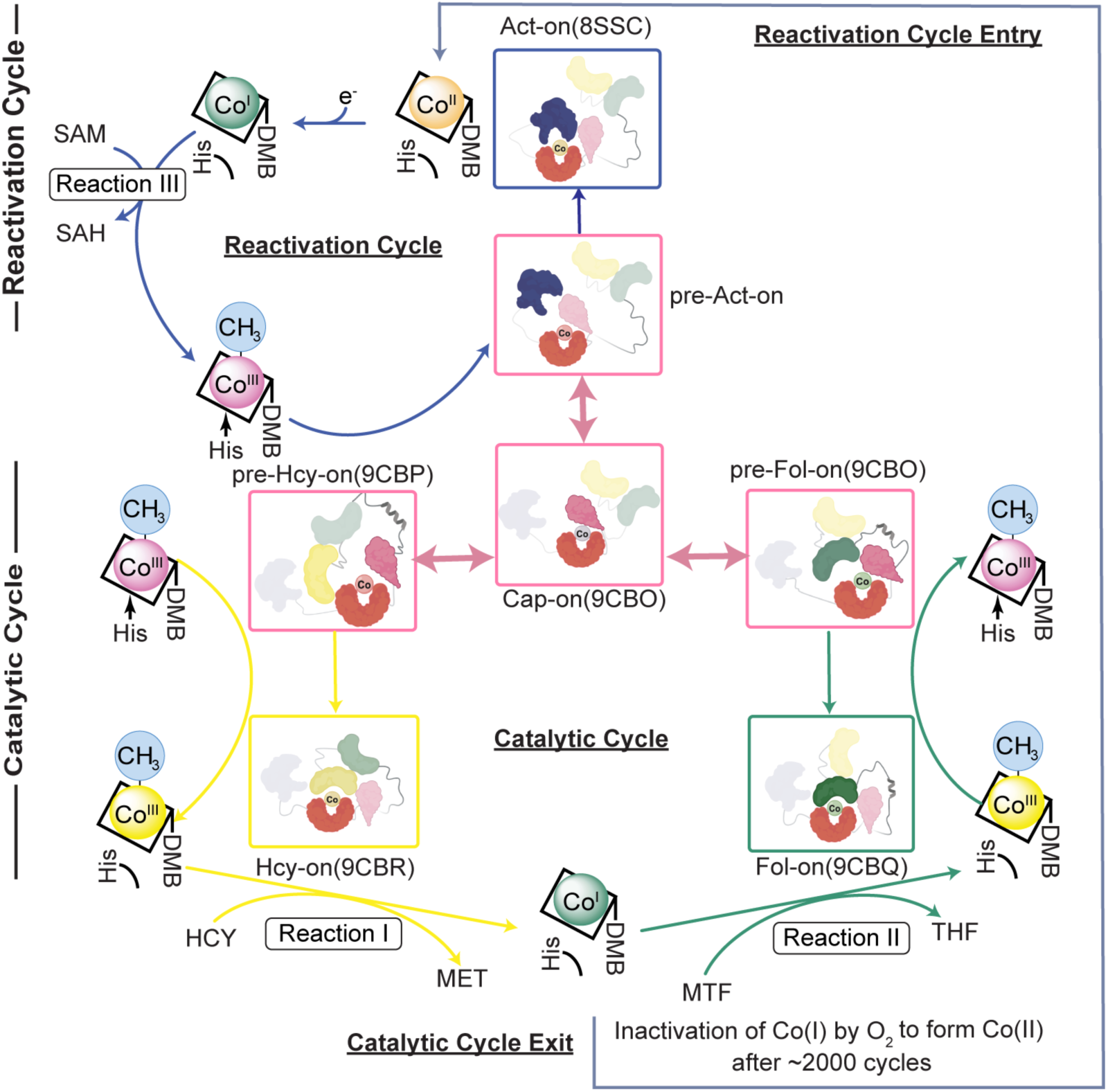
Conformational transitions during catalysis are mediated by the Hcy:Fol linker and signaled by His761. Methionine synthase can cycle through Cap-on, His-on states that function as pre-catalytic conformations, obviating the need for uncapping. Linker II (Fol:Cap linker) allows for facile transition between these pre-catalytic states, what we term ‘resetting’. In our pre-catalytic conformations, hexacoordinate, His-on methylcobalamin (Co(III)) highlights its role as a bridge between the primary catalytic and reactivation cycles. Depending on the cobalamin cofactor’s status (redox state, coordinate number) and substrate presence, the appropriate pre-catalytic state can grant entry to reactive conformations by uncapping and allowing for cobalamin flexibility by binding in the His-off state. Notably, the active methylating agent for methionine formation has been revised from hexa-to penta-coordinate His-off methylcob(III)alamin. Each methylation is associated with a distinct cobalamin redox state (Co(III) for homocysteine methylation, Reaction I; Co(I) for methyltetrahydrofolate demethylation, Reaction II; Co(II) for reductive methylation via SAM/AdoMet, Reaction III). The primary catalytic cycle of methionine synthase involves cycling between Co(III) and Co(I) states; Co(I) production acts to lock MS to either the catalytic cycle (homocysteine methylation) or the reactivation cycle (Co(II) reductive alkylation) until methylcobalamin is regenerated or Co(I) undergoes oxidative inactivation.

Our findings reveal a five-coordinate His-off methylcob(III)alamin intermediate during catalysis (Hcy-on, Figs. 4, and 5, Supplementary Figs. 11, 12, and 13)^28^ and suggest the captured pentacoordinate His-on Co(III) state in the Fol-on complex represents a post-catalytic configuration, where cpCbl is in the Co(III) state and thus mimics methylcob(III)alamin (Supplementary Fig. 4). While a four-coordinate Co(I) species is implicated in folate demethylation^28^, the role of the five-coordinate His-off cobalamin post-demethylation warrants further study. Although pentacoordinate His-on cob(I)alamin has not been directly observed, there is literature precedence for its potential role in enhancing folate demethylation, presumably through transition state stabilization (mimicking hexacoordinate methylcobalamin)^13,31^. Biochemical studies of cobalamin-dependent methyltransferases indicate His-on ligation (hexacoordinate) stabilizes methylcobalamin rather than being crucial for methyl transfer, while His-off, pentacoordinate methylcobalamin would promote homocysteine methyl transfer^13,31,32,33,34,35,36,37,38^. An additional factor that must be considered are distortions in the corrin ring of the cobalamin cofactor, which are associated with the cobalt’s oxidation state and coordination number/ligation status; these changes are, in turn, associated with new stabilizing (e.g. hydrogen bond networks between cobalamin amide/propionamide arms and protein residues) and destabilizing (e.g. steric clashes between the corrin ring and protein residues) interactions that can result in an overall net stabilizing or destabilizing effect^39^.

Significantly, all Cap-on structures observed to date exhibit alkylcob(III)alamin bound in the His-on state. Previous biochemical work (Fig. 1d) has settled on a model where a minimal set of four conformations were proposed to compromise the MS catalytic ensemble, along with the notion that methylcobalamin serves as a key branching point in both the catalytic and reactivation cycles^8^; while His-off cobalamin was emphasized for its importance in steering the conformational ensemble towards reactivation, we note that His-off methylcob(III)alamin is the active methylating agent during homocysteine methylation, and that His-off binding of cobalamin is associated with Cap-off transitions. The catalytic cycle of methionine synthase could be reconstituted *in trans* using two mutants that uncoupled homocysteine methylation and folate demethylation, positing ‘turnover in two encounters’, *i.e.* multiple uncapping events (Model 2)^13^. Using our structural data, we offer a nuanced revision to the model^8,13^. Methylcobalamin generated from folate demethylation can either proceed to the Hcy-on state in the presence of homocysteine (Fig. 7, Supplementary Fig. 17, Model 1, Supplementary Figs. 15 and 18) or ‘recap’ to safeguard the cofactor (Fig. 7, Supplementary Fig. 17, Model 2, Supplementary Figs. 15 and 18, Supplementary Movie 8). Similarly, methylcobalamin produced during reductive methylation can *exit* the reactivation cycle, likely through recapping, before re-entering the catalytic cycle. In contrast, cob(I)alamin is exclusively generated during homocysteine methylation or cob(II)alamin reduction. Once formed, cob(I)alamin is committed to its respective cycle (catalytic or reactivation) until methylcobalamin is regenerated or oxidative inactivation to cob(II)alamin occurs. This could explain the observed specificity of cob(I)alamin reactivity towards MTF in the catalytic cycle and AdoMet in the reactivation cycle^8^. Our models highlight the central role of hexacoordinate His-on methylcobalamin as a bridge between the catalytic and reactivation cycles, with the generation of Co(I) representing a commitment step to either cycle. Future work will focus on elucidating the fate of Co(I) during catalysis and further characterizing the dynamic interplay between His ligation, cobalamin reactivity, and structural transitions; said approach will also aid in characterizing the nature of the cobalamin cofactor during holoenzyme formation/cofactor loading and the conformation(s) associated therein.

Overall, this revised mechanistic and structural model synthesizes the wealth of biochemical data with unprecedented molecular level structural insights, providing a new unified framework with which to understand cobalamin-dependent methyltranferases such as methionine synthase (Fig. 7, Supplementary Figs. 15, 17, and 18). Starting from one of three Cap-on, His-on states (Fig. 7), MS can alternate between these three pre-catalytic conformations through rigid body motion of the Hcy:Fol domain towards the Cob domain (pre-Fol-on, pre-Hcy-on) or away from it (pre-Act-on/resting), all mediated by the remarkable polymorphism of Linker II (Supplementary Fig. 1, Supplementary Movie 1). Linker II is a versatile element capable of adopting at least three distinct structural conformations. This flexibility enables it to rearrange and reposition the domains attached to its termini. His-off ligation of cobalamin acts as a switch to allow for gated entry from pre-catalytic to catalytic transitions and vice-versa (Fig. 7). Structural transitions to catalytic states are primarily cofactor dependent, signaled by His ligation and the coordination number/environment, mediated by the Fol:Cap hinge (Linker II), programmed rigid body movement of the Cap:Cob domain via the Cap:Cob linker (Linker III), and substrate-bound domain association to form a ternary complex ready for catalysis. In this manner, MS harnesses the catalytic potential of cobalamin while avoiding futile cycling, leveraging unheralded structural motifs to orchestrate the dramatic rearrangements required to perform three improbable chemistries.

## Methods

### Cobalamin Analogue Synthesis

Cyanocobalamin (CNCbl) was purchased from Sigma/MilliporeSigma. cobalamin analogues were synthesized following previously established methods^40,41,42^. Briefly, CNCbl reduced by NaBH_4_ was mixed with 4-chlorobutyric acid (Sigma/MilliporeSigma), 2-chloroethylamine hydrochloride (Sigma/MilliporeSigma), or n-propyl iodide (Sigma/MilliporeSigma) to yield γ-carboxypropyl-cobalamin (cpCbl), aminoethyl-cobalamin (aeCbl), and propyl-cobalamin (prCbl), respectively.

### Construction of *t*MS Expression Vectors

Expression vectors for truncated *t*MS expression were generated using the ligation-independent cloning (LIC). Molecular cloning was conducted using pMCSG7 as the expression plasmid DNA vector and *E. coli* XL1-Blue as the host cell. All constructed bacterial expression vectors were designed to produce truncated *t*MS with a hexa-histidine tag and a Tobacco Etch Virus (TEV) protease cleavable sequence at the N-terminus.

The genes encoding truncated *t*MS were amplified by PCR, using pET(*t*MS^wt^) as the template, obtained from the Riken BioResource Center^43^. Subsequently, these amplicons were integrated into a pMCSG7 vector using the ligation-independent cloning technique to generate truncated *t*MS expression vectors. To express tridomain *t*MS, pMCSG7(*t*MS^FolCapCob^) was constructed, encompassing the Fol, Cap, and Cob domains; pMCSG7(*t*MS^HcyFolCapCob^) was prepared in a similar fashion to express tetradomain *t*MS, containing all domains except for the Act domain. Didomain and single domain vectors to express the constructs used for assays were similarly generated, designated pMCSG7(*t*MS^CapCob^), pMCSG7(*t*MS^HcyFol^), and pMCSG7(*t*MS^ΔΝ35Hcy^).

Mutant constructs were made by site-directed mutagenesis (QuikChange, Agilent). Mutant constructs of the tetradomain *t*MS used for crystallography include several mutations: pMCSG7(*t*MS^HcyFolCapCobF110A,E123A,E124A,Y296A,D651A,P652A,G653A^/*t*MS^HcyFolCapCobDPG2A dEE F110A,Y296A^) used for crystallization of pre- Hcy-on (mutant 1/construct 1) and pMCSG7(*t*MS^HcyFolCapCobF110A,E123A,E124A,R268A, D651A,P652A,G653A,D759A,N774H,R826A^/*t*MS^HcyFolCapCobDPG2A dEE dRR F110A,D759A,N774H^) used for crystallization of Hcy-on (mutant 2/construct 2), will be referred to as such in the methods for the sake of simplicity. A complete list of the bacterial strains, plasmids, and primers used in this study are listed in Supplementary Table 2.

### Expression of *t*MS Constructs

*E. coli BL21star*(*DE3)* was transformed with the desired expression vectors for protein expression. *E. coli* transformed with the desired pMCSG7(*t*MS) constructs were propagated at 37 °C in Luria Broth containing 50 µg/mL ampicillin, and protein overexpression was induced using autoinduction media^44,45^. *E. coli BL21star(DE3)* transformed with pMCSG7(*t*MS^FolCapCobD759A^) were propagated at 37 °C in Luria Broth containing 50 µg/mL ampicillin and protein overexpression was induced using Isopropyl β-D-1-thiogalactopyranoside (IPTG) (final concentration of 1 mM). In the case of *t*MS^HcyFolCapCobmut1^, ZnCl_2_ was added to the medium (0.5 mM) and protein overexpression was induced using autoinduction media. Cells were propagated at 30 °C with shaking at 250 rpm overnight prior to harvesting via centrifugation and stored at −80 °C.

### *t*MS Protein Purification

Immobilized metal chelate affinity (IMAC) chromatography was employed to purify recombinant *t*MS (HisTrap Chelating HP, Cytvia, 5 mL), using a NiSO4 charged column for all constructs except for *t*MS^HcyFolCapCob^, where a ZnSO4 charged column was used.

#### *t*MS^FolCapCobD759A•MeCbl^ (pre-Fol-on)

For holoenzyme purification with Mecobalamin (*t*MS^FolCapCobD759A^, *t*MS^CapCob^), the harvested cell pellet was first resuspended in 50 mM potassium phosphate buffer (KPB) (pH 7.4), 0.3 M sodium chloride, (4-5 mL/1 g of pellet), to which lysozyme (0.1 mg/mL) and PMSF (1 mM) were added. The resuspended cell pellet was lysed via sonication (4 °C, 5 s on, 5 s off, 5 min total). The crude lysate was centrifuged (15 min × 3, 10,000 × g, 4 °C) decanting the supernatant to remove any cellular debris/pellet. Methylcobalamin (MeCbl) (MilliporeSigma) was added to the crude supernatant (20 mM) to form holo-*t*MS. The crude extract with Mecobalamin was incubated at 70 °C for 15 min, then centrifuged (15 min, 10,000 × g, 4 °C). The supernatant was pooled and filtered via syringe (0.45 µm), to which imidazole was added (20 mM) and the supernatant was applied/loaded onto a Ni-affinity column (His-trap Chelating HP, Cytvia, 5 mL) equilibrated with 50 mM KPB, pH 7.4, and 20 mM imidazole. The column was washed with 50 mM KPB, pH 7.4, 0.3 M NaCl, 20 mM Imidazole (5 CV), then 50 mM imidazole (5 CV), and the protein was eluted in bulk with buffer consisting of 50 mM KPB, pH 7.4, 0.3 M NaCl, and 80 mM imidazole (8 CV) and 150 mM Imidazole (3 CV). Red-colored fractions were collected and subjected to a TEV digest, with dialysis at 4 °C overnight (50 mM KPB, pH 7.4). The dialyzed sample was incubated at 70 °C for 15 min to quench the TEV digest, then centrifuged (15 min, 10,000 × g, 4 °C). The resulting supernatant was clarified via centrifugation and concentrated via centrifugation in 50 mM KPB, pH 7.4 to yield purified *t*MS^FolCapCobD759A•MeCbl^ (∼10 mg/mL), which was stored at 4 °C or flash-frozen for long-term storage at −80 °C.

#### *t*MS^CapCob•MeCbl^

*t*MS^CapCob^ was prepared similarly, save for the use of a TEV digest. Following IMAC purification and pooling of red-colored fractions, the combined fractions were subject to dialysis at 4 °C overnight (50 mM KPB, pH 7.4). The dialyzed sample was concentrated via centrifugation in 50 mM KPB, pH 7.4 to yield purified *t*MS^CapCob•MeCbl^ (∼10 mg/mL), which was stored at 4 °C or flash-frozen for long-term storage at −80 °C.

#### *t*MS^FolCapCobD762G•cpCbl^ (Fol-on)

Purification of *t*MS^FolCapCobD762G^ was conducted similarly, with no added cobalamin. The harvested cell pellet was resuspended in 50 mM KPB (pH 7.4) (∼5 mL/1 g of the pellet), to which lysozyme (0.1 mg/mL) and PMSF (1 mM) were added. The resuspended cell pellet was lysed via sonication (4 °C, 5 s on, 5 s off, 5 min total). The crude lysate was centrifuged (45 min, 20,000 × g, 4 °C) decanting the supernatant to remove any cellular debris/pellet. To the crude supernatant was added Imidazole (final concentration, 20 mM) before purification using a modified IMAC bulk elution (equilibration buffer: 50 mM KPB, pH 7.4, 20 mM Imidazole, wash buffer: 50 mM KPB, pH 7.4, 0.3 M NaCl, 50 mM Imidazole, elution buffer A: 50 mM KPB, pH 7.4, 0.3 M NaCl, 80 mM Imidazole, elution buffer B: 50 mM KPB, pH 7.4, 0.3 M NaCl, 150 mM Imidazole), with the desired protein eluting at 80 mM and 150 mM Imidazole. Fractions containing protein as confirmed by SDS-PAGE were then pooled, and an ammonium sulfate cut was used to remove any impurities, using an equal volume of saturated ammonium sulfate (∼4 M) to achieve a final concentration of 50% (w/v). The mixture was centrifuged (10,000 × g, 10 min, 4 °C) to pellet any precipitated protein. The supernatant was carefully decanted to avoid disturbing the pellet; the pellet was resuspended in 0.1 M KPB, pH 7.4, 2 mM DTT (10 mL). A TEV digest and overnight dialysis (4 °C, 0.1 M KPB, pH 7.4, 2 mM DTT) followed. To the dialyzed sample containing ∼135 μM apo-*t*MS^FolCapCobD762G^ was added γ-carboxypropyl cobalamin (cpCbl, ∼170 μM) and the TEV digest quenched by heating at 60 °C (15 min), then centrifuged (10 min, 4,000 × g, 4 °C). The resulting supernatant was loaded onto a pre-equilibrated desalting column (PD-10, GE/Cytvia, 50 mM KPB, pH 7.4) to remove excess cpCbl and concentrated via centrifugation in 50 mM KPB, pH 7.4 to yield purified holo*-t*MS^FolCapCobD762G•cpCbl^ (∼10 mg/mL), which was stored at 4 °C or flash-frozen for long-term storage at −80 °C.

Purification of apo constructs (*t*MS^ΔN35Hcy^, *t*MS^HcyFol^) was conducted similarly, with no added cobalamin.

#### *t*MS^ΔN35Hcy^

The harvested cell pellet was resuspended in 50 mM Tris (pH 7.4), 0.1 M NaCl (∼5 mL/1 g of the pellet), to which lysozyme (0.1 mg/mL) and PMSF (0.1 mM) were added. The resuspended cell pellet was lysed via sonication (4 °C, 5 s on, 5 s off, 5 min total). The crude lysate was centrifuged (10 min, 8,000 × g, 4 °C) decanting the supernatant to remove any cellular debris/pellet. The crude supernatant was subjected to a heat-treatment step at 60 °C (15 min), then centrifuged (10 min, 4,000 × g, 4 °C), decanting the supernatant to remove any cellular debris/pellet. To the crude supernatant was added Imidazole (final concentration, 50 mM) purification using a modified IMAC bulk elution (equilibration buffer: 50 mM Tris, pH 7.4, 0.1 M NaCl, 50 mM Imidazole, wash buffer: 50 mM Tris, pH 7.4, 0.3 M NaCl, 50 mM Imidazole, elution buffer: 50 mM Tris, pH 7.4, 0.3 M NaCl, 200 mM Imidazole), with the desired protein eluting at 200 mM Imidazole. Fractions containing protein as confirmed by SDS-PAGE were then pooled. The combined fractions were subject to dialysis at 4 °C overnight (50 mM Tris, pH 7.4), to which TCEP was added (final concentration, 1 mM), followed by a second heat-treatment step at 60 °C (15 min) then centrifuged (10 min, 4,000 × g, 4 °C). The resulting supernatant was concentrated via centrifugation in 50 mM Tris, pH 7.4, 1 mM TCEP to yield purified *t*MS^ΔN35Hcy^ (∼10 mg/mL), which was stored at 4 °C or flash-frozen for long-term storage at −80 °C.

#### *t*MS^HcyFol^

The harvested cell pellet was resuspended in 50 mM KPB (pH 7.4) (∼5 mL/1 g of the pellet), to which lysozyme (0.1 mg/mL) and PMSF (0.1 mM) were added. The resuspended cell pellet was lysed via sonication (4 °C, 5 s on, 5 s off, 5 min total). The crude lysate was centrifuged (10 min, 8,000 × g, 4 °C) decanting the supernatant to remove any cellular debris/pellet. The crude supernatant was subjected to a heat-treatment step at 70 °C (15 min), then centrifuged (10 min, 4,000 × g, 4 °C), decanting the supernatant to remove any cellular debris/pellet. To the crude supernatant was added Imidazole (final concentration, 20 mM) before purification using a modified IMAC bulk elution (equilibration buffer: 50 mM KPB, pH 7.4, 50 mM Imidazole, wash buffer: 50 mM KPB, pH 7.4, 0.3 M NaCl, 50 mM Imidazole, elution buffer: 50 mM Tris, pH 7.4, 0.3 M NaCl, 80 mM Imidazole), with the desired protein eluting at 80 mM Imidazole. Fractions containing protein as confirmed by SDS-PAGE were then pooled. The combined fractions were subject to dialysis at 4 °C overnight (50 mM KPB, pH 7.4). The resulting dialyzed sample was concentrated via centrifugation in 50 mM KPB, pH 7.4 to yield purified *t*MS^HcyFol^ (∼10 mg/mL), which was stored at 4 °C or flash-frozen for long-term storage at −80 °C.

Purification of *t*MS^HcyFolCapCob^ constructs was conducted similarly, without added cobalamin, but using a ZnSO4-loaded IMAC column (GE/Cytvia, 5 mL).

#### *t*MS^HcyFolCapCobmut1•prCbl^ (pre-Hcy-on)

The harvested cell pellet was first resuspended in 50 mM potassium phosphate buffer (KPB) (pH 7.4) (4-5 mL/1 g of pellet), to which lysozyme (0.1 mg/mL) and PMSF (1 mM) were added. The resuspended cell pellet was lysed via sonication (4 °C, 5 s on, 5 s off, 5 min total). The crude lysate was centrifuged (45 min, 20,000 × g, 4 °C) decanting the supernatant to remove any cellular debris/pellet. The supernatant was pooled and syringe filtered (0.45 µm), to which imidazole was added (20 mM) and the supernatant was applied/loaded onto a Zn-affinity column (His-trap Chelating HP, Cytvia, 5 mL) equilibrated with 50 mM KPB, pH 7.4, and 20 mM imidazole. The column was washed with 50 mM KPB, pH 7.4, 0.3 M NaCl, 20 mM Imidazole (1 CV), then 50 mM imidazole (5 CV), and the protein was eluted in bulk with buffer consisting of 50 mM KPB, pH 7.4, 0.3 M NaCl, and 80 mM imidazole (10 CV) and 150 mM Imidazole (3 CV). Fractions containing protein as confirmed by SDS-PAGE were then pooled and subjected to a TEV digest, with dialysis at 4 °C overnight (50 mM KPB, pH 7.4). The dialyzed sample was incubated at 50 °C for 15 min to quench the TEV digest. The supernatant was clarified via centrifugation (10,000 × g, 10 min, 4 °C), then loaded onto a pre-equilibrated HiLoad 16/600 Superdex 200 pg column (Cytvia) (2 CV of SEC Buffer: 25 mM Tris, pH 7.4, 0.1 M KCl, 1 mM TCEP), yielding the desired protein (∼95 kDa). The protein was found to elute as a monomer, with minimal to no aggregation visible. Fractions containing the desired protein, as judged by SDS-PAGE, were pooled and concentrated via centrifugation in SEC buffer to yield purified apo-*t*MS^HcyFolCapCobmut1^(∼15 mg/mL). To the purified apo protein (∼150 μM) was added propylcobalamin (prCbl, ∼170 μM). The resulting mixture was loaded onto a pre-equilibrated desalting column (PD-10, GE/Cytvia, SEC buffer) to remove excess prCbl and concentrated via centrifugation in SEC buffer to yield purified holo*-t*MS^HcyFolCapCobmut1•prCbl^ (∼15 mg/mL), which was stored at 4 °C or flash-frozen for long-term storage at −80 °C.

#### *t*MS^HcyFolCapCobmut2•aeCbl^ (Hcy-on)

Purification of *t*MS^HcyFolCapCobmut2•aeCbl^ was conducted similarly, with no added cobalamin. Protein purification yielded purified apo*-t*MS^HcyFolCapCobmut2^ (∼20 mg/mL); to the purified apo protein (∼150 μM) was added aminoethylcobalamin (aeCbl, ∼170 μM). The resulting mixture was loaded onto a pre-equilibrated desalting column (PD-10, GE/Cytvia, 25 mM Tris, pH 7.4, 0.1 M KCl, 1 mM TCEP) to remove excess aeCbl and concentrated via centrifugation in 25 mM Tris, pH 7.4, 0.1 M KCl, 1 mM TCEP to yield purified holo-*t*MS^HcyFolCapCobmut2•aeCbl^ (∼18 mg/mL), which was stored at 4 °C or flash-frozen for long-term storage at −80 °C.

### *t*MS Qualitative UV-Vis Assay

The methylcobalamain:homocysteine methyltransferase reaction was monitored in the presence or absence of *t*MS constructs spectrophotometrically in the dark, tracking the time-dependent spectral changes of methylcobalamin (MeCbl) in the presence of homocysteine at 50 °C for up to 10 minutes. Briefly, the reaction mixture (1 mL total volume) was carried out under aerobic conditions containing either free MeCbl (20 µM) or protein-bound MeCbl (*t*MS^CapCob•MeCbl^, 23 µM) and homocysteine (100 µM) in 50 mM KPB, pH 7.4, 0.5 mM TCEP using *t*MS if applicable (tMS^ΔN35Hcy^ or *t*MS^HcyFol^, 2 µM). Reactions were incubated with the reaction mixture in the absence of homocysteine at 50 °C, then initiated by adding homocysteine. Time-dependent spectral changes were recorded were recorded for 0-10 min using a Cary 300 Bio UV-Vis spectrophotometer (Varian, Inc.)

### Crystallization of *t*MS Constructs

Crystals were grown via sitting drop vapor diffusion. *t*MS^FolCapCobD759A•MeCbl^ (∼10 mg/mL in 25 mM KPB, pH 7.4) was mixed with a reservoir solution containing 25% PEG 3350, 0.1 M Tris (pH 8.5), and 0.2 M ammonium acetate in a 1:1 ratio (0.4 µL each) and incubated at 4 °C. Crystals were briefly transferred to a cryoprotectant solution containing approximately 20% glycerol and 20% PEG 3350, 0.08 M Tris (pH 8.5), and 0.16 M ammonium acetate for 30 seconds prior to harvesting and flash freezing in liquid nitrogen.

*t*MS^FolCapCobD762G•cpCbl^ (∼10 mg/mL in 25 mM KPB, pH 7.4) was mixed with a reservoir solution containing 1.26 M (NH_4_)_2_SO_4_, 0.1 M CHES-Na (pH 9.5), and 0.2 M NaCl in a 1:1 ratio (0.4 µL each) and incubated at 20 °C. Crystals were briefly transferred to a cryoprotectant solution containing approximately 20% glycerol and 1 M (NH_4_)_2_SO_4_, 0.08 M CHES-Na (pH 9.5), and 0.16 M NaCl for 3 minutes prior to harvesting and flash freezing in liquid nitrogen.

*t*MS^HcyFolCapCobmut1•prCbl^ (∼15 mg/mL in 25 mM Tris, pH 7.5, 50 mM KCl, 1 mM TCEP) was mixed with a reservoir solution containing 30% PEG 3350, 0.1 M Bis-Tris (pH 6.5), and 0.2 M NaCl in a 1:1 ratio (0.4 µL each) and incubated at 20 °C. Crystals were briefly transferred to a cryoprotectant solution containing approximately 20% glycerol and 24% PEG 3350, 0.08 M Bis-Tris (pH 6.5), and 0.16 M NaCl for ∼30 seconds prior to harvesting and flash freezing in liquid nitrogen.

*t*MS^HcyFolCapCobmut2•aeCbl^ (∼18 mg/mL in 25 mM Tris, pH 7.5, 50 mM KCl, 1 mM TCEP) was mixed with a reservoir solution containing 20% PEG 3350, 0.1 M Tris (pH 8.5), and 0.2 M NaCl in a 1:1 ratio (0.4 µL each) and incubated at 4 °C. Crystals were briefly harvested and flash frozen in liquid nitrogen.

Data collection and processing statistics are summarized in Supplementary Table 1. Data for *t*MS^FolCapCobD759A•MeCbl^ were indexed to spacegroup P4_1_2_1_2 (unit-cell parameters a = b = 187.77, c = 325.55 Å) with six molecules in the asymmetric unit (Matthew’s coefficient VM = 4.22 Å^3^ Da^−1^, 71% solvent content). Data for *t*MS^FolCapCobD762G•cpCbl^ were indexed to spacegroup P6_1_22 (unit-cell parameters a = b = 95.15, c = 274.29 Å) with one molecule in the asymmetric unit (Matthew’s coefficient VM = 3.17 Å^3^ Da^−1^, 61% solvent content). Data for *t*MS^HcyFolCapCobmut1•prCbl^ were indexed to spacegroup P2_1_2_1_2_1_ (unit-cell parameters a = 69.95, b = 83.82, c = 163.72 Å) with one molecule in the asymmetric unit (Matthew’s coefficient VM = 2.51 Å^3^ Da^−1^, 51% solvent content). Data for *t*MS^HcyFolCapCobmut2•aeCbl^ were indexed to spacegroup P2_1_2_1_2_1_ (unit-cell parameters a = 70.81, b = 115.10, c = 120.80 Å) with one molecule in the asymmetric unit (Matthew’s coefficient VM = 2.58 Å^3^ Da^−1^, 52% solvent content).

### Data Collection and Refinement

X-ray data sets were collected at 100 K on GM/CA beamline 23-ID-B at the Advanced Photon Source, Argonne National Laboratory (Argonne, IL) for *t*MS^FolCapCobD759A•MeCbl^ and *t*MS^FolCapCobD762G•cpCbl^, on GM/CA beamline 23-ID-D for *t*MS^HcyFolCapCobmut2•aeCbl^, and on LS-CAT beamline 21-ID-D at the Advanced Photon Source, Argonne National Laboratory (Argonne, IL) for *t*MS^HcyFolCapCobmut1•prCbl^. Data sets were processed using xia2/DIALS^46^. Initial phases were obtained using MOLREP^47^. Individual domains of the full-length *t*MS structure (8SSC) were used as search models. Iterative model building and corrections were performed manually using Coot^48^ following molecular replacement and subsequent structure refinement was performed with CCP4 Refmac5^49^. Initial refinement was conducted using BUSTER^50^ to rapidly fix Ramachandran, rotamer, and density fit outliers, refining to convergence and adding waters in the final automated round of refinement.

Phenix eLBOW^51^ was used to generate the initial ligand restraints using ligand ID “COB” or “B_12_”. To perform ligand fitting of structures containing non-native cobalamins, the process was as follows. Briefly, B_12_ was used to model the cofactor as Cob(II) in the case of all structures save for 9CBO (MeCbl holoenzyme), for which COB (methylcobalamin) was used. Ligand fitting was done towards the tail end of the structural refinement, to minimize any potential bias/overfitting. Phenix LigandFit^52^ was used to provide initial fits and placements of the ligands, followed by iterative refinement automated refinement using BUSTER^50^ and PDBREDO^53^ and manual corrections via Coot^48^; CCP4 Refmac5^49^ was used once the automated ligand workflows converged on an optimized ligand placement/geometry, using a manually edited B_12_ ligand restraint file that was initially obtained via minimization in Phenix eLBOW^51^.

Manual inspection of the resulting maps (2Fo-Fc) and composite omit electron density maps (Fo-Fc) as calculated by Phenix Composite_omit_map)^54^ were used to verify all models, and the structures containing non-native cobalamins (9CBP, pre-Hcy-on, prCbl; 9CBQ, Fol-on, cpCbl; 9CBR, Hcy-on, aeCbl) showed clear, visible, residual, unmodelled electron density in the upper axial position (β-face) of cobalamin. The corresponding upper axial ligands were added manually, followed by automated refinement (BUSTER, PDBREDO)^50,53^, with a final round of refinement conducted using CCP4 Refmac5^49^ including the respective upper axial ligand’s restraint file.

PDB-REDO^53^ was used to assess the model quality in between refinements and to fix any rotamer and density fit outliers automatically. The model quality was evaluated using MolProbity^55^. The Hcy-on structure data set showed partial crystal twinning and final rounds of refinement were conducted using twin refinement, as suggested by PDB-REDO. Figures showing crystal structures were generated in PyMOL^56^.

### Statistical Analysis and Reproducibility

Unless otherwise stated, functional assays were conducted using n = 2 independent replicates. At least three independent experiments were conducted for each functional assay. All attempts at replication were successful. Analysis and curve-fitting was performed using Prism 10.2.3.

## Supporting information

Suppplementary Information

## Data Availability

The structure coordinates and structure factors reported in this study have been deposited in the Protein Data Bank under accession codes 9CBO (*t*MS^FolCapCobD759A•MeCbl^), 9CBP (*t*MS^HcyFolCapCobmut1•prCbl^), 9CBQ (*t*MS^FolCapCobD762G•cpCbl^), and 9CBR (*t*MS^HcyFolCapCobmut2•aeCbl^). PDB codes of previously published structures used in this study are 8SSC, 8SSD, 8SSE, and 8G3H. All other data are available from the corresponding authors upon request. Source data are provided with this paper.

## Corresponding author

Correspondence should be addressed to Kazuhiro Yamada and Markos Koutmos.

## Competing Interests

The authors declare no competing interests.

## Notes

### Competing Interest Statement

The authors have declared no competing interest.

### Summary of Updates

Revised main text, associated figures, and supplemental information. Revised deposition for all structures and associated analysis.

### Funding

- U.S. National Science Foundation: 1945174

