## Supplementary material for "Structural Snapshots of B_12_-Dependent Methionine Synthase’s Catalytic Conformations": Suppplementary Information

### Supplementary Information

Supplementary Figures

1. Captured Methionine Synthase Cap-on states demonstrate the flexibility of FolCap linker (Linker II)
2. Captured Methionine Synthase Cap-on states are gateway conformations
3. Cap-off MS structures highlight the preorganized/conserved nature of the Cob and Cap domains
4. Fol-on conformation represents a 'right-after-catalysis' state
5. Methionine Synthase gateway states are *en route* to catalysis
6. Folate activation mechanism
7. *t*MS electron density around the cobalamin cofactor
8. Composite Polder omit map of non-native cobalamin cofactors used
9. Limited Proteolysis of *t*MS$^{HcyFolCapCob}$ in its wildtype form and in combination with different cobalamins
10. Mutations introduced to *t*MS$^{HcyFolCapCob}$ to aid in crystallization
11. Hcy-on conformation demonstrates the role of five coordinate Cbl in catalysis
12. Methionine Synthase cofactor changes associated with the pre-Hcy-on to Hcy-on transition
13. Homocysteine methylation mechanism confirms Zn inversion/elasticity
14. The role of the Fol domain in guiding catalytic transitions
15. Methionine Synthase (MS) conformational ensemble as determined from this study
16. Structural alignment of the domains in the pre-catalytic and catalytic states of *t*MS
17. Unified models for MS catalysis provide a predictive framework for focused hypothesis testing
18. Expanded MS catalytic model including conformation ensemble and potential transitions
19. Substrate binding sites and electron density in *t*MS, pre-catalytic and catalytic states
20. Methionine synthase (MS) amino acid alignment
21. Weblogo representation of a multiple sequence alignment of Methionine Synthase (MS) (n=1983)
22. Non-native cobalamins used to capture catalytic conformations compete with substrate binding
23. Structural elements that can potentially gate the transition between the pre-Hcy-on and Hcy-on states
24. *t*MS$^{HcyFolCapCob}$ copurifies with tetrahydrofolate (THF) in both the pre-Hcy-on and Hcy-on states
25. Corrin ring distortion analysis reveals changes associated with redox and ligation status
26. Corrin ring overlay of captured states reveals changes associated with redox and ligation status

Supplementary Tables

Table 1. X-Ray Data Collection and Refinement Statistics
Table 2. Bacterial strains, plasmids, and synthetic oligonucleotides used in this study

### Supplementary Figures

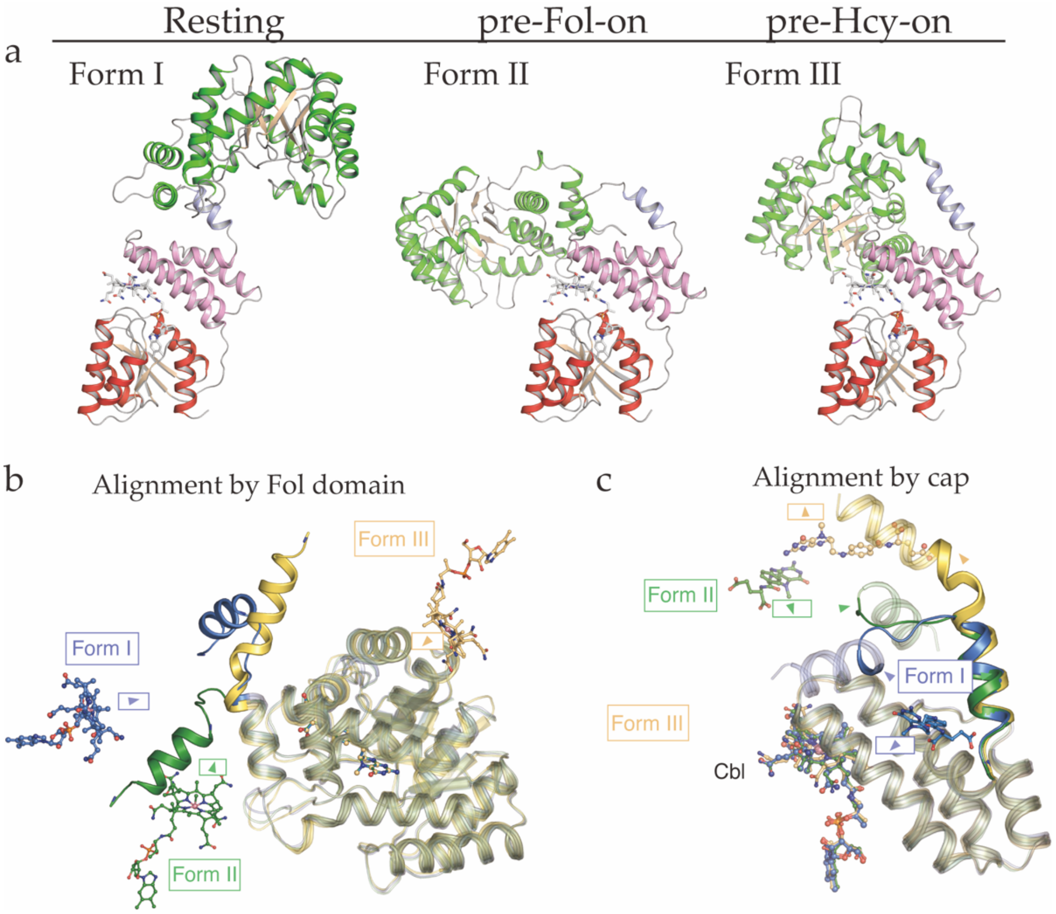


**Supplementary Figure 1. Captured Methionine Synthase Cap-on states demonstrate the flexibility of FolCap linker (Linker II). a** Cap-on states captured in this paper (Forms I and II, 9CBO; Form III, 9CBP). All structures are aligned using the Cob domain as reference and color-coded by domain and the FolCap linker (Linker II, slate; Fol, green; Cap, pink; Cob, red). Note, the main difference is with regards to the helicity/structuring of the Fol:Cap linker, being most-structured in the pre-Hcy-on (Form III) state and undergoing a helix-loop transition that shrinks the helical linker by 6 amino acids in the pre-Fol-on (Form II) state and similarly in the 'resting' state. Note, **b** depicts the same structures aligned using the Foll domain as reference, demonstrates the Fol domain active site and the movement that must accompany its transition to face the Cob domain. Notably, the pre-Fol-on state has the Fol domain and its active site primed for ternary complex formation with the Cob domain and its bound cobalamin cofactor (Form II, forest green); the pre-Hcy-on structure has the Fol domain and its active site completely facing away from the Cob domain, though the Hcy domain is primed to form a ternary complex with the Cob domain (Form III, golden yellow). The 'resting' state structure has the Fol domain facing the Cob domain but would require a much larger translation to interact with the Cob domain (Form I, navy blue). **c** Similarly, when the structures are aligned using the Cap domain as reference, the orientation of manually-docked methyltetrahydrofolate (MTF) reveals that the pre-Fol-on state has the methyl group of MTF oriented towards cobalamin, while the pre-Hcy-on state has the methyl group of MTF pointed away from cobalamin. Though the 'resting' state has the methyl group pointed towards the cobalamin domain, the uncapping motion associated with Cap-on to Cap-off transitions would occlude its interaction, while allowing for the pre-Fol-on bound MTF to approach cobalamin from its upper axial face.

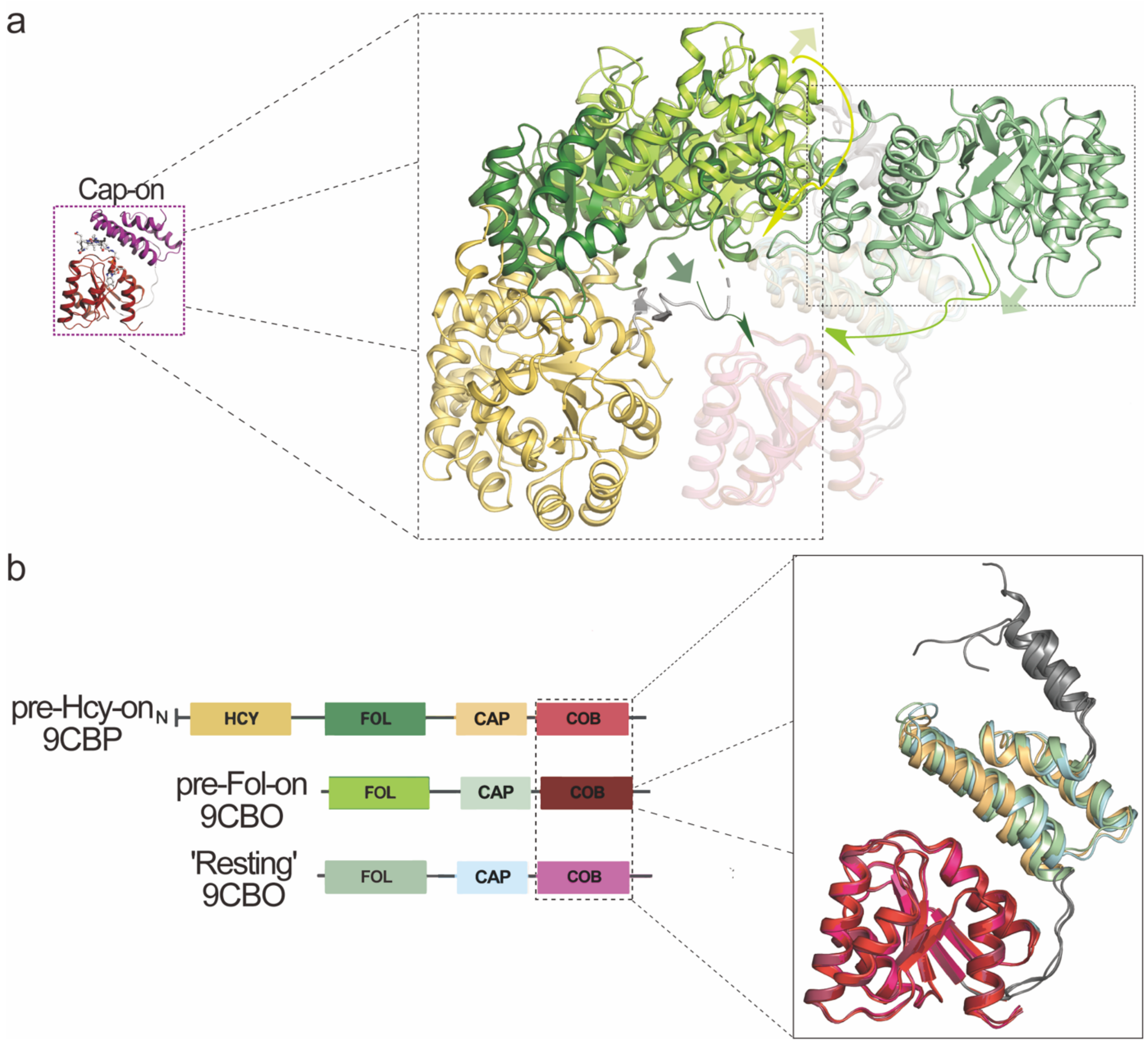


**Supplementary Figure 2. Captured Methionine Synthase Cap-on states are gateway conformations.** Cap-on states captured in this paper. All structures are aligned using the Cob domain as reference and color-coded according to the scheme shown in **b**. Note, the main difference is with regards to the helicity/structuring of the Fol:Cap linker, being most-structured in the pre-Hcy-on state and undergoing a helix-loop transition that shrinks the helical linker by 6 amino acids in the pre-Fol-on state and similarly in the 'resting' state. Note, **a** demonstrates the Fol domain active site and the movement that must accompany its transition to face the Cob domain. Notably, the Pre-Fol-on state has the Fol domain and its active site primed for ternary complex formation with the Cob domain; the pre-Hcy-on structure has the Fol domain and its active site completely facing away from the Cob domain, though the Hcy domain is primed to form a ternary complex with the Cob domain. The 'resting' structure has the Fol domain facing towards the Cob domain, but would require a much larger translation to interact with the Cob domain.

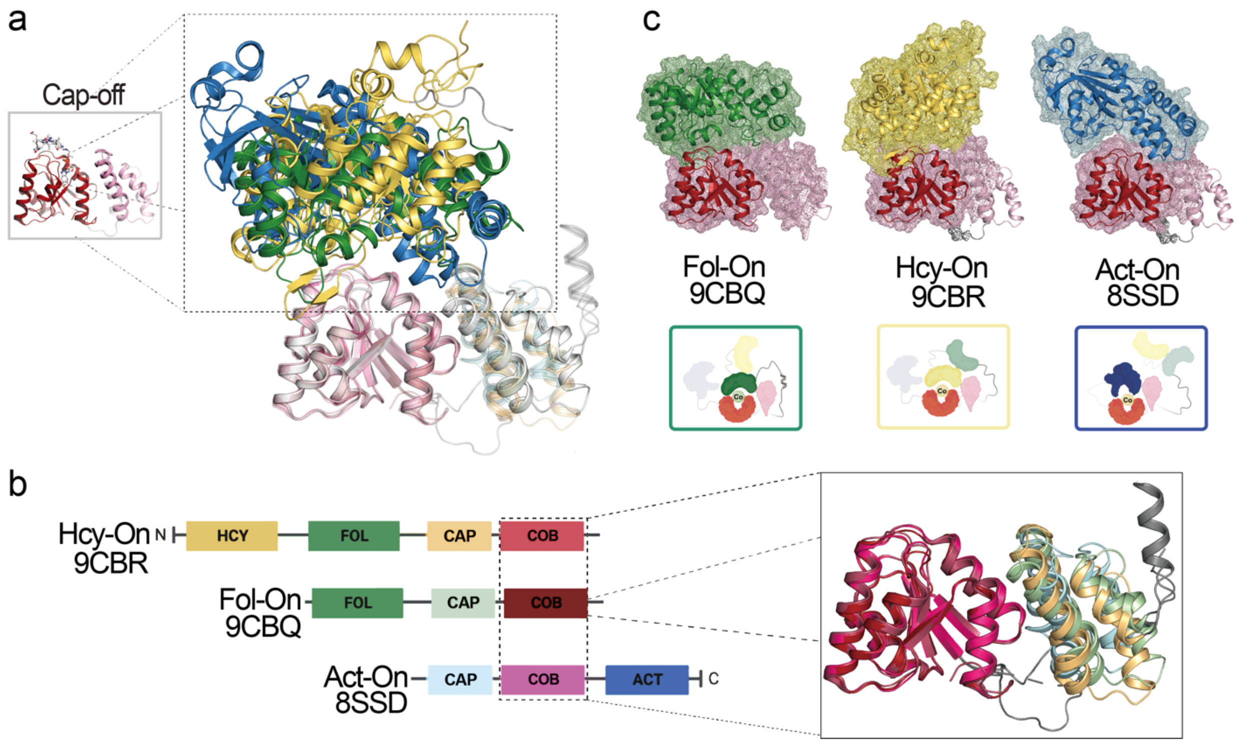


**Supplementary Figure 3. Cap-off MS structures highlight the preorganized/conserved nature of the Cob and Cap domains.** Cap-off states captured in this paper as compared to a previously captured Act-on structure (8SSD). All structures are aligned using the Cob domain as reference and color-coded according to the scheme shown in **b**. Note, the main difference is with regards to the helicity/structuring of the Fol:Cap linker, being most-structured in the Fol-on state and undergoing a helix-loop transition that unstructured the helical linker in the Hcy-on state and similarly in the Act-on state. Note, **a** demonstrates that the associated substrate binding domain forms a ternary complex with the Cob domain, with the associated surface renderings highlighting a novel ternary active site formed depending on the reactive state/conformation (**b**). The Cap domain adopts a similar position relative to the Cob domain, and the associated Cap:Cob linker is more flexible as compared to the gateway conformations.

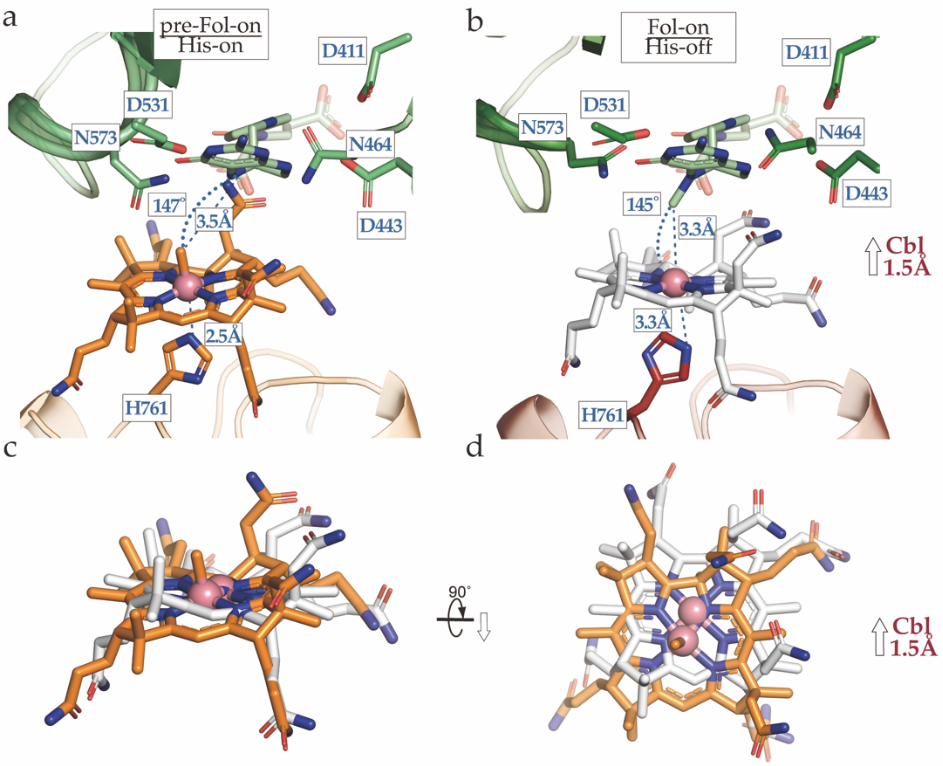


**Supplementary Figure 4. Fol-on conformation represents a 'right-after-catalysis' state.** Folate was docked manually by aligning the Fol domain of 9CBP (pre-Hcy-on) (this paper) to the Fol domain of the pre-Fol-on (9CBO, Cbl, orange) and Fol-on model (9CBQ, Cbl, light grey). **a** Folate docked model of pre-Fol-on state (9CBO) and **b** Fol-on state (9CBQ). Note the increased His761 distance in the Fol-on state (~Δ0.9 Å) and lateral shift of the cobalt center (**c, d**). The Cbl cofactor is shifted laterally 1.5 Å, and the corrin ring is more planar in the Fol-on vs. pre-Fol-on state.

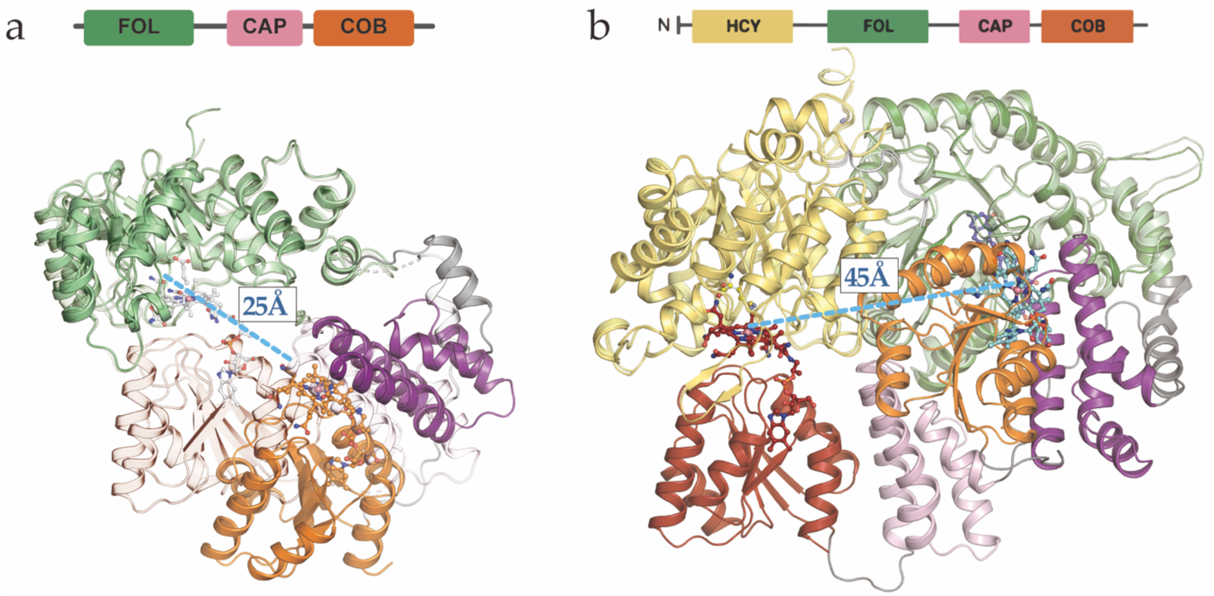


**Supplementary Figure 5. Methionine Synthase gateway states are *en route* to catalysis. a** Alignment of pre-Hcy on (9CBP) and pre-Fol-on (9CBO) states with respect to the Cob domain using the coloring scheme from Figures **2** and **3**. The Hcy domain is not shown for simplicity. Note that the Cap domain remains essentially unchanged. Transition between the pre-Hcy-on to pre-Fol-on state would require coupled helix-loop transition of the Fol:Cap linker (Linker II), which would amount to a 18 Å translation of Linker II (**b)** and 120° rotation of the Fol domain (**c**). **d** pre-Fol-on (9CBO) and Fol-on (9CBQ) states aligned using the Fol domain as reference. The Cob domain must traverse 25 Å following 'uncapping' to form a ternary complex with the Fol domain. **e** pre-Hcy-on (9CBP) and Hcy–on (9CBR) states aligned using the Fol domain as reference. The Cob domain must traverse 45 Å following 'uncapping' to form a ternary complex with the Hcy domain.

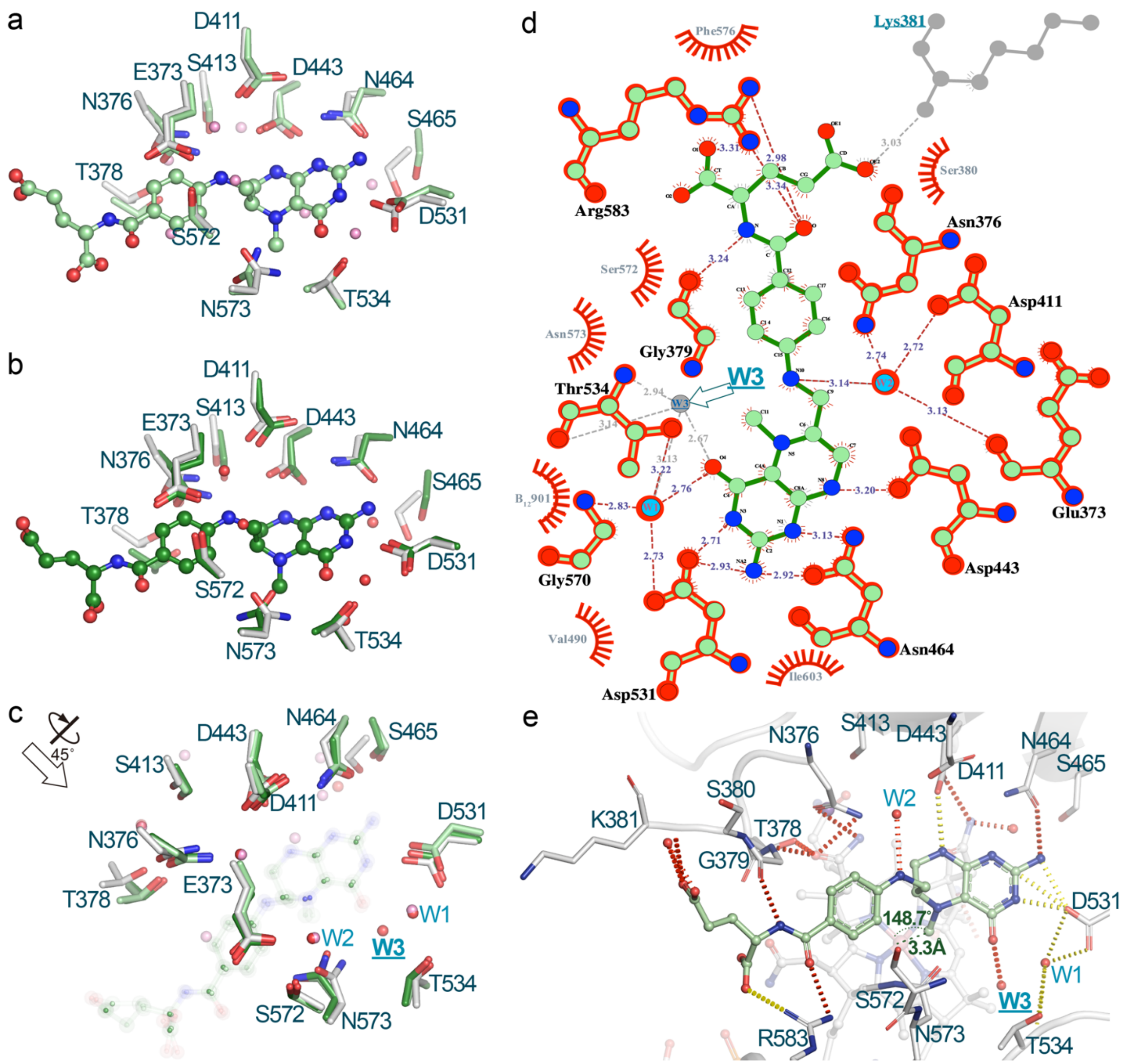


**Supplementary Figure 6. Folate activation mechanism.** Folate was docked manually by aligning the Fol domain of 5VOP (dark green, **b, c**) and 9CBP (light green, **a, c**). (this paper) to the Fol domain of the Fol-on model (light grey, 9CBQ). Water 3 (W3) coordinated via Thr534 directly interacts with O4 of the pterin ring of methyltetrahydrofolate in the manually docked folate Fol-On model (using 5VOP) (**c, d, e**). Water 1 (W1), coordinated via Asp531, Thr534, and Gly570 also interacts with O4 of the pterin ring, while Water 2 (W2), coordinated via Glu373, Asn376, Asp411 interacts with N10 of the folate moiety; these water-mediated interactions are conserved in both the pre-Hcy-on manually docked folate Fol-on model (9CBP) and the aforementioned model using 5VOP (**c, d, e**). Notably, no direct residue or water interacts with C11 or N5 of the pterin ring, indicating that direct protonation cannot occur without substantial changes in the Fol domain or that activation of N5 bearing the C11 methyl group occurs via an indirect hydrogen bonding network. The distance between C11 and the Co center is 3.3 Å, with a corresponding angle of ~148.7°, with a linear arrangement consistent with an $S_N2$ mechanism proposed for methyltetrahydrofolate demethylation by cob(I)alamin.

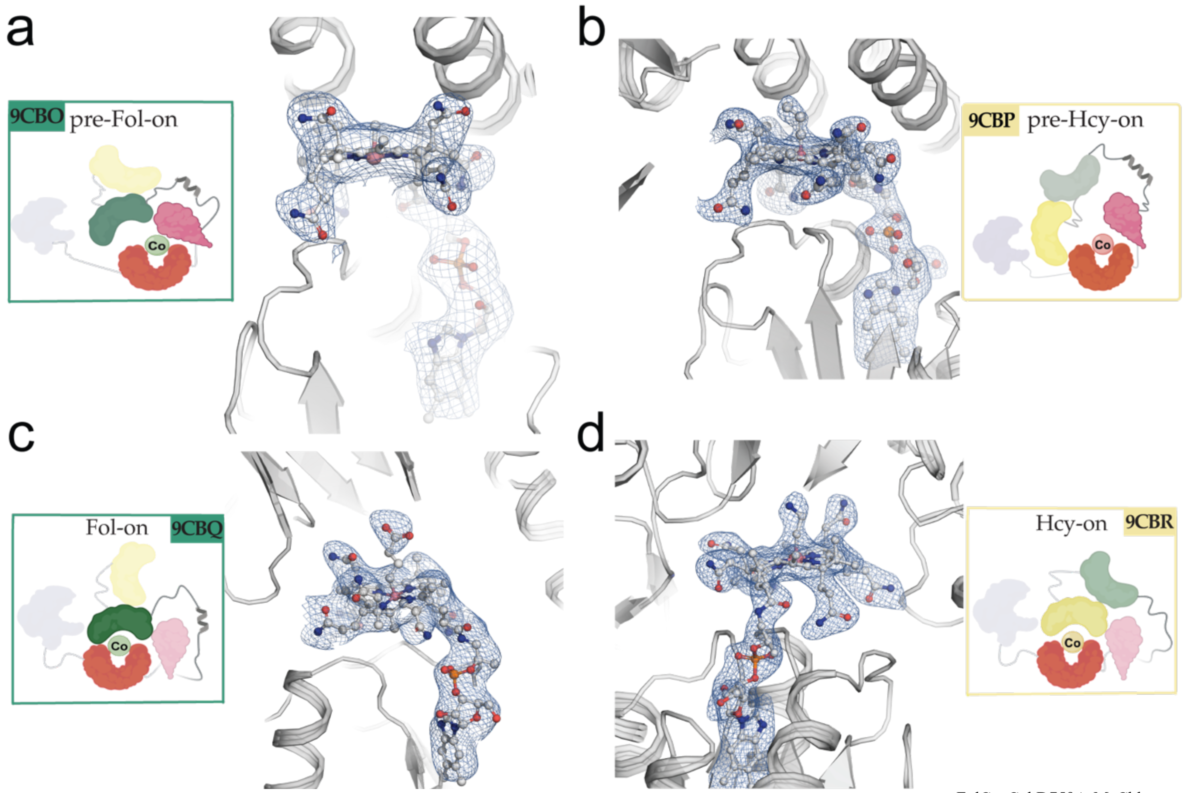


**Supplementary Figure 7. *t*MS electron density around the cobalamin cofactor. a** *t*MS$^{FolCapCobD759A•MeCbl}$ and its cobalamin cofactor (gray), along with Phe713 and His761 in the axial cofactor positions. Their corresponding electron density (2$F_o$-$F_c$) contoured at 1.5 σ are shown in blue. **b** *t*MS$^{HcyFolCapCobmut1•prCbl}$ and its cobalamin cofactor shown using the same coloring scheme as in **a**. **c** *t*MS$^{FolCapCobD762G•cpCbl}$ and its cobalamin cofactor shown using the same coloring scheme as in **a**. **d** *t*MS$^{HcyFolCapCobmut2•aeCbl}$ and its cobalamin cofactor shown using the same coloring scheme as in **a**. Cartoon insets in panels **a-d** were created with BioRender.com.

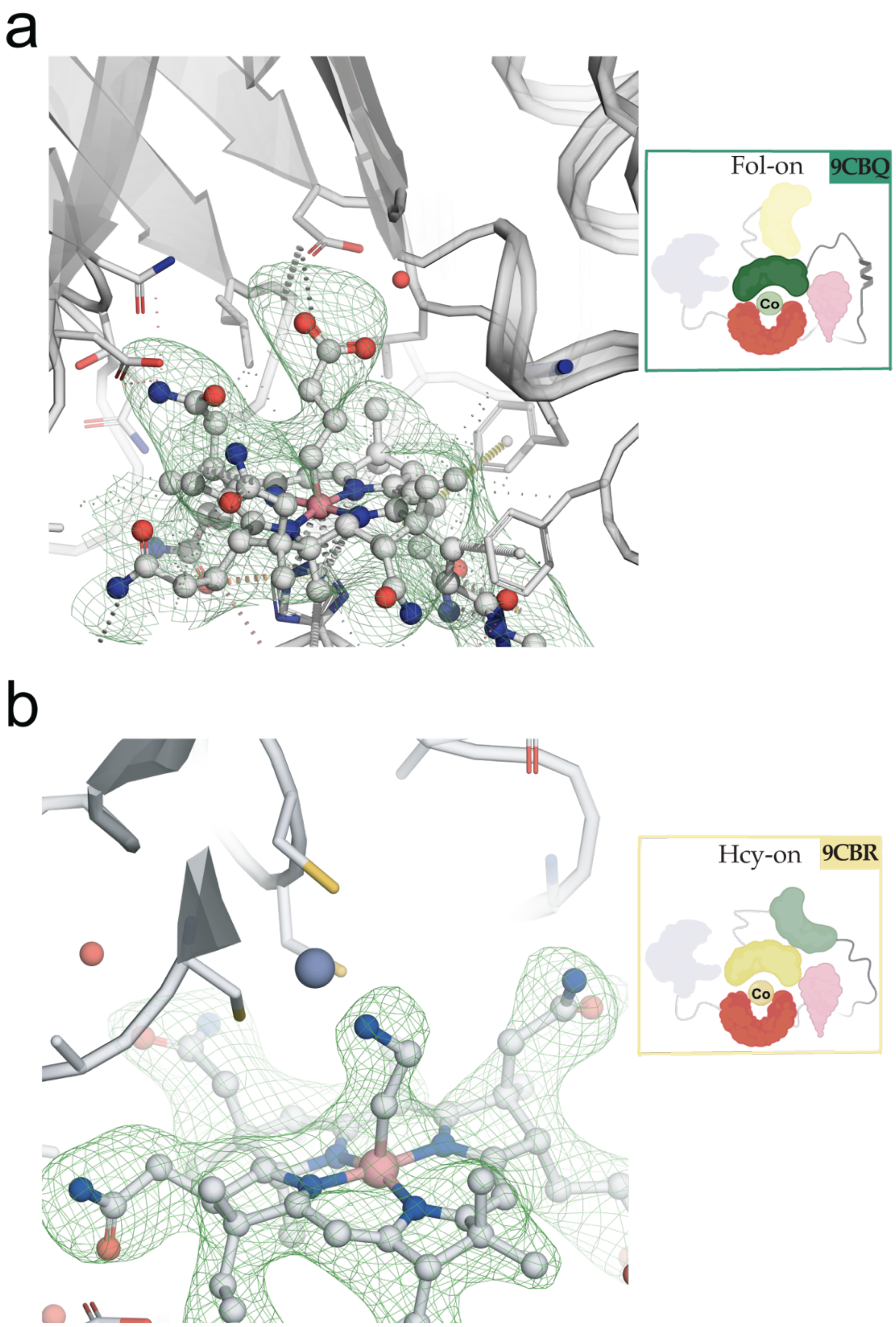


**Supplementary Figure 8. Composite Polder omit map of non-native cobalamin cofactors used. a** *t*MS$^{FolCapCobD762G\bullet cpCbl}$ and its cobalamin cofactor (gray) with its corresponding electron density from a polder omit map ($F_o$-$F_c$) contoured at 3 σ are shown in green. Note the electron density present in the upper axial portion of the non-native cobalamin cofactor, corresponding to the γ-carboxypropyl upper axial ligand. **b** *t*MS$^{HcyFolCapCobmut2\bullet aeCbl}$ and its cobalamin cofactor shown using the same coloring scheme as in **a**, with its corresponding electron density from a polder omit map ($F_o$-$F_c$) contoured at 3 σ are shown in green. Note the electron density present in the upper axial portion of the non-native cobalamin cofactor, corresponding to the aminoethyl upper axial ligand. Cartoon insets in panels **a-b** were created with BioRender.com.

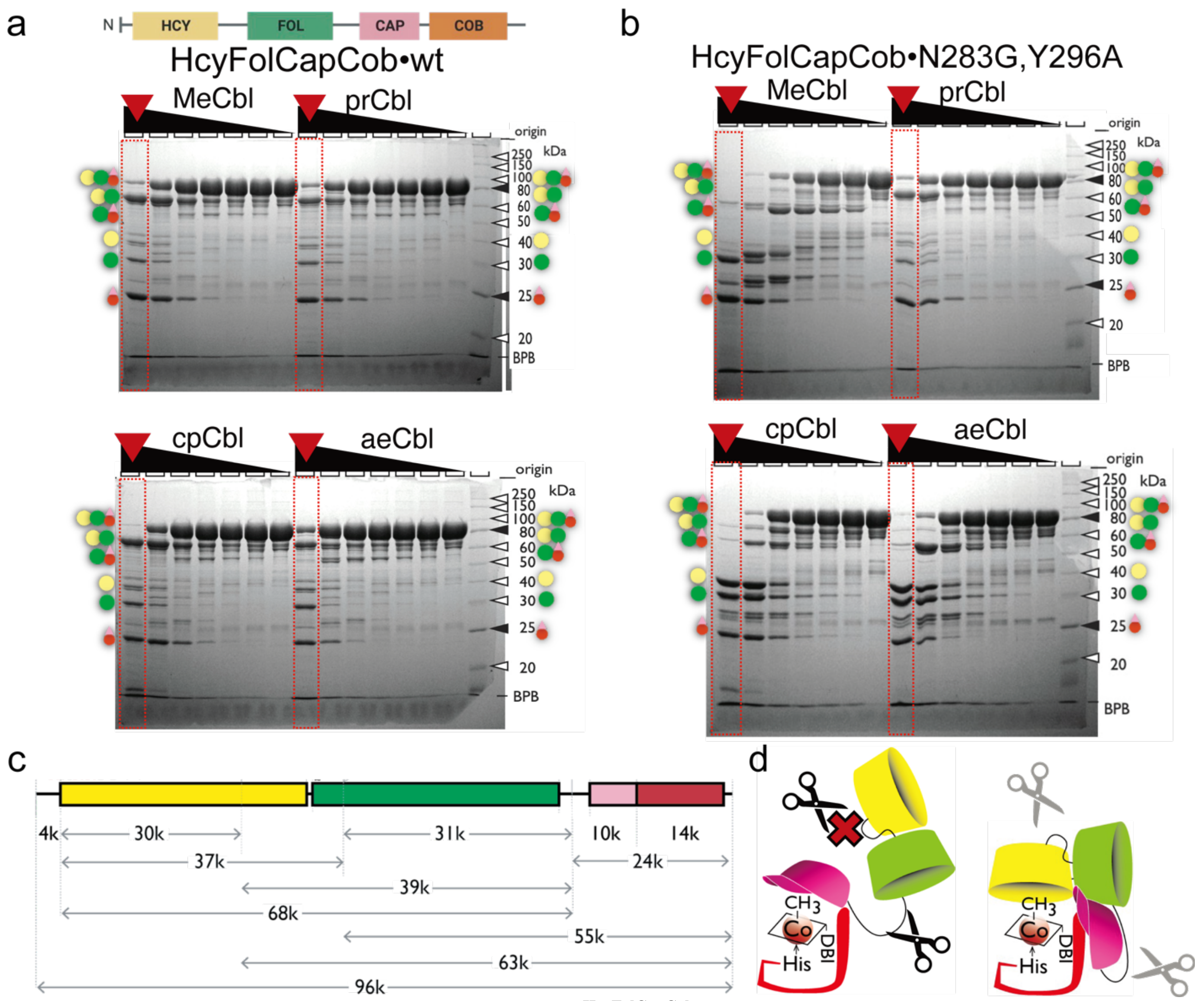


**Supplementary Figure 9. Limited Proteolysis of $t$MS$^{HcyFolCapCob}$ in its wildtype form and in combination with different cobalamins. a** A quantity of 14 µg of purified protein ($t$MS$^{HcyFolCapCob•wt}$) was incubated with varying amounts of trypsin (from ~2 µg to ~0.2 ng) for 20 min at room temperature. The reaction was carried out both in the absence as well as in the presence of 100 µM cobalamin. Trypsin activity was quenched after the reaction by the addition of sample buffer for SDS-PAGE containing 1% SDS followed by heating at 95 ˚C for 10 minutes. The resulting $t$MS$^{HcyFolCapCob•wt}$ fragments were separated by SDS-PAGE and visualized by Coomassie brilliant blue staining. **b** A mutant construct ($t$MS$^{HcyFolCapCob•N283G,Y296A}$) was similarly studied. **c** The schematic representation of the domain structure of $t$MS$^{HcyFolCapCob}$ is shown, along with the theoretical molecular weight of each domain. The Hcy domain (yellow), Fol domain (green), the Cap domain (pink), and the Cob domain (red) accommodate the homocysteine substrate, methyltetrahydrofolate substrate, protect the cobalamin cofactor and bind the cobalamin cofactor, respectively. **d** The protein conformations of $t$MS$^{HcyFolCapCob}$ in the Cap-on, pre-Hcy-on and Cap-off, Hcy-on states are shown in cartoon mode. The color scheme is the same as in panel **c**. In the Cap-on state, the FolCap linker is exposed to the solvent, facilitating trypsin access to the solvent-exposed retractable region, and allowing cleavage of the linker region. In contrast, protease access to the HcyFol linker is difficult when in the Cap-on state, while it is more accessible in the Cap-off state. Cartoon insets in panels **c-d** were created with BioRender.com.

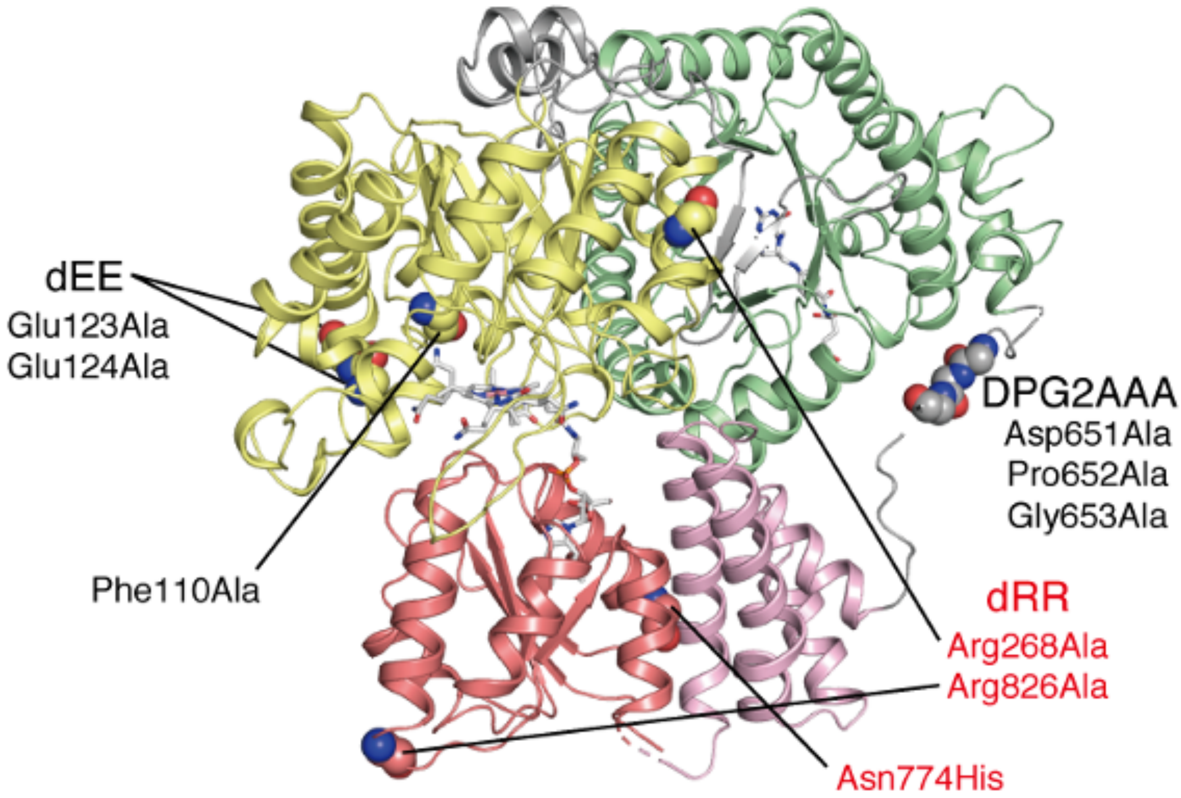


**Supplementary Figure 10. Mutations introduced to *t*MS$^{HcyFolCapCob}$ to aid in crystallization.** The structure of *t*MS$^{HcyFolCapCob}$ in the Hcy-on mode is depicted in the ribbon diagram, colored using yellow, green, pink, and brown used to denote the Hcy, Fol, Cap, and Cob domains, respectively. Gray is used to denote interdomain linkers. Glu123Ala and Glu124Ala substitutions (dEE) are incorporated to enhance the crystal contact through reduction in surface energy[1]. Furthermore, mutations Asp651Ala, Pro652Ala, and Gly653Ala (denoted as DPG2AAA) lie in the Fol:Cap linker (Linker II) and were introduced to promote the formation of a helix, favoring the pre-Hcy-on conformation over the pre-Fol-on conformation. The side chain of Phe110, proximate to the homocysteine-binding site, is replaced with Ala to reduce steric clashes that would allow it to accommodate the bulky upper ligand of cobalamin analogs. All crystals of *t*MS$^{HcyFolCapCob}$ in pre-Hcy-on and Hcy-on modes contain those mutations, represented by black letters. Crystals of *t*MS in the Hcy-on conformation were obtained with additional mutations indicated in red, such as Arg268Ala and Arg826Ala (dRR), along with the Asn774His mutation. The dRR mutations are expected to attenuate the Hcy:Cob domain interaction observed in the Cap-on pre-Hcy-on state; the His residue possesses a slightly bulkier side chain compared to Asn, and the Asn774His mutant is presumed to lead to steric clashes with nearby amino acid side chains, thereby destabilizing the Cap-on conformation. The Hcy-on structure has an additional Asp759Ala mutation, which interacts with His761 as part of the ligating triad (Ser812 being the third residue), in addition to the mutations shown in black letters. This construct is designed to preferentially bind cobalamin in the five-coordinate, His-off state by weakening the His-Cbl interaction.

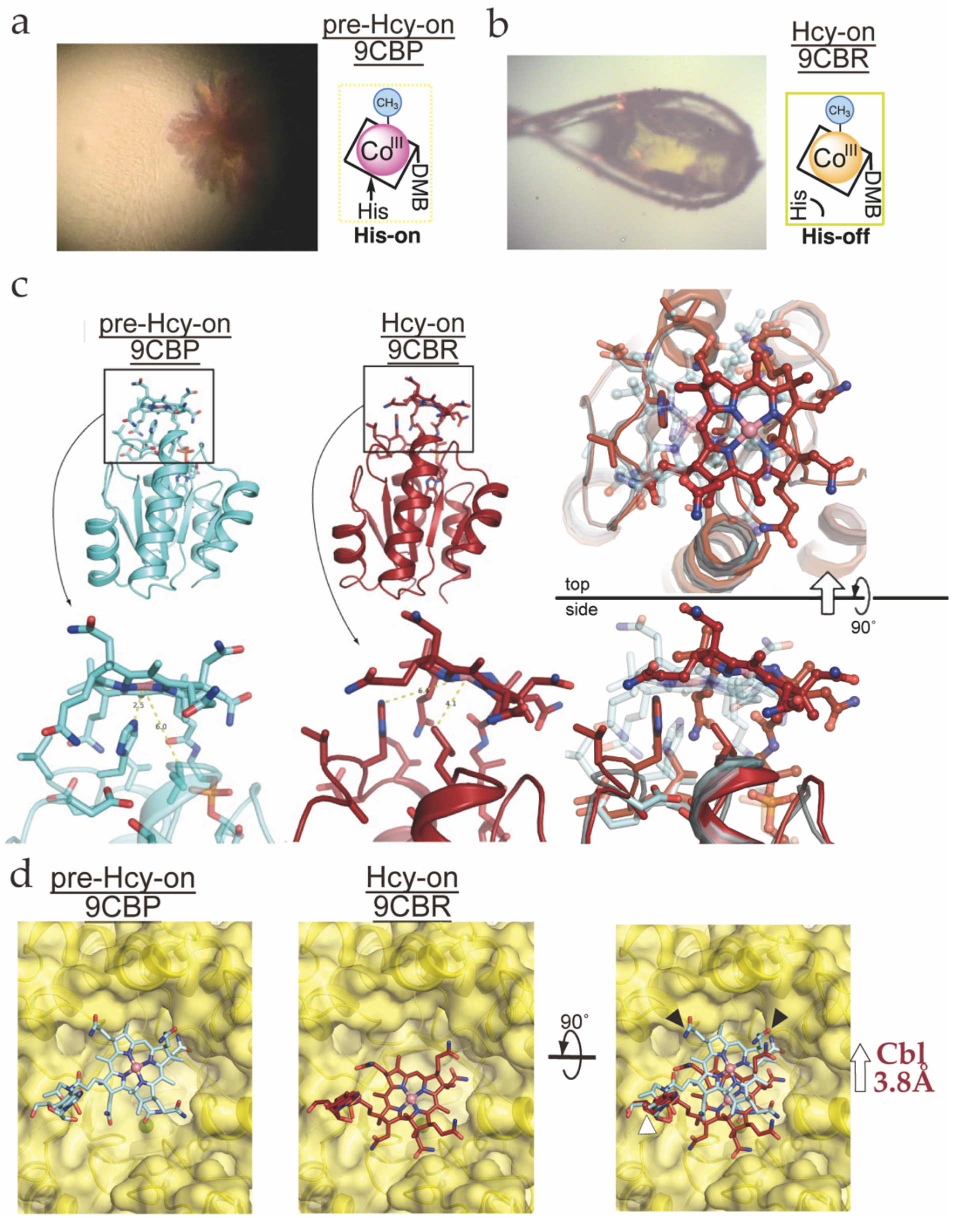


**Supplementary Figure 11. Hcy-on conformation demonstrates the role of five coordinate Cbl in catalysis. a** Color of the crystal which captured the pre-Hcy-on (9CBP) conformation versus **b** that of the crystal used to capture the Hcy-on (9CBQ) conformation. Note the change in color from red to yellow associated with the His-on to His-off ligation associated with the Co(III) state. Superposition of those structures are illustrated in **c** and **d** using the Cob domain was used as reference. **c** The His761 distance between pre-catalytic (light cyan, pre-Hcy-on) and catalytic states in *t*MS$^{HcyFolCapCob}$ (brick red, Hcy-on). **d** Surface depiction of the Hcy domain (yellow) using the same colors as **c**. Note that the DMB tail of Cbl remains unchanged between states (white arrows); black arrows indicate the steric clashes observed with the corrin ring of the pre-Hcy-on state. The Hcy-on state and its associated His-off ligation allows for the Cbl cofactor to move 3.8 Å laterally into the Hcy domain and its active site with an associated 4.2 Å tilt, relieving the steric clashes.

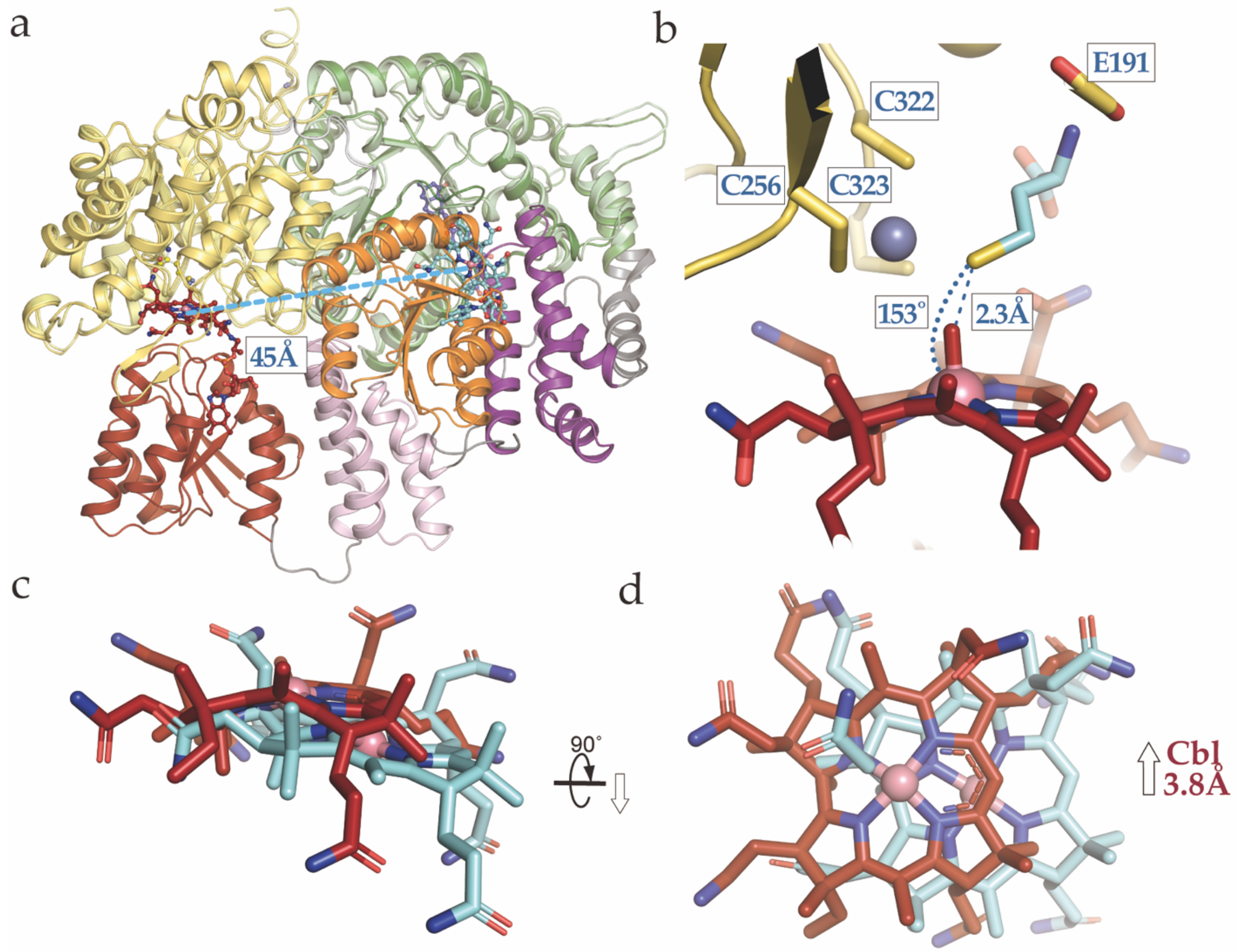


**Supplementary Figure 12. Methionine Synthase cofactor changes associated with the pre-Hcy-on to Hcy-on transition. a** pre-Hcy-on (9CBP) and Hcy-on (9CBR) states aligned using the Fol domain as reference color coded according to Fig. 3. The Cob domain must traverse 45 Å following 'uncapping' to form a ternary complex with the Hcy domain. **b** Homocysteine was docked manually by aligning the Hcy domain of 9CBP and 9CBR (yellow) using the Hcy domain as reference. **c, d** pre-Hcy-on (9CBP, cyan) and Hcy-on (9CBR, brick red) Cbl corrin ring comparison. Note the lateral shift of the cobalt center. The Cbl cofactor is shifted laterally 3.8 Å with an associated tilt of ~18°, and the corrin ring is more planar in the Hcy-on vs. pre-Hcy-on state.

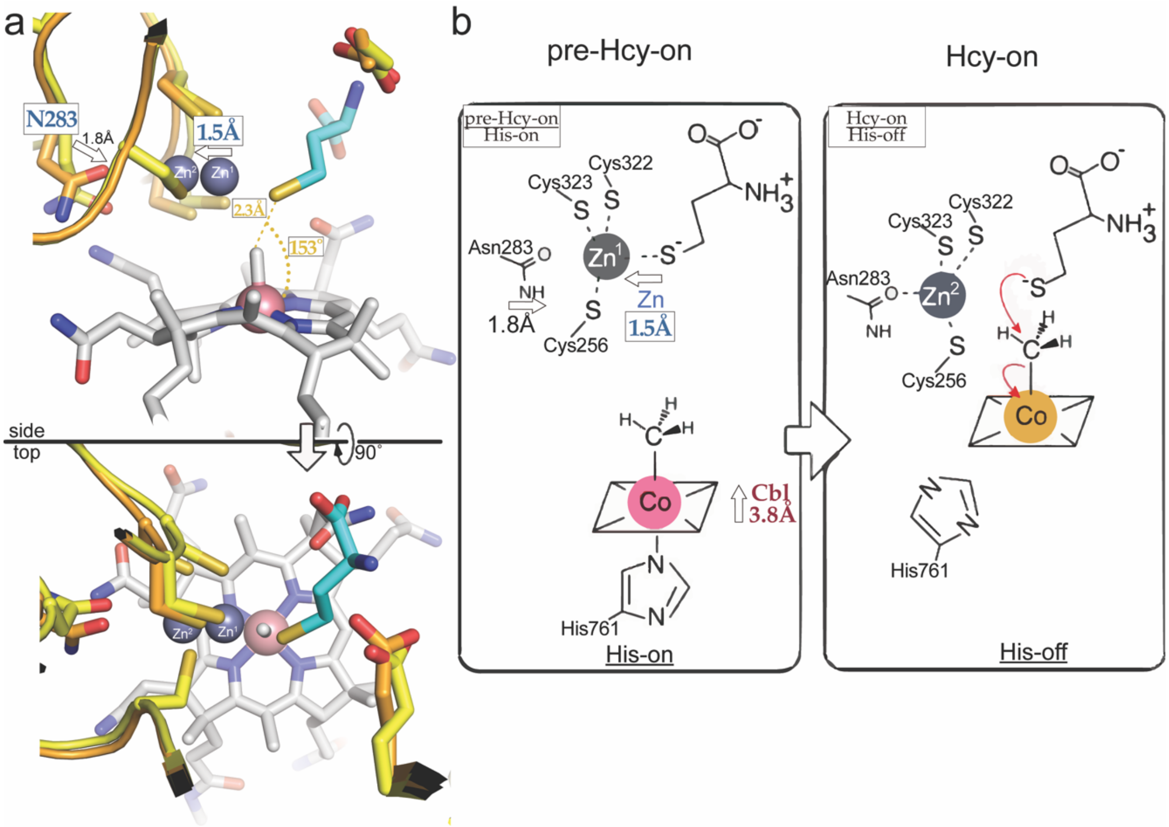


**Supplementary Figure 13. Homocysteine methylation mechanism confirms Zn inversion/elasticity.** Homocysteine was docked manually by aligning the Hcy domain of 9CBP (orange) and 9CBR (yellow) using the Hcy domain as reference. Consistent with a previously proposed zinc inversion mechanism, the Zn ion shifts 1.5 Å away from the thiol, aided by a new coordinating interaction with Asn283. This Zn "inversion" is coupled with a His-on to His-off ligation between the pre-Hcy-on and Hcy-on states, which simultaneously yields a five coordinate Cbl that is laterally shifted 3.8 Å towards the Hcy domain. The distance between the thiol and the Co center is 2.3 Å, with a corresponding angle of ~155.2°, with a linear arrangement consistent with an $S_N2$ mechanism proposed for homocysteine methylation by methylcobalamin.

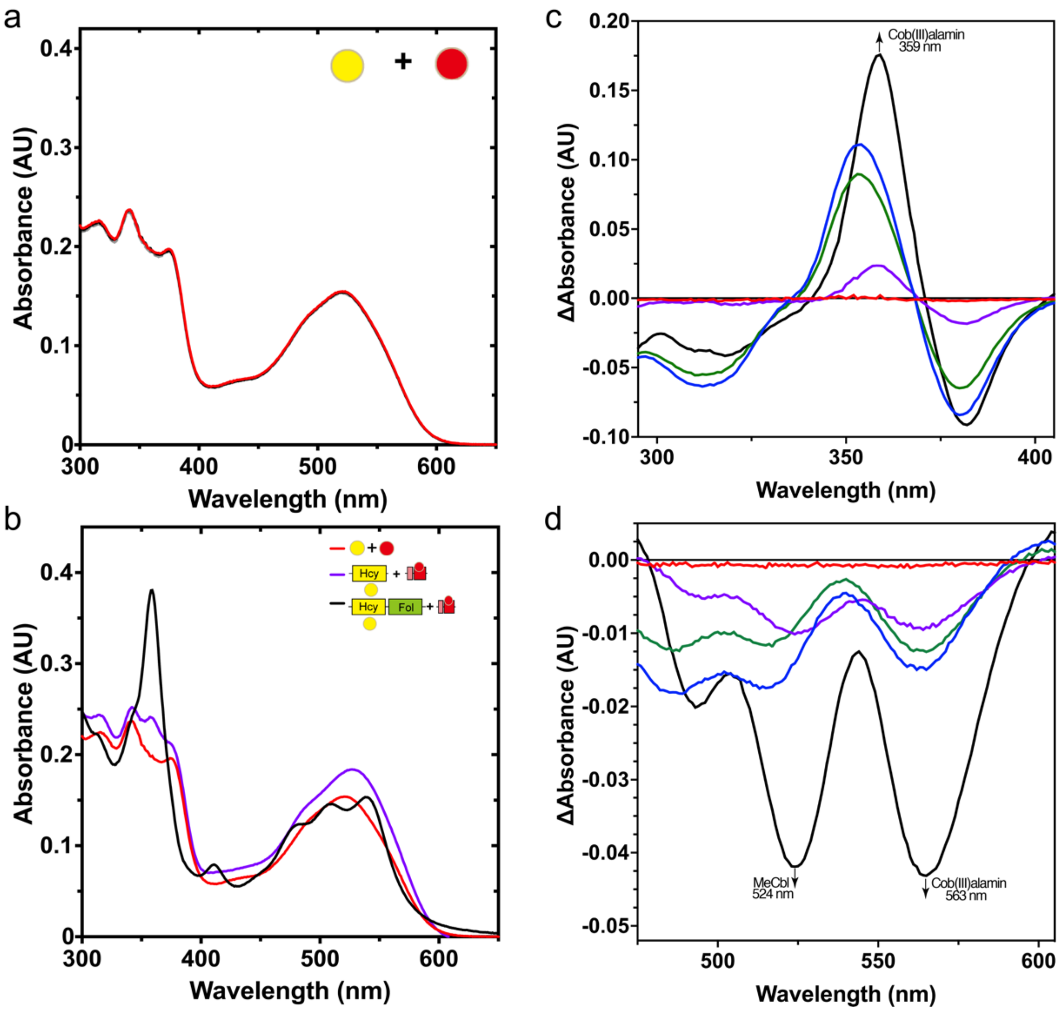


**Supplementary Figure 14. The role of the Fol domain in guiding catalytic transitions.** Effects of *t*MS domain fragments on MeCbl:homocysteine methyltransferase reaction. The time-dependent spectral changes of $CH_3$-Cbl in the presence of homocysteine were monitored at 50 ˚C for up to 10 minutes after adding homocysteine (final concentration of 100 μM) using 20 μM of free (non-protein bound) MeCbl incubated in the presence of 2 μM *t*MS constructs or MeCbl (23 μM) was bound to the isolated *t*MS$^{CapCob}$ domains first. **a** Free homocysteine and MeCbl did not react. **b** Final time points associated with free homocysteine + MeCbl, *t*MS bound Hcy (ΔN35Hcy) and MeCbl, and *t*MS bound Hcy (HcyFol) and MeCbl. Note that the presence of the Fol domain results in the associated consumption of MeCbl and production of aquacob(III)alamin. **c** Difference absorbance spectra of the MeCbl:homocysteine transferase assay, highlighting the importance of the Fol domain in the homocysteine methyltransferase reaction (black line) (300-400 nm) and **d** 450-600 nm. Data are of representative experiments, which have been repeated ≥2 times.

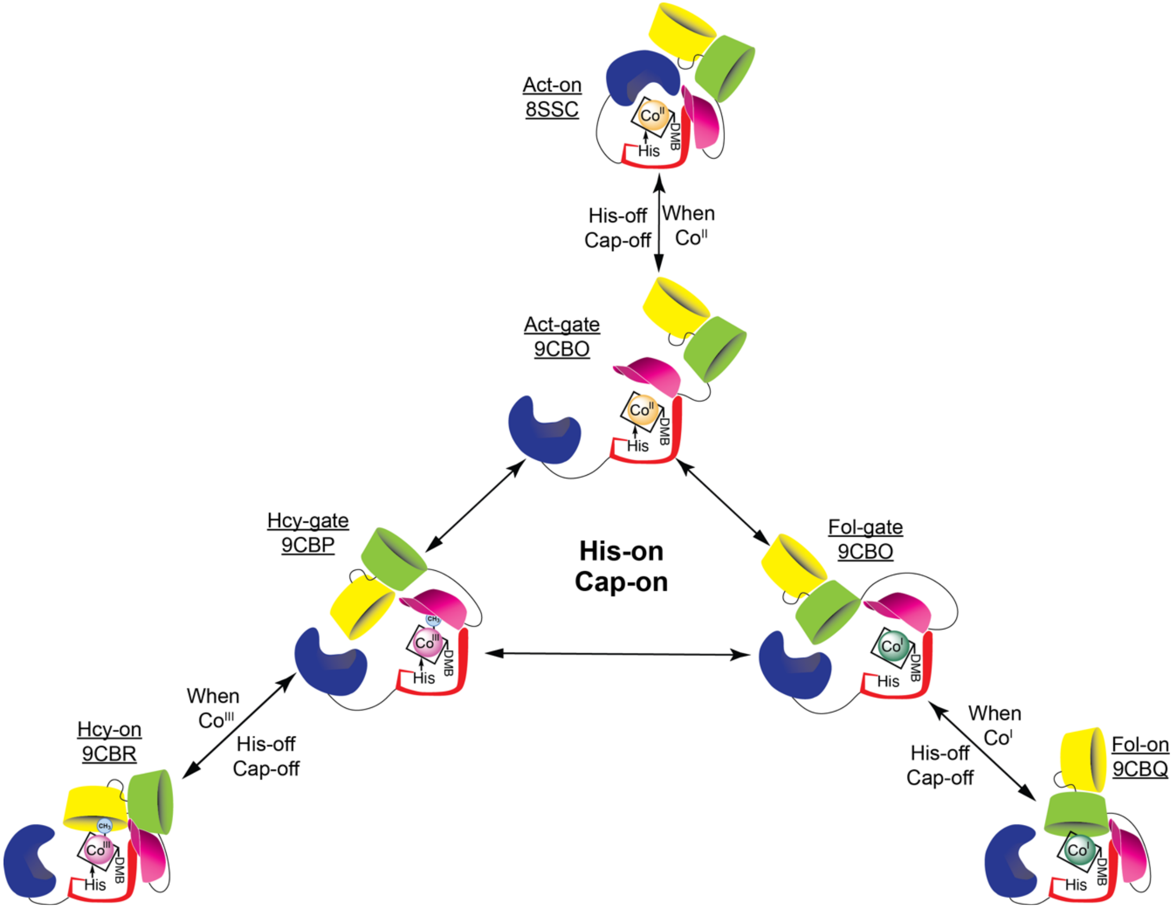


**Supplementary Figure 15. Methionine Synthase (MS) conformational ensemble as determined from this study.** The conformational ensemble of MS shown in cartoon form as determined from our *t*MS model. Yellow, green, pink, red, and blue are used to denote the Hcy, Fol, Cap, Cob, and Act domains, respectively. Corresponding state assignments and their respective PDB codes are shown; all structures save for Act-on (8SSC) are from this work. Cap-on states (pre-Hcy-on, 9CBP; pre-Fol-on, 9CBO; pre-Act-on, 9CBO) act as gateway conformations to their respective reactive conformations (Hcy-on, 9CBR; Fol-on; 9CBQ; Act-on, 8SSC). Note that the Cap-on gateway/pre-catalytic conformations are found to bind the cobalamin cofactor in the His-on form (methylcob(III)alamin) and can freely interchange, allowing for MS conformational 'resetting'. Uncapping corresponds with His-off ligation of the cobalamin cofactor, and each methylation is associated with a distinct cobalamin redox state (Co(III) for homocysteine methylation, Reaction I; Co(I) for methyltetrahydrofolate demethylation, Reaction II; Co(II) for reductive methylation via SAM/AdoMet, Reaction III).

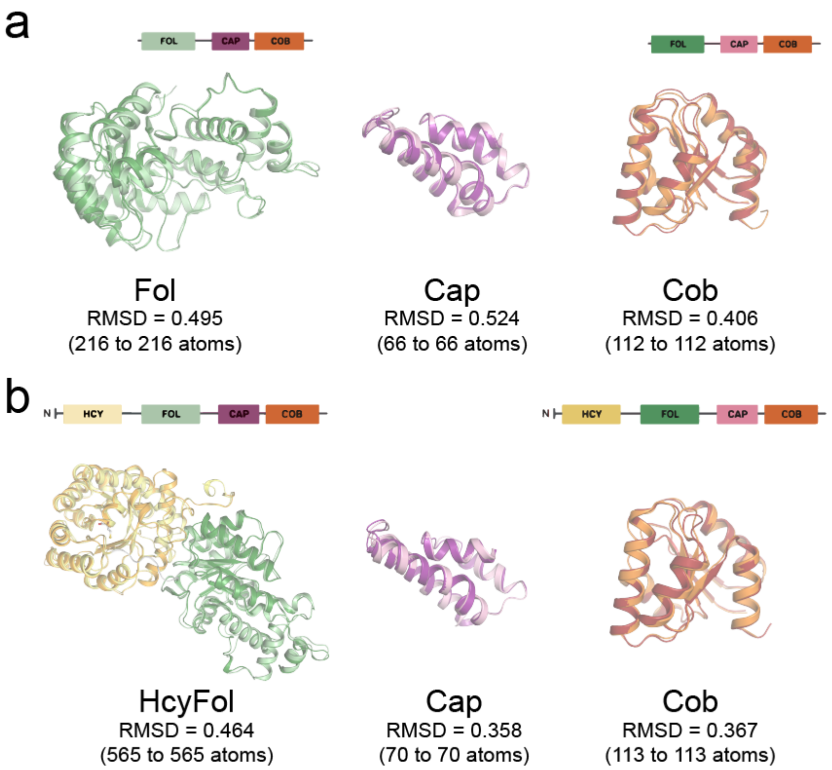


**Supplementary Figure 16. Structural alignment of the domains in the gateway and catalytic states of *t*MS**. **a** Comparison of individual domains from the pre-Fol-on (left) and Fol-on (right) structures are superimposed, with their respective root mean square deviation (RMSD) shown. **b** Comparison of individual domains from the pre-Hcy-on (left) and Hcy-on (right) structures are superimposed, with their respective root mean square deviation (RMSD) shown. Note that the individual domains are essentially identical, indicating that the structural differences lie primarily in the interdomain linkers and intradomain contacts.

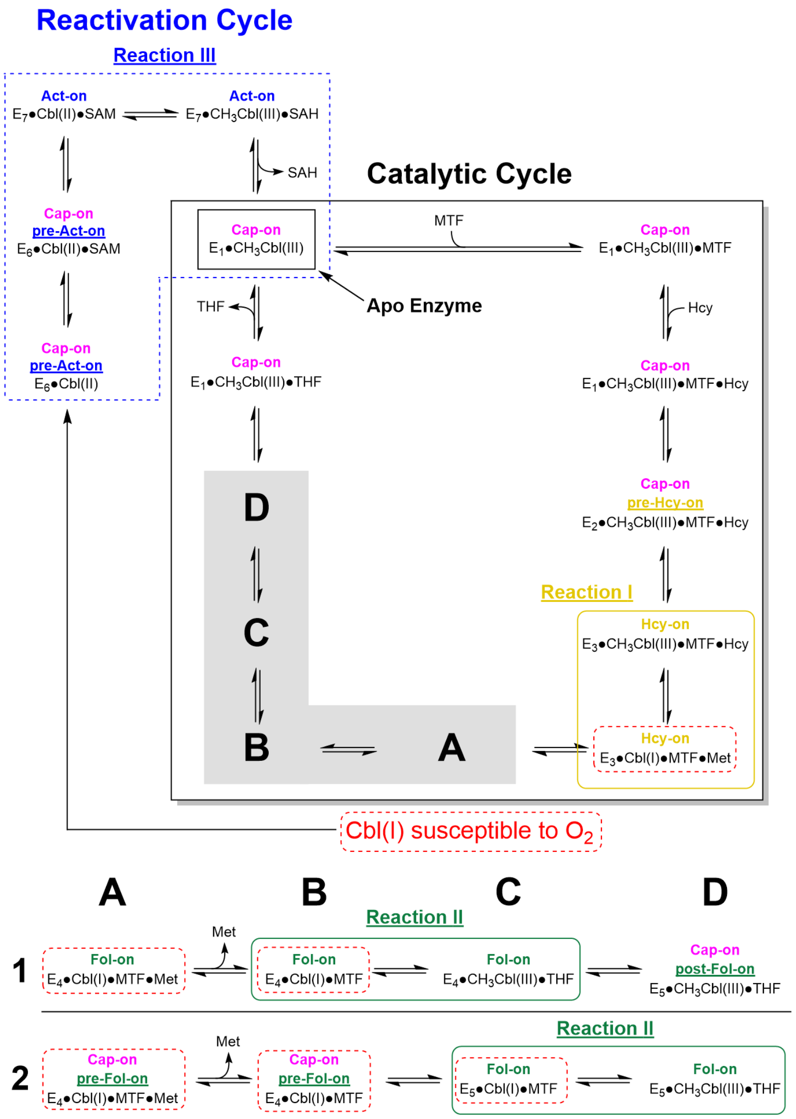


**Supplementary Figure 17. Unified models for MS catalysis provide a predictive framework for focused hypothesis testing.** States captured in this current study are highlighted in bold, while those predicted by this revised model are shown in dashed boxes. Note that while the gate states (pre-Fol-on, 9CBO; pre-Act-on, 9CBO) were captured in this study, their respective His-on ligation states were not. The same holds true for the catalytic Fol-on state (9CBQ), for which a His-off, Co(I) cofactor state is predicted in Model 2, but has not been observed. Model 1 precludes the need for Cap-on transitions during catalysis, while Model 2 assumes multiple Cap-on/off transitions occur during catalysis.

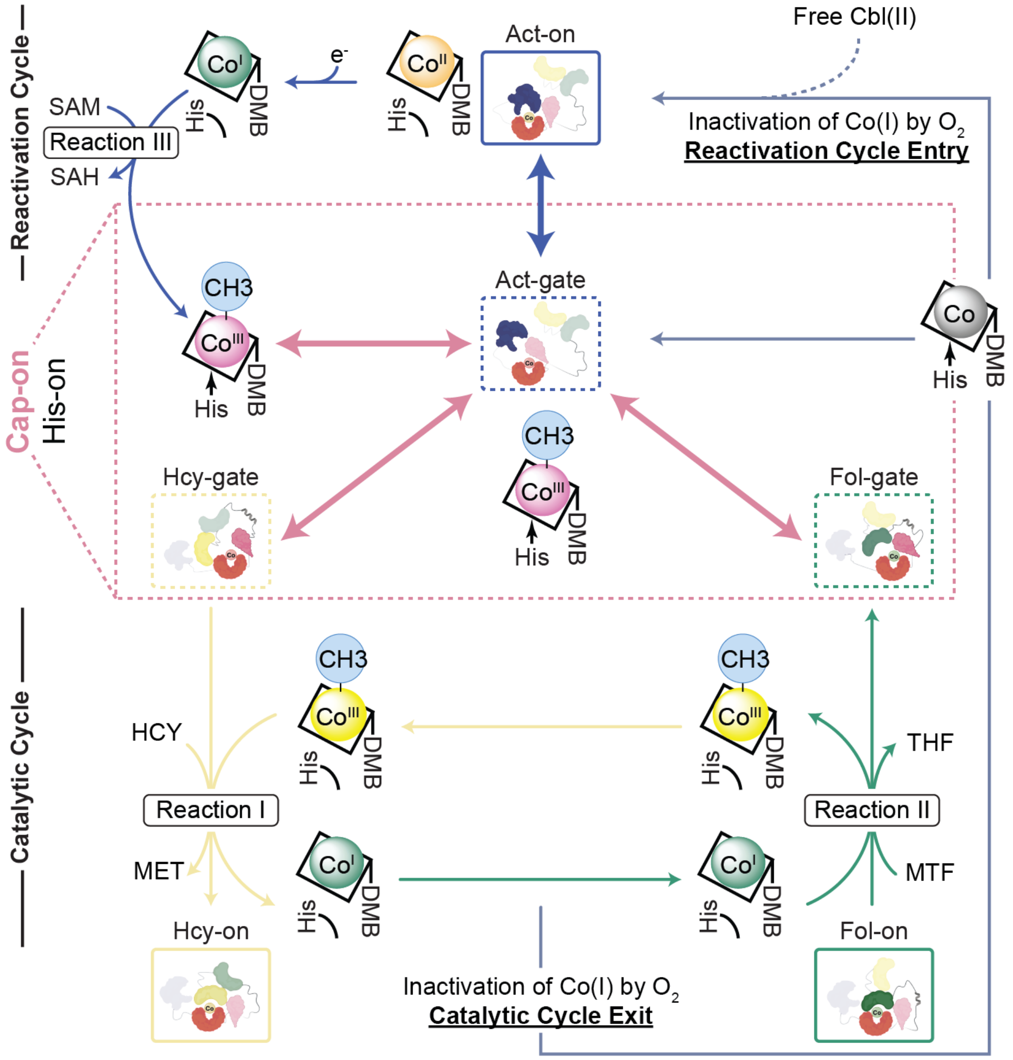


**Supplementary Figure 18. Expanded MS catalytic model including conformation ensemble and potential transitions.** This model is an expanded and more detailed map of the model shown in Fig. 7. Methionine synthase can cycle through Cap-on, His-on states that function as gateway conformations, obviating the need for uncapping. Linker II (Fol:Cap linker) allows for facile transition between these gateway states, what we term 'resetting'. In our gateway conformations, hexacoordinate, His-on methylcobalamin (Co(III)) highlight its role as a bridge between the primary catalytic and reactivation cycles. Depending on the cobalamin cofactor's status (redox state, coordinate number), the appropriate gateway state can grant entry to reactive conformations by uncapping and allowing for cobalamin flexibility by binding in the His-off state. Notably, the active methylating agent for methionine formation has been revised from hexa- to penta-coordinate His-off methylcob(III)alamin. Each methylation is associated with a distinct cobalamin redox state (Co(III) for homocysteine methylation, Reaction I; Co(I) for methyltetrahydrofolate demethylation, Reaction II; Co(II) for reductive methylation via SAM/AdoMet, Reaction III). The primary catalytic cycle of methionine synthase involves cycling between Co(III) and Co(I) states; Co(I) productions acts to lock MS to either the catalytic cycle (homocysteine methylation) or the reactivation cycle (Co(II) reductive alkylation) until methylcobalamin is regenerated or Co(I) undergoes oxidative inactivation.

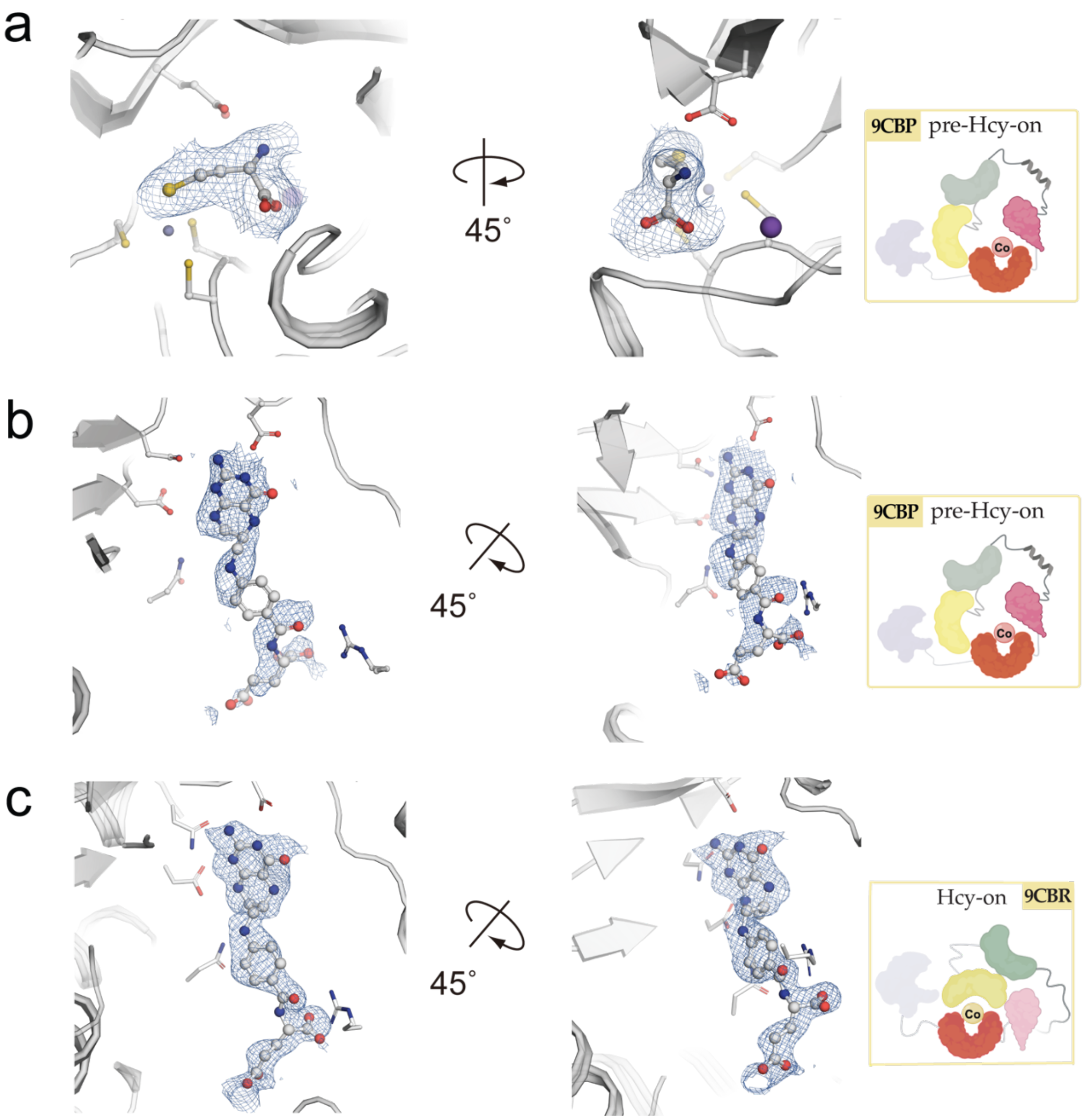


**Supplementary Figure 19. Substrate binding sites and electron density in *t*MS. a** Homocysteine binding site captured in 9CBP (pre-Hcy-on). **b** Tetrahydrofolate binding site captured in 9CBP (pre-Hcy-on). **c** Tetrahydrofolate binding site captured in 9CBR (Hcy-on). Their corresponding electron density ($2F_o$-$F_c$) contoured at 1.5 σ are shown in blue. Each substrate is shown in gray.

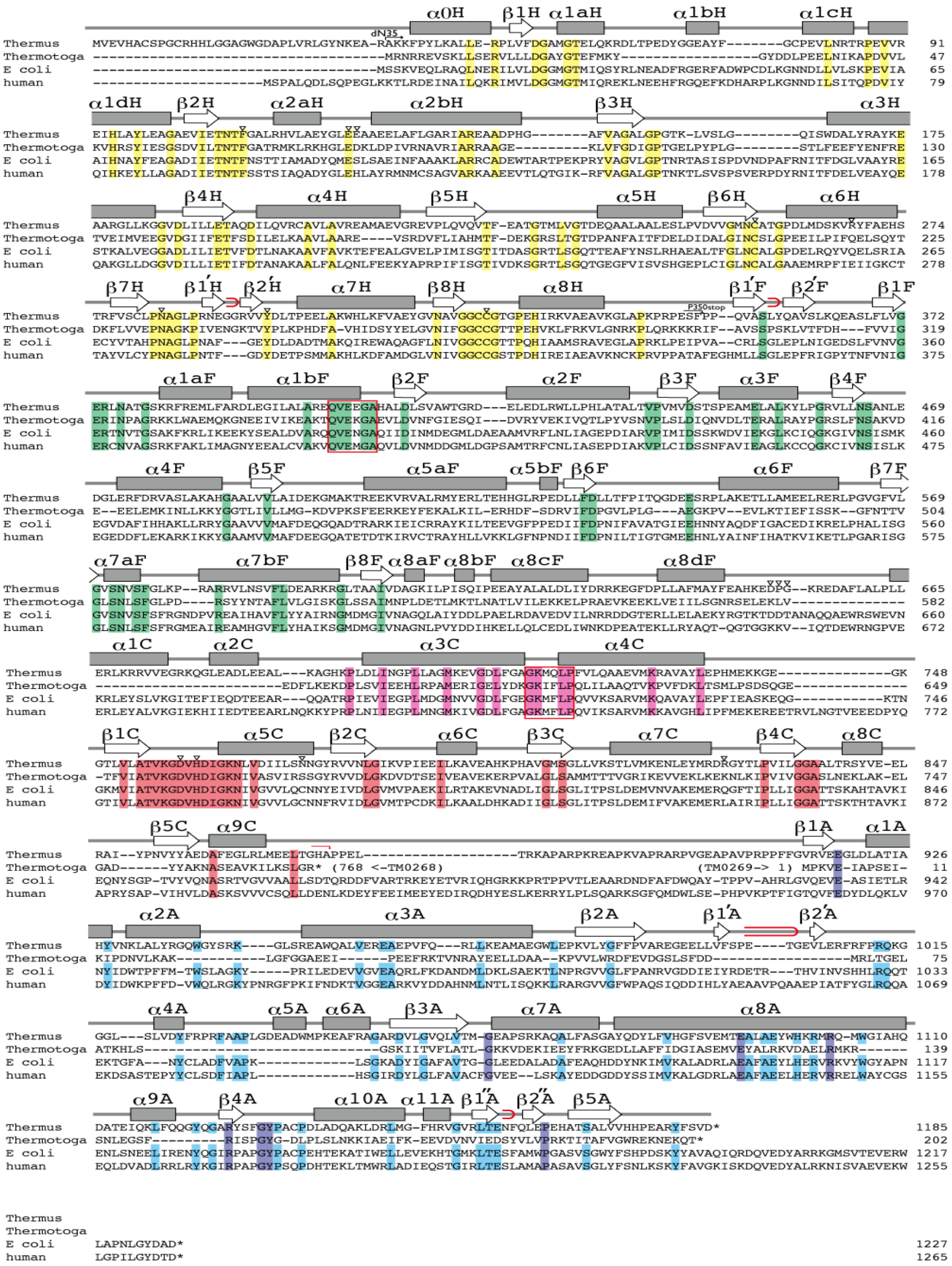

**Supplementary Figure 20. Methionine Synthase (MS) sequence alignment.** Schematic illustration of the secondary structure of *t*MS and alignments with the amino acid sequence of methionine synthase (MS) from *Thermus thermophilus* [Genbank accession number NC_00646], *Thermotoga maritima* [NC_000853], *Escherichia coli* [J04975], and *human* [U73338]. α-helices and β-sheets are shown in boxes and arrows, respectively. Red loops indicate β-hairpins. Conserved amino acid residues are highlighted by yellow, green, pink, red, and blue for the Hcy, Fol, Cap, Cob, and Act domains, respectively. In the activation (Act) domain, cyan is used to highlight conserved amino acid residues in *T. thermophilus*, *E. coli*, and *human* MS, while purple is used to highlight residues conserved among tetradomain MS.

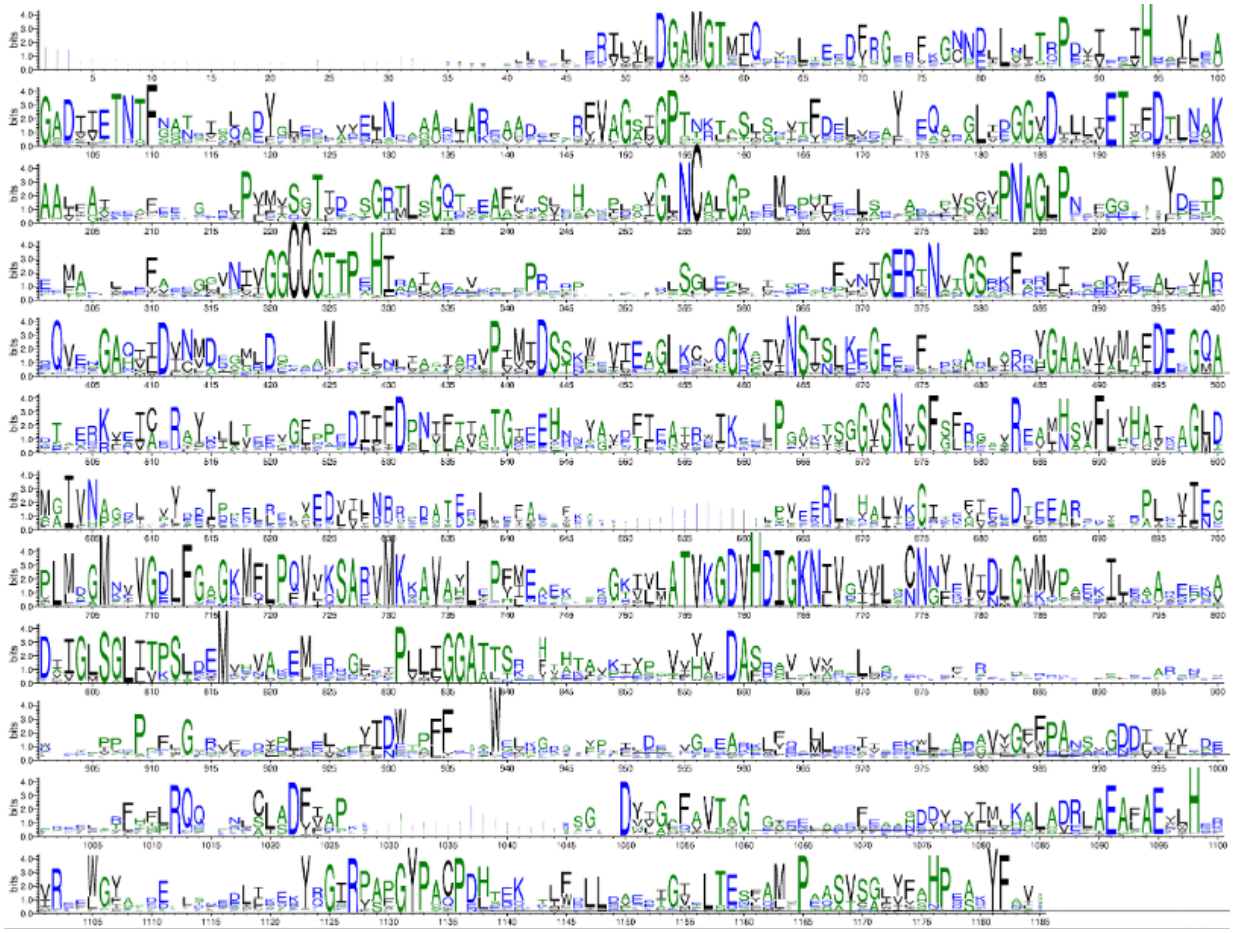

**Supplementary Figure 21. Weblogo representation of a multiple sequence alignment of Methionine Synthase (MS) (n=1983).** *t*MS (Uniprot ID Q5SKM5) was used as a query sequence. The domains are color-coded using the same scheme as in Supplementary Fig. 17 (yellow, green, pink, red, and blue for the Hcy, Fol, Cap, Cob, and Act domains, respectively). Residue numbering corresponds to that of the thermophilic enzyme. The plot was generated using DeepMSA2[1] and Weblogo3[2].

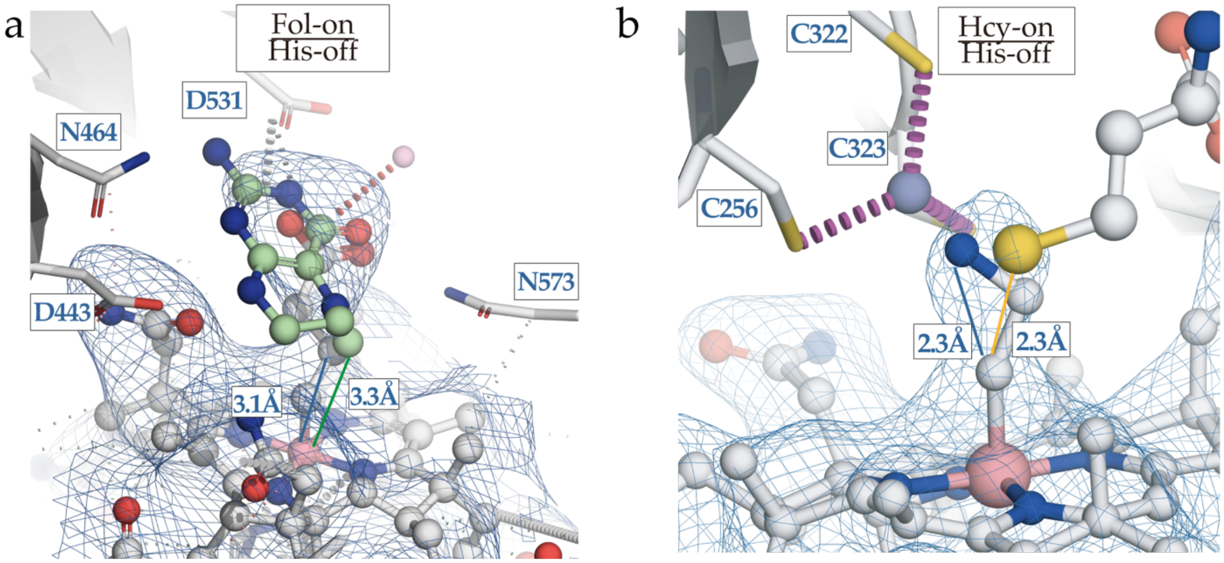


**Supplementary Figure 22. Non-native cobalamins used to capture catalytic conformations compete with substrate binding. a** The ternary complex required for methyltetrahydrofolate demethylation (MTF, Reaction II), Fol-on (9CBQ), was captured using cpCbl (γ-carboxypropyl upper axial ligand). Manual docking of MTF using structural alignment (5VOP, Fol domain containing MTF) shows that MTF and the upper axial ligand of cpCbl compete for the same residues in the Fol domain and occupy the same space above cobalamin. **b** The ternary complex required for homocysteine methylation (HCY, Reaction I), Hcy-on (9CBR), was captured using aeCbl (aminoethyl upper axial ligand). Manual docking of MTF using structural alignment (9CBP, Hcy domain containing HCY) shows HCY and the upper axial ligand of aeCbl compete for the same residues in the Hcy domain, and occupy the same space above cobalamin.

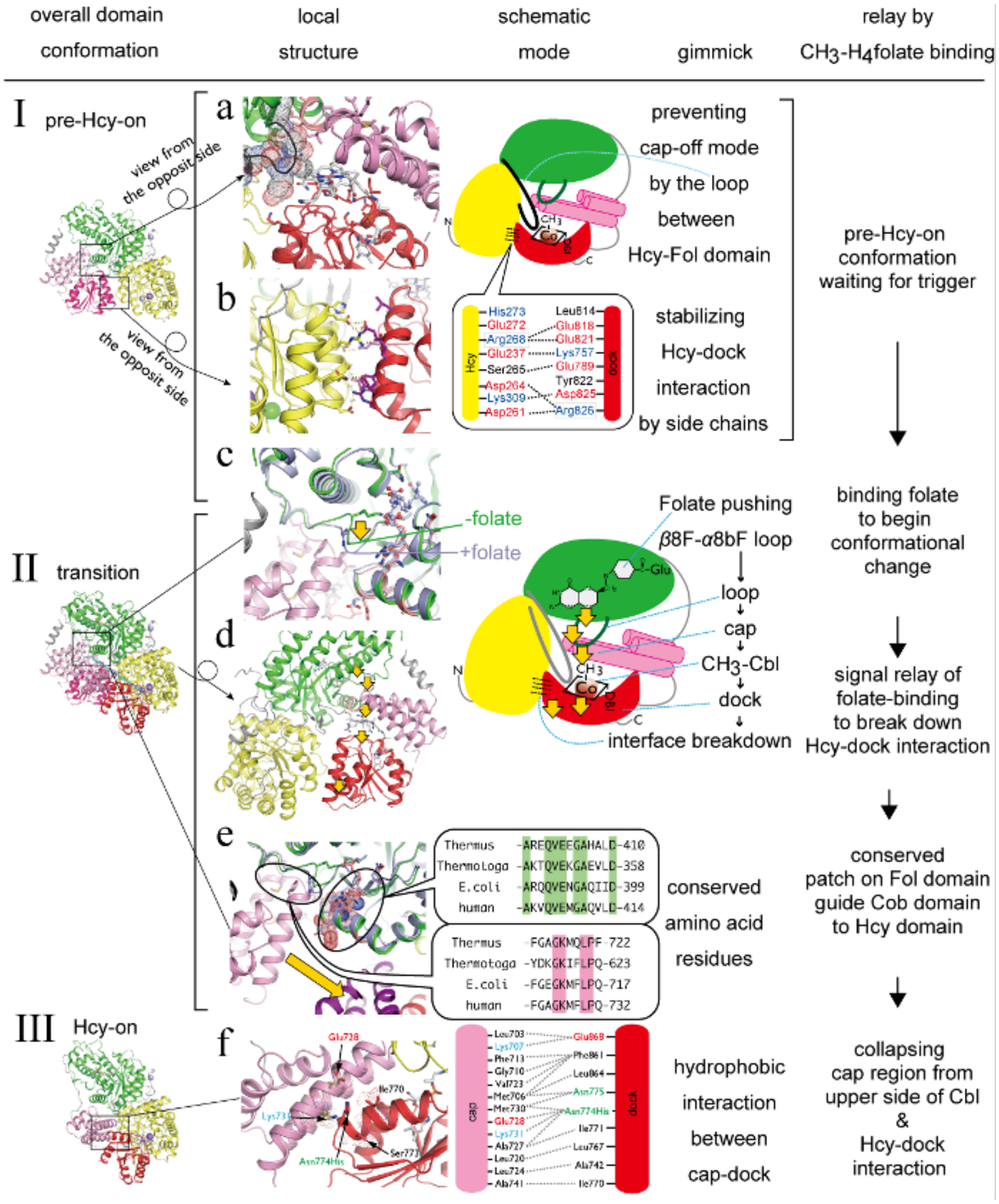


**Supplementary Figure 23. Structural elements that can potentially gate the transition between the pre-Hcy-on and Hcy-on states.** Conformational transitions from the pre-Hcy-on to Hcy-on mode and the role of methyltetrahydrofolate (MTF) binding to the Fol domain. Left column: the *t*MS$^{HcyFolCapCob}$ structures in the pre-Hcy-on (I) and Hcy-on (III) states are shown, color-coded according to domain (Hcy, yellow; Fol, green; Cap, pink; Cob, red). Their superimposition is shown in panel II. **a** The linker connecting the Hcy and Fol domains is drawn in the black loop for the backbone and in dot representation for side chains. **b** The Hcy and Cob domain interaction in the pre-Hcy-on conformation is schematically drawn, based on PDBSum analysis. **c** The positional differences of the loop between β8F and α8bF in the Fol domain are illustrated. The loop is repositioned in the presence (pale blue) and absence (green) of MTF. The orange arrow indicates the possible loop motion upon binding of MTF to the Fol domain. **d** The cascade triggered by MTF binding, leading to the collapsing interaction between the Hcy and Cob domains, is illustrated with five orange arrows. **e** Amino acid sequence alignment of MS from Thermus (*Thermus thermophilus*), Thermotoga (*Thermotoga maritima*), E. coli (*Escherichia coli*), and human (*Homo sapiens*) is presented. Conserved amino acid residues in the Fol domain are highlighted in green, while those in the Cob domain (cap region) are marked in pink. **f** The Cap-Cob interaction in the Hcy-on (Cap-off) conformation is illustrated schematically, based on PDBSum analysis[3].

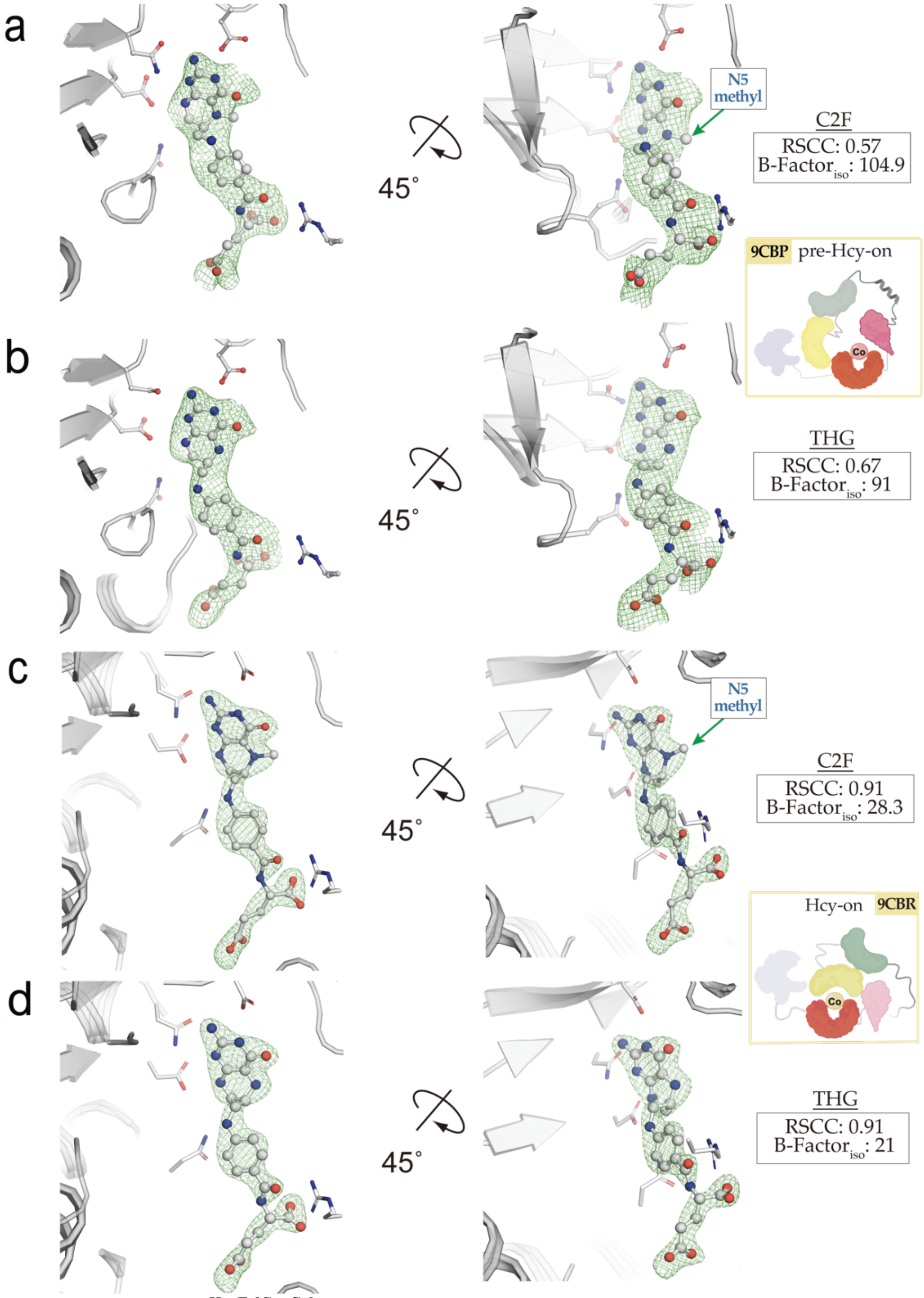


**Supplementary Figure 24. *t*MS$^{HcyFolCapCob}$ copurifies with tetrahydrofolate in both the pre-Hcy-on and Hcy-on states. a** Methyltetrahydrofolate (ligand code C2F) with its corresponding electron density from a polder omit map ($F_o$-$F_c$) contoured at 3 σ are shown in green for 9CBP (pre-Hcy-on). **b** Tetrahydrofolate (ligand code THG) with its corresponding electron density from a polder omit map ($F_o$-$F_c$) contoured at 3 σ are shown in green for 9CBP (pre-Hcy-on). **c** Methyltetrahydrofolate (ligand code C2F) with its corresponding electron density from a polder omit map ($F_o$-$F_c$) contoured at 3 σ are shown in green for 9CBR (Hcy-on). **d** Tetrahydrofolate (ligand code THG) with its corresponding electron density from a polder omit map ($F_o$-$F_c$) contoured at 3 σ are shown in green for 9CBR (Hcy-on). The ligand fit was assessed using the real-space correlation coefficient (RSCC) and isotropic B-factor (B-Factor$_{iso}$); a corresponding increase in the RSCC and decrease in the B-Factor$_{iso}$ was observed when THG was fit into the electron density versus C2F. The ligand assignment was thus ascribed to tetrahydrofolate (THG) for both 9CBP and 9CBR. Note that the observed occupancy for THG was 0.88 for 9CBP, with full occupancy observed in 9CBR.

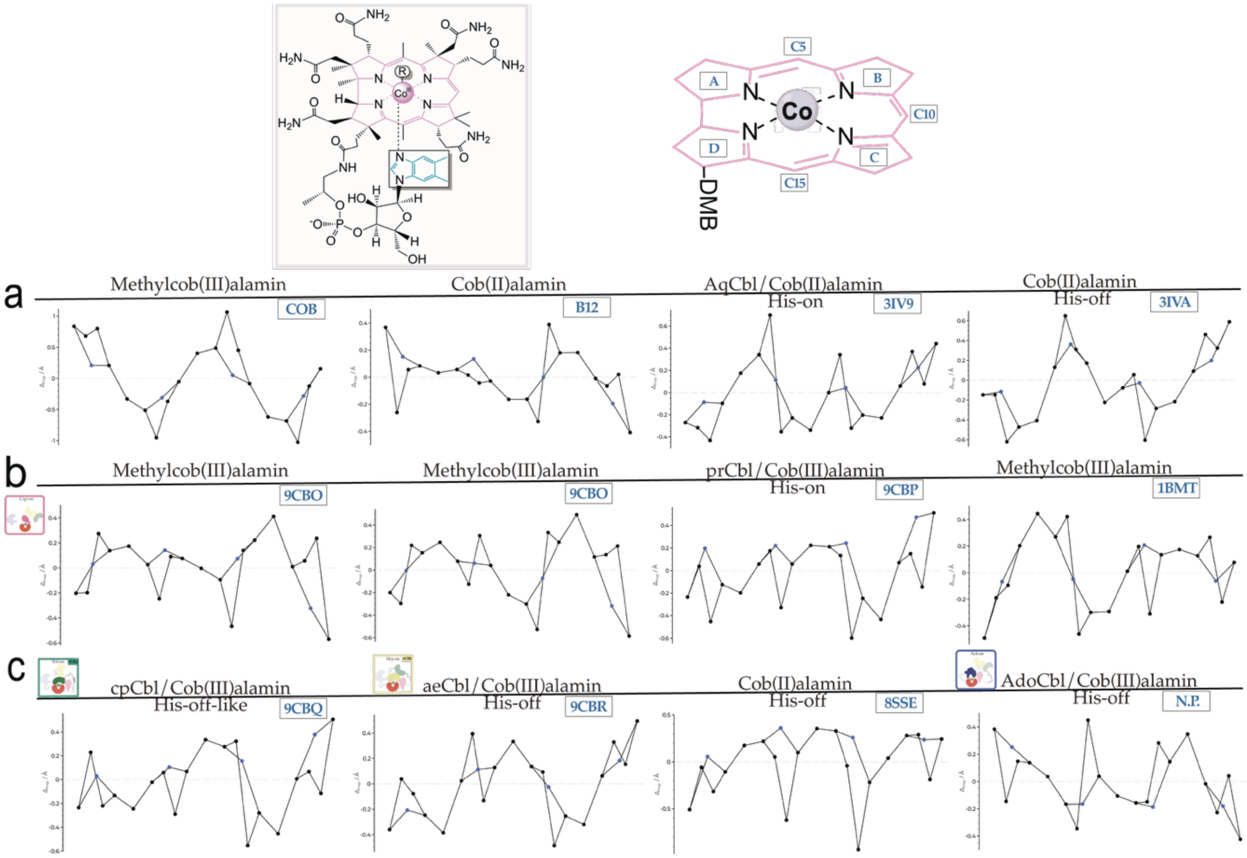


**Supplementary Figure 25. Corrin ring distortion analysis reveal changes associated with redox and ligation status. a** Corrin ring distortion analysis of reference cobalamin ligands and deposited structures (COB, B12, 3IV9, 3IVA). **b** Corrin ring distortion analysis of Cap-on structures discussed in this paper (9CBO, pre-Fol-on, Chain A, left; 9CBO, 'Resting', Chain D, $2^{nd}$ from left; 9CBP, pre-Hcy-on, $2^{nd}$ from right; 1BMT, right). **b** Corrin ring distortion analysis of Cap-off structures discussed in this paper (9CBQ, Fol-on, left; 9CBR, Hcy-on, $2^{nd}$ from left; 8SSE, Chain B, Act-on, $2^{nd}$ from right; unpublished/not published Act-on, His-off, right). Normal-coordinate structure decomposition (NSD) analysis was conducted using PorphyStruct (delta out-of-plane ($\Delta oop$) distortions in Å)[4].

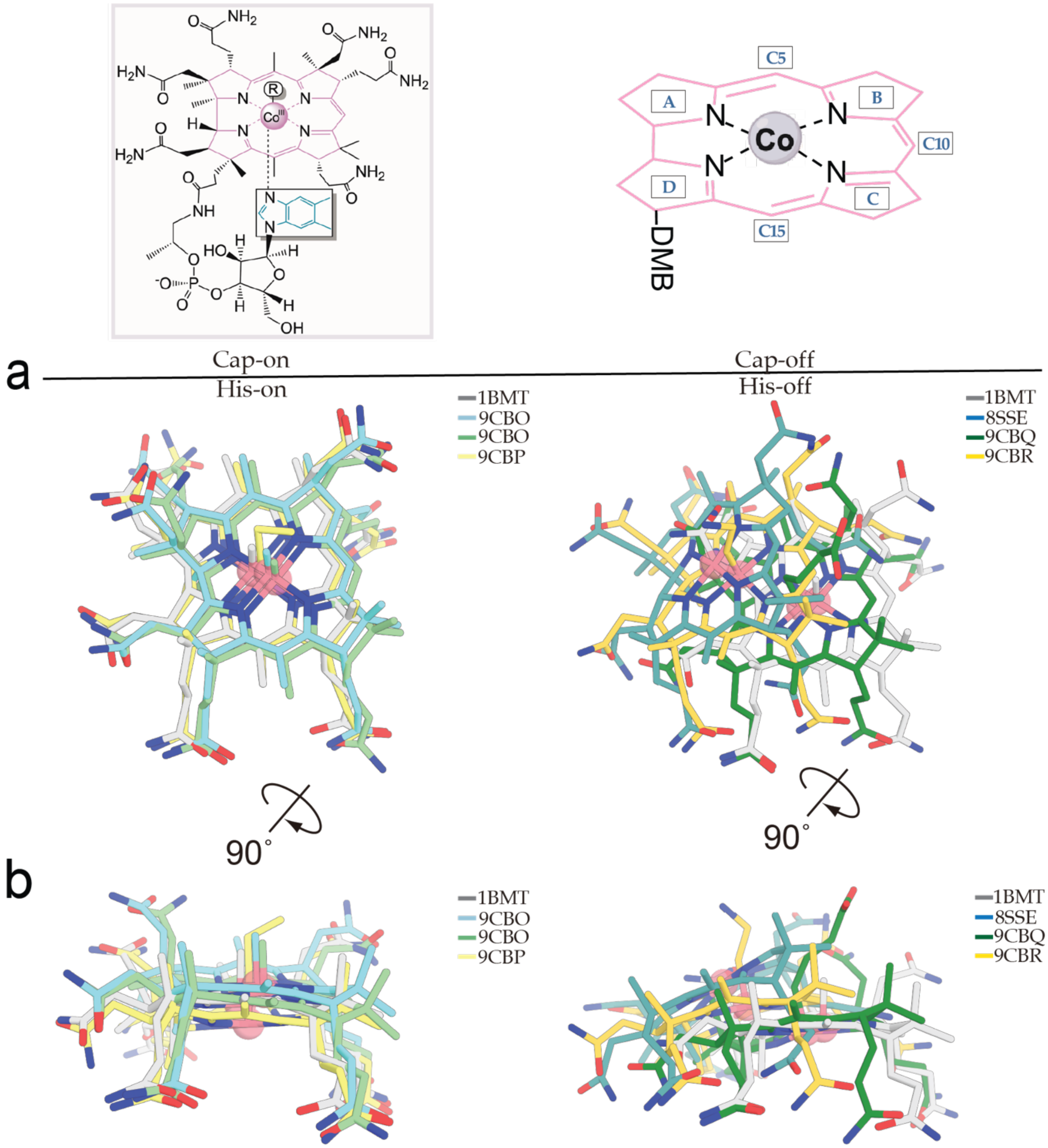


**Supplementary Figure 26. Corrin ring overlay of captured states reveals changes associated with redox and ligation status. a** Corrin ring overlay of cobalamin ligands from their deposited structures, using 1BMT (light grey, Chain A) and 8SSE (teal, Chain B) for reference. Note, the side view shown in **b** demonstrates the large effect His-off ligation has on the lateral displacement of the corrin ring of cobalamin. 9CBR shows a drastic lateral movement akin to that of His-off cob(II)alamin (8SSE). There are subtle corrin ring distortions even in the His-on, Cap-on states, with 9CBP showing downwards doming.

### Supplementary Table 1. X-Ray Data Collection and Refinement Statistics

| | $t\mathrm{MS}^{\mathrm{FolCapCobD759A•MeCbl}}$ | $t\mathrm{MS}^{\mathrm{HcyFolCapCobmut1•prCbl}}$ | $t\mathrm{MS}^{\mathrm{FolCapCobD762G•cpCbl}}$ | $t\mathrm{MS}^{\mathrm{HcyFolCapCobmut2•aeCbl}}$ |
|---|---|---|---|---|
| **Data collection** | | | | |
| Beamline | APS, GMCA 23-IDB | APS, LS-CAT 21-IDD | APS, GMCA 23-IDB | APS, GMCA 23-IDD |
| Wavelength (Å) | 1.0332 | 1.2713 | 1.0332 | 1.0332 |
| Temperature (K) | 100 | 100 | 100 | 100 |
| Resolution (Å) | 325.55-3.34 (3.40-3.34)* | 83.82-2.45 (2.51-2.45)* | 52.77-2.87 (2.95-2.87)* | 120.79-2.38 (2.47-2.38)* |
| Space group | $P4_12_12$ | $P2_12_12_1$ | $P6_122$ | $P2_12_12_1$ |
| Cell dimensions | | | | |
| *a*, *b*, *c* (Å) | 187.77, 187.77, 325.55 | 69.95, 83.82, 163.72 | 95.16, 95.16, 274.34 | 70.81, 115.10, 120.80 |
| α, β, γ (°) | 90, 90, 90 | 90, 90, 90 | 90, 90, 120 | 90, 90, 90 |
| Observed reflections | 702,906 (36,503) | 228,743 (21,780) | 236,810 (35,228) | 222,476 (19,913) |
| Unique reflections | 85,017 (4,350) | 35,755 (3,794) | 17,680 (2,476) | 39,748 (3,707) |
| $R_{\mathrm{meas}}$ (%) | 25.6 (233.3) | 13.0 (67.90) | 12.6 (235.6) | 60.5 (460.4) |
| $R_{\mathrm{merge}}$ (%) | 24.0 (219.4) | 12.0 (61.70) | 12.1 (227.2) | 54.9 (416.6) |
| <I/σ> | 6.7 (1.2) | 9.4 (2.5) | 11.7 (0.7) | 3.8 (0.6) |
| CC(1/2) | 0.986 (0.428) | 0.996 (0.880) | 0.999 (0.893) | 0.908 (0.221) |
| Multiplicity | 8.3 (8.4) | 6.4 (5.7) | 13.4 (14.2) | 5.6 (5.4) |
| Completeness (%) | 99.9 (98.1) | 98.2 (91.0) | 99.8 (97.6) | 98.8 (85.12) |
| Wilson *B*-factor ($Å^2$) | 100.1 | 38.7 | 106.9 | 32.51 |
| **Refinement** | | | | |
| Resolution (Å) | 163.18 - 3.34 | 81.99 - 2.44 | 52.77 - 2.87 | 83.33 - 2.38 |
| No. reflections | 80,679 (4,225)‡ | 33,900 (2,296)‡ | 16,703 (1,161)‡ | 37,865 (2,403)‡ |
| $R_{\mathrm{work}}/R_{\mathrm{free}}$ (%) | 17.3/21.6 | 16.1/23.7 | 21.3/27.2 | 19.5/21.7 |
| No. of non-H atoms | | | | |
| Protein | 23,836 | 6,624 | 3,869 | 6,646 |
| Water | 110 | 200 | 29 | 248 |
| Ligand | 588 | 147 | 122 | 130 |
| B-factors ($Å^2$) | | | | |
| Protein | 120.10 | 68.87 | 130.16 | 36.59 |
| Water | 62.95 | 63.39 | 94.66 | 20.52 |
| Ligand | 102.96 | 70.05 | 119.14 | 24.16 |
| R.m.s. deviations | | | | |
| Bond lengths (Å) | 0.015 | 0.017 | 0.016 | 0.014 |
| Bond angles (°) | 2.31 | 2.41 | 2.72 | 2.21 |
| Ramachandran Plot | | | | |
| Favored/allowed/outliers | 97.27/2.66/0.07 | 97.3/2.5/0.2 | 95.51/3.88/0.61 | 96.18/3.70/0.12 |
| MolProbity Score | 1.44 (100th percentile) | 1.72 (98th percentile) | 1.54 (100th percentile) | 1.04 (100th percentile) |
| PDB | 9CBO | 9CBP | 9CBQ | 9CBR |

* Highest-resolution shell is shown in parentheses.

‡ Number of reflections used for cross-validation

**Supplementary Table 2. Bacterial and insect strains, plasmids, and synthetic oligonucleotides used in this study**

| Strains, Plasmids, Primers | Relevant characteristics | Ref. or Sources |
|---|---|---|
| ***E. coli strains*** | | |
| *XL1-Blue* | Routine cloning strain, tetracycline resistance | Stratagene |
| BL21star(DE3) | Widely used T7 expression system, no antibiotic resistance | Invitrogen |
| ***Plasmids*** | | |
| pET11a(*t*MS$^{wt}$) | wild-type *t*MS in pET11a vector, containing the *T. thermophilus* gene MetH, Amp$^{R}$ | Riken |
| pMCSG7 – *t*MS Clones | | |
| wt | wild-type *t*MS in pMCSG7 vector, encoding an N-terminal His-tag with TEV cleavage site, Amp$^{R}$ | Mendoza *et al.* 2023 |
| ΔN35Hcy | *t*MS N-terminal 35aa truncation (ΔN35), Hcy domain (Ala36-Phe349) | This Work |
| HcyFol | *t*MS N-terminal half, Hcy and Fol domains (Met1-His648) | This Work |
| CapCob | *t*MS Cap and Cob domains (Leu660-Ala874) | This Work |
| FolCapCob | *t*MS Fol, Cap, and Cob domains (tridomain) (Gln364-Ala874) | This Work |
| FolCapCob$^{D759A}$ | *t*MS tridomain Asp759Ala mutant | This Work |
| FolCapCob$^{D762G}$ | *t*MS tridomain Asp762Gly mutant | This Work |
| HcyFolCapCob | *t*MS Hcy, Fol, Cap, and Cob domains (tetradomain) (Met1-Ala874) | This Work |
| HcyFolCapCob$^{F110A,E123A,E124A,Y296A,D651A,P652A,G653A}$ | tetradomain, Phe110Ala, Glu123Ala, Glu124Ala, Tyr296Ala, Asp651Ala, Gly653Ala mutant (mutant 1) | This Work |
| HcyFolCapCob$^{F110A,E123A,E124A,R268A,D651A,P652A,G653A,D759A,N774H,R826A}$ | tetradomain, Phe110Ala, Glu123Ala, Glu124Ala, Arg268Ala, Asp651Ala, Pro652Ala, Gly653Ala, Asp759Ala, Asn774His, Arg826Ala mutant (mutant 2) | This Work |

***Oligonucleotides***

| Primer Names | Sequences | Descriptions |
|---|---|---|
| pMCSG7(*t*MS$^{wt}$) | | |
| *t*MS_LIC-f | 5' - ACTTCCAATCCAATGCCATGGTGGAGGTCCACGCCTG - 3' | LIC, wt, vector |
| *t*MS_LIC-r | 5' - TTATCCACTTCCAATGCTAGTCCACGCTGAAGTAGCG - 3' | pET(*t*MS$^{wt}$) template |
| pMCSG7(*t*MS$^{\Delta N35Hcy}$) | | |
| ΔN35Hcy-f | 5' - TACTTCCAATCCAATGCGAAGAAGTTTCCCTACCTCAAG - 3' | LIC, *t*MS$^{HcyFol}$ template |
| ΔN35Hcy-r | 5' - TTATCCACTTCCAATGCTAGAAGCTTTCAGGCCTTGG - 3' | |
| pMCSG7(*t*MS$^{HcyFol}$) | | |
| HcyFol-f | 5' - TACTTCCAATCCAATGCCATGGTGGAGGTCCACGCCTG - 3' | LIC, pET(*t*MS$^{wt}$) template |
| HcyFol-r | 5' - TTATCCACTTCCAATGCTAGTGGGCCTCAAAGTAGGC - 3' | |
| pMCSG7(*t*MS$^{CapCob}$) | | |
| CapCob-f | 5' - TACTTCCAATCCAATGCCCTGGCCCTTCCCCTCCTGGAG - 3' | LIC, *t*MS$^{FolCapCob}$ template |
| CapCob-r | 5' - TTATCCACTTCCAATGCTAGGCGTGGCCCGTGAGCTC - 3' | |
| pMCSG7(*t*MS$^{FolCapCob}$) | | |
| FolCapCob-f | 5' - TACTTCCAATCCAATGCGCAGGAGGCGAGCCTTTTCCTC - 3' | LIC, pET(*t*MS$^{wt}$) template |
| FolCapCob-r | 5' - TTATCCACTTCCAATGCTAGGCGTGGCCCGTGAGCTC - 3' | |
| D759A-f | 5' - CGTCAAGGGGGCCGTGCACGACATC - 3' | Site-directed mutagenesis |
| D759A-r | 5' - GATGTCGTGCACGGCCCCCTTGACG - 3' | |
| D762G-f | 5' - GGGACGTGCACGGCATCGGCAAGAA - 3' | Site-directed mutagenesis |
| D759G-r | 5' - TTCTTGCCGATGCCGTGCACGTCCC - 3' | |
| pMCSG7(*t*MS$^{HcyFolCapCob}$) | | |
| HcyFolCapCob-f | 5' - TACTTCCAATCCAATGCCATGGTGGAGGTCCACGCCTG - 3' | LIC, pET(*t*MS$^{wt}$) template |
| HcyFolCapCob-r | 5' - TTATCCACTTCCAATGCTAGGCGTGGCCCGTGAGCTC - 3' | |
| D651A, P652A, G653A-f | 5' - GAGGCCCACAAGGAGGCCGCGGCGAAGAGGGAGGACGCC - 3' | Site-directed mutagenesis |
| D651A, P652A, G653A-r | 5' - GGCGTCCTCCCTCTTCGCCGCGGCCTCCTTGTGGGCCTC - 3' | DPG2A |
| E123A, E124A-f | 5' - CGCCGAGTACGGCCTGGCGGCGGCGGCGGAGGAGCTCGCC - 3' | Site-directed mutagenesis |
| E123A, E124A-r | 5' - GGCGAGCTCCTCCGCCGCCGCCGCCAGGCCGTACTCGGCG - 3' | dEE |

| | | |
|---|---|---|
| F110A-f | 5' - CATAGAGACCAACACCGCCGGGGCCCTGCGCC - 3' | Site-directed mutagenesis |
| F110A-r | 5' - GGCGCAGGGCCCCGGCGGTGTTGGTCTCTATG - 3' | |
| Y296A-f | 5' - GAGGGTGGTCGCCGACCTCACCCCC - 3' | Site-directed mutagenesis |
| Y296A-r | 5' - GGGGGTGAGGTCGGCGACCACCCTC - 3' | |
| R268A-f | 5' - CATGGACAGCAAGGTGGCCTACTTCGCCGAGCAC - 3' | Site-directed mutagenesis |
| R268A-r | 5' - GTGCTCGGCGAAGTAGGCCACCTTGCTGTCCATG - 3' | dRR (1/2) |
| R826A-f | 5' - GAGTACATGCGGGATGCGGGCTACACCCTCCCCG - 3' | Site-directed mutagenesis |
| R826A-r | 5' - CGGGGAGGGTGTAGCCCGCATCCCGCATGTACTC - 3' | dRR(2/2) |
| D759A-f | 5' - CGTCAAGGGGGCCGTGCACGACATC - 3' | Site-directed mutagenesis |
| D759A-r | 5' - GATGTCGTGCACGGCCCCCTTGACG - 3' | |
| N774H-f | 5' - GACATCATCCTCAGCCACAACGGCTACCGGGTGG - 3' | Site-directed mutagenesis |
| N774H-r | 5' - CCACCCGGTAGCCGTTGTGGCTGAGGATGATGTC - 3' | |
